# “Roles for ELMOD2 and Rootletin in Ciliogenesis”

**DOI:** 10.1101/2021.01.04.425267

**Authors:** Rachel E. Turn, Joshua Linnert, Eduardo D. Gigante, Uwe Wolfrum, Tamara Caspary, Richard A. Kahn

## Abstract

ELMOD2 is a GTPase activating protein (GAP) with uniquely broad specificity for ARF family GTPases. We previously showed that it acts with ARL2 in mitochondrial fusion and microtubule stability and with ARF6 during cytokinesis. Mouse embryonic fibroblasts deleted for ELMOD2 also displayed changes in cilia related processes including increased ciliation, multiciliation, ciliary morphology, ciliary signaling, centrin accumulation inside cilia, and loss of rootlets at centrosomes with loss of centrosome cohesion. Increasing ARL2 activity or overexpressing Rootletin reversed these defects, revealing close functional links between the three proteins. This was further supported by the findings that deletion of Rootletin yielded similar phenotypes, which were rescued upon increasing ARL2 activity but not ELMOD2 overexpression. Thus, we propose that ARL2, ELMOD2, and Rootletin all act in a common pathway that suppresses spurious ciliation and maintains centrosome cohesion. Screening a number of markers of steps in the ciliation pathway support a model in which ELMOD2, Rootletin, and ARL2 act downstream of TTBK2 and upstream of CP110 to prevent spurious release of CP110 and to regulate ciliary vesicle docking. These data thus provide evidence supporting roles for ELMOD2, Rootletin, and ARL2 in the regulation of ciliary licensing.

## Introduction

Members of the ARF (ADP-ribosylation factor) family of regulatory GTPases, as well as their downstream effectors and GTPase activating proteins (GAPs) that regulate their activities, drive an incredibly diverse array of cellular functions (Casalou et al., 2020; Fisher et al., 2020; Francis et al., 2016; Sztul et al., 2019). Consisting of 6 ARFs, 22 ARLs (ARF-like proteins), and 2 SARs in mammals, the ARF family is ancient, with multiple members traced back to the last eukaryotic common ancestor (Li et al., 2004). One critical feature of these proteins is that they localize to multiple cellular compartments and can perform discrete functions at each site, making them both critical to healthy cell function and technically challenging to dissect functionality. Because individual family members repeatedly have been found capable of regulating multiple processes at distinct cellular sites, they have been proposed as key players in inter-pathway communication or higher order signaling (Francis et al., 2016). Although the canonical model for regulatory GTPase actions is that a guanine nucleotide exchange factor (GEF) activates the GTPase (by promoting release of GDP and binding of GTP) and that GAPs terminate the activated state (by promoting hydrolysis of the bound GTP), ARF GAPs consistently have been found to possess both GAP and effector activities (East and Kahn, 2011; Sztul et al., 2019; Zhang et al., 2003; Zhang et al., 1998). Because previous studies have identified ARF GAPs as downstream mediators/effectors of the GTPases they bind, they, too, act in multiple pathways. This is particularly true for the ELMOD family of ARF GAPs, at least in part due to their uniquely broad specificity towards both ARFs and ARLs (Ivanova et al., 2014). This is in contrast to the much larger family of 24 known ARF GAPs (including ACAPs, ASAPs, ARAPs) that bind and promote GTP hydrolysis only on ARFs, but not ARLs (Cuthbert et al., 2008; Sztul et al., 2019; Vitali et al., 2019).

Like the ARF family, the ELMODs are also ancient and were present in the last eukaryotic common ancestor (East et al., 2012). The three mammalian ELMOD family members, ELMOD1-3, share a single (ELMO) domain that gives the protein its GAP activity. This domain contains a predicted “arginine finger” that is directly involved in GTP hydrolysis (Ahmadian et al., 1997; East et al., 2012). Mutation of this single arginine is sufficient to eliminate in vitro GAP activity (Ivanova et al., 2014). ELMODs are also implicated in a number of pathologies, including deafness in mammals (ELMOD1, ELMOD3 (Jaworek et al., 2013; Johnson et al., 2012; Lahbib et al., 2018; Li et al., 2019; Li et al., 2018)), intellectual disability (ELMOD1, ELMOD3 (Miryounesi et al., 2019)), idiopathic pulmonary fibrosis, and antiviral response (ELMOD2 (Hodgson et al., 2006; Pulkkinen et al., 2010)). The mechanisms by which mutations or disruption of these proteins causes disease are unclear. Because of the apparent importance of ELMODs to cell regulation and their predicted impact on our understanding of multiple disease states, we have undertaken a broad analysis of cellular roles for ELMODs using a number of technical approaches.

ELMOD2 is a ~37 kDa protein that was first purified as an ARL2 GAP (Bowzard et al., 2007) and found to localize at lipid droplets (Suzuki et al., 2015), ER (Suzuki et al., 2015), rods and rings (Schiavon et al., 2018), and mitochondria (Schiavon et al., 2019). Among its first known cellular functions was in mediating mitochondrial fusion as an ARL2 effector (Newman et al., 2017a; Newman et al., 2017b; Newman et al., 2014; Schiavon et al., 2019). Recent studies from our lab, though, revealed that ELMOD2 also acts with ARL2 on aspects of microtubule biology and with ARF6 in cytokinesis/abscission (Turn et al., 2020). This recent study also provided evidence that ELMOD2 localizes to centrosomes and Flemming bodies, consistent with its effects on microtubules and abscission. These novel and unexpected roles were found in cells deleted for ELMOD2 using the CRISPR/Cas9 system in immortalized mouse embryonic fibroblasts (MEFs). These lines were generated in part due to our inability to document its knockdown by siRNA because of its low abundance in cultured cells (Turn, et al 2020). Interestingly, deletion of neither ELMOD1 nor ELMOD3 in MEFs resulted in any of the phenotypes described previously or below (manuscript in preparation), suggesting a high degree of specificity of ELMOD2 within this small family.

With the knowledge that (1) ELMOD2 localizes to centrosomes, (2) many regulators of cell cycle also have close links to cilia, (3) multiple ARF family members (including at least ARL2/3/6/13B) are implicated in ciliary signaling (Fisher et al., 2020), and (4) ELMOD2 has in vitro GAP activity for at least two of these ciliary ARFs, we hypothesized that ELMOD2 also may play a role in ciliary function. Primary cilia serve as signaling hubs that mediate essential intracellular and intercellular functions, particularly during development (Gigante and Caspary, 2020; Goetz and Anderson, 2010; Huangfu et al., 2003; Pazour and Rosenbaum, 2002). Within the past few decades, there has been a steady increase in the study of primary cilia because of their link to a range of human pathologies. These diseases, collectively called ciliopathies, include polycystic kidney disorder, Bardet-Biedl Syndrome, situs inversus, primary cilia dyskinesia, and Joubert Syndrome, as well as others (Chen et al., 2020; Goetz and Anderson, 2010; Waters and Beales, 2011). Further studies have implicated primary cilia as signaling hubs, sequestering receptors needed for development, metabolism, recognition of sensory stimuli, cell cycle, and others (Nachury and Mick, 2019; Reiter and Leroux, 2017).

Primary cilia are composed of (1) a basal body tethered to the plasma membrane by pinwheel-like structures called distal appendages, (2) microtubules that project from the distal end of the basal body to create a single, intact axoneme, and (3) a ciliary membrane encasing the axoneme as it projects into the extracellular space. Cells grown in culture typically lack cilia until they approach confluence, or enter G_0_, which is promoted by serum starvation. Primary ciliogenesis is tightly regulated to ensure that one and only one primary cilium is formed per cell. For many cell types (excluding RPE1 and NIH3T3 cells (Munger, 1958; Sorokin, 1962; Wang and Dynlacht, 2018)), the intracellular pathway of ciliogenesis involves a series of incompletely understood steps that include movement of centrosomes toward the cell surface where the mother centriole becomes established as the basal body. During this process, the centrosomal protein Cep164 is recruited to the distal appendages of the mother centriole (Cajanek and Nigg, 2014; Graser et al., 2007a; Schmidt et al., 2012), giving them license to recruit TTBK2 (Bouskila et al., 2011; Cajanek and Nigg, 2014; Goetz et al., 2012), a kinase that phosphorylates Cep83 (Lo et al., 2019) and MPP9 (Huang et al., 2018). These factors lead to the release of the capping protein complex CP110-Cep97 (Huang et al., 2018; Spektor et al., 2007). Afterward, ciliary vesicles dock at the basal body and proceed with building the transition zone and extending the axoneme to generate the elongating cilium. The commitment to initiate ciliogenesis, also called licensing, is often monitored by the obligate recruitment of Cep164 and later release of CP110 as markers of specific steps in this process.

Both during ciliogenesis and in existing cilia, the appropriate localization and distribution of proteins is critical for ciliary functions. Proteins can either diffuse through the transition zone (TZ, the presumptive physical barrier) or enter via active transport. In the case of active transport, cargoes are carried by at least three protein complexes: IFT-A, IFT-B, and the BBSome. Once inside the cilium, IFT complexes and their cargos can be actively transported along the axoneme via kinesin or dynein driven motors. The selective traffic in and out of cilia provides an exclusive environment where signaling components are highly enriched due to the confined intraciliary space (Nachury 2014). Of the signaling pathways linked to cilia, the best known is the Sonic Hedgehog (SHH) pathway (Huangfu et al., 2003). Key components of the SHH pathway localize to the ciliary membrane, and several change dynamically in response to SHH ligand: the SHH receptor Patched1 (Ptch1) and GPR161 each act as negative regulators of the pathway and exit the cilium, the G-protein coupled receptor Smoothened (Smo) is recruited into cilia, and the Gli transcription factors are proteolytically processed and exit (Corbit et al., 2005; Haycraft et al., 2005; Mukhopadhyay et al., 2013; Pal et al., 2016; Rohatgi et al., 2007; Rohatgi and Scott, 2007). Other signaling pathway components also localize to the ciliary membrane, including the ARF family GTPase ARL13B, Somatostatin Receptor 3 (SSTR3), and adenylyl cyclase III (ACIII). Indeed, ARL13B is commonly used as a marker of cilia (along with acetylated tubulin) due to its strong signal in cell imaging. Even less well understood than traffic in and out of cilia is specific transport of newly synthesized proteins from the ER, through the Golgi, to cilia. Such ciliary traffic may be targeted directly to the basal body for regulated import, but some have also implicated a role for rootlets in traffic to cilia (Yang and Li, 2005).

Rootlets are cytoskeleton-like structures that project from the proximal end of the basal body and are thought to be composed primarily of the ~225 kDa protein Rootletin. The *Crocc* gene encodes Rootletin (**<u>c</u>**iliary **<u>ro</u>**otlet **<u>c</u>**oiled-**<u>c</u>**oil). Because the protein is consistently termed Rootletin in the literature, we will conform to this usage. Other proteins reported to bind and localize to rootlets include kinesins, amyloid precursor protein (APP), and presenilins (Yang and Li, 2005). The rootlet’s function is incompletely understood but is proposed to help stabilize cilia against external flow and to regulate ciliary traffic (Yang et al., 2005; Yang and Li, 2005; Yang and Li, 2006; Yang et al., 2002). Rootlets are also important in centrosomal cohesion, along with the centrosomal proteins C-NAP1, Cep68, and Cep44 (which is believed to help anchor the rootlet to the centrosome) (Hossain et al., 2020). Any functional relationship(s) between the roles of Rootletin in ciliary biology and centrosome cohesion have not been described previously, to our knowledge.

Our work has uncovered a novel aspect of regulation of ciliary licensing and function in ELMOD2 with links to Rootletin. To test the model that ELMOD2 plays a role at cilia, we used our previously generated ELMOD2 KO mouse embryonic fibroblast (MEF) lines (Turn et al., 2020). We discovered novel roles for ELMOD2 at cilia and ciliary rootlets and resolved these effects from ELMOD2’s roles in microtubule and mitochondrial functions as well as cytokinesis. We also uncovered close functional links between ELMOD2 and Rootletin in both centrosome cohesion and ciliogenesis, with links also to ARL2. Finally, using previously characterized markers of specific steps in ciliary licensing, we identify the site or step(s) at which we propose that ELMOD2 and Rootletin act in ciliogenesis. Together, we believe that these data provide several new insights into fundamental aspects of ciliary biology, including ciliary licensing and rootlet function.

## Results

### ELMOD2 deletion causes increased ciliation and multiciliation

We have shown previously (Turn et al., 2020) that the introduction of frame-shifting mutations into both alleles results in functional nulls in ELMOD2 (which we will term ELMOD2 KO for the sake of simplicity) in immortalized MEFs. Loss of ELMOD2 results in centrosome amplification as well as decreased microtubule stability and nucleation from centrosomes. Given the roles of centrosomes/basal bodies and microtubules in ciliogenesis and ciliary functions, we explored effects of ELMOD2 deletion on ciliation in these cells, predicting there to be defects/loss in ciliation. We used the same 10 KO clones described previously (Turn et al., 2020) in which frame shifting mutations in ELMOD2 were introduced by CRISPR/Cas9 and confirmed by DNA sequencing, as well as both a parental and a “CRISPR WT” line that underwent transfection and cloning but had no mutations in the targeted region of the ELMOD2 gene. Four of the KO clones were transduced with a lentivirus directing expression of ELMOD2-myc to assess rescue of any observed phenotypes and to protect against off-target effects of CRISPR. Note that we observed positive myc staining in ~70-90% of lentiviral transduced cells and that scoring of such rescued lines was done by counting all cells, likely explaining the incomplete rescue observed in most cases (though always close to WT levels).

We stained cells for ciliary (ARL13B, acetylated tubulin) and centrosomal (γ-tubulin, centrin) markers under normal growth conditions (10% fetal bovine serum (FBS)) and after serum starvation (0.5% FBS for 24 hr) to induce ciliation, as described under Materials and Methods. In contrast to our prediction, that was based on microtubule defects (Turn et al., 2020) and their role in the axoneme, ELMOD2 KO lines displayed *increased* ciliation compared to WT controls (Figure 1A) in both normal medium and after serum starvation. All 10 ELMOD2 KO lines displayed higher rates of ciliation compared to WT, both with (89.1% versus 42.7%, respectively; Figure 1B) or without (62.5% versus 16.0%, respectively; Figure 1C) serum starvation to induce ciliation. This increase in the percentage of ciliated cells in ELMOD2 KO lines was reversed upon expression of ELMOD2-myc via lentivirus. These “rescued” KO lines had clearly reduced ciliation compared to their uninfected KO cells: decreasing from 89.1% to 42.8% upon rescue with serum starvation (Figure 1B), and from 62.5% to 28% in normal serum conditions (Figure 1C), in each case approaching numbers seen in WT lines. In contrast, there was no difference in the percentage of ciliated cells in WT compared to WT transduced with the lentivirus directing expression of ELMOD2-myc, indicating that overexpression of ELMOD2-myc does not overtly impact the extent of ciliogenesis (Fig. 1B-C).

**Figure 1:**
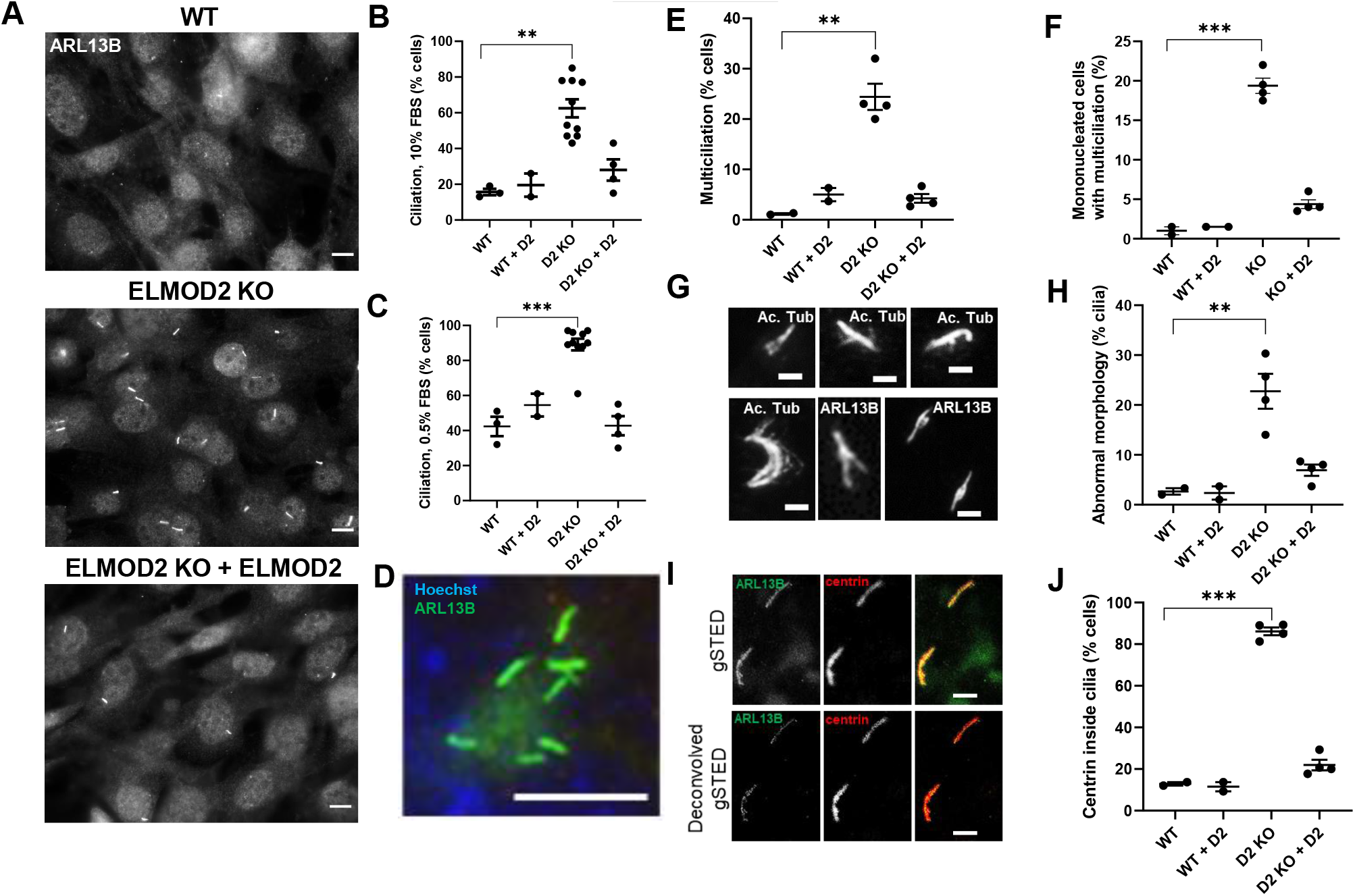
Deletion of ELMOD2 causes ciliary defects. **(A)** ELMOD2 KO cells display increased ciliation and multiciliation, compared to WT MEFs. Cells were grown to ~80% confluence, fixed with 4% PFA, permeabilized with 0.1% Triton X-100, and stained for ARL13B as a marker of ciliation. Representative images were collected at 60x magnification using widefield microscopy. Scale bar = 10 μm. **(B)** Using the same conditions described in (A), ciliation was scored in 2 WT, 10 ELMOD2 KO, and 4 ELMOD2-rescued lines. One hundred cells per cell line were scored for the presence of one or more cilia. ARL13B and acetylated tubulin were used as markers to detect cilia. (**C**) The same experiment was performed as described for (B), except cells were serum starved and plated at 90-100% confluence. **(D)** Loss of ELMOD2 leads to increased multiciliation. Cells were fixed with 4% PFA, permeabilized with 0.1% Triton X-100, and stained for ARL13B to detect cilia and with Hoechst to identify individual cells. Images were collected using widefield microscopy at 100x magnification. Scale bar = 10 μm. **(E)** The same experiment was performed as described for (C), except multiciliation (>1 cilia) was scored. **(F)** The same experiment was performed as described for (E), except multiciliation was only scored in mononucleated cells with only 1-2 centrosomes. This was performed to ensure that the multiciliation phenotype was not simply a consequence of cell cycle defects. **(G)** Examples of cilia with abnormal morphology are shown. Images were collected using widefield microscopy at 100x magnification, highlighting the branching/splaying. Panels are labeled to indicate whether ARL13B or acetylated tubulin staining is shown, though no differences were noted. Scale bar = 2 μm. **(H)** Serum-starved cells stained for ARL13B and acetylated tubulin were scored for abnormal morphology (*i.e*., branching or splaying). Only ciliated cells were scored. **(I)** g-STED microscopy (100x magnification) confirms the localization of centrin to cilia in ELMOD2 KO cells. The two cilia shown in this image are in a single cell. These cilia have centrin localization along the length of the cilium as well as at buds. **(J)** Percentage of cells with cilia positive for centrin were scored. Experiments were performed in triplicate, and the average of the triplicate for each line was plotted. Results were tabulated in an interleaved scatterplot via GraphPad Prism. Statistical significance was assessed using One-Way ANOVA; *=p<0.05; **=p<0.01; ***=p<0.0001.

In addition to increased frequency of ciliation, we also observed clear increases in instances of multiciliation. This is unusual, as WT MEFs typically have a single primary cilium with uniform staining of acetylated tubulin and ARL13B throughout its length, and each cilium has a single basal body. The four ELMOD2 null lines examined revealed an increase in multiciliation, with an average of 24.4% of ciliated cells having at least two cilia, compared to only 1.2% of WT cells (Figure 1D-F). Though it was far more common to see 2-3 cilia per cell, some cells had 9 or more cilia (see Figure 1D). We previously reported (Turn et al., 2020) that ELMOD2 KO cells have supernumerary centrosomes at an increased frequency, which may enable this increase in ciliation. To examine whether the increased ciliation is solely a consequence of cell cycle defects, we repeated the scoring in cells with no evidence of cell cycle defects (i.e., one nucleus of normal size and morphology, only 1-2 centrosomes). Even after restricting the phenotyping to ELMOD2 null cells of normal cell cycle morphology, 19.4% of ciliated ELMOD2 nulls are multiciliated (Figure 1F). Thus, ELMOD2 appears to play a role in suppressing ciliation, as the loss of ELMOD2 in MEFs results in both an increase in the frequency at which cells ciliate and also alterations in the processes that control ciliation numbers per cell.

### ELMOD2 KO cells display abnormal ciliary morphology and protein content

We next asked if the cilia in ELMOD2 KO cells displayed normal morphology and function, as we predicted that spurious ciliation may be accompanied with failed regulation of ciliary structure. We noted an increase in frequency of cilia displaying non-uniform ARL13B and/or acetylated tubulin staining. Using structured illumination microscopy (SIM), these abnormalities in staining could be resolved into what appear to be buds coming off the surface along the length of the cilium (Figure S1). There are even instances in which these buds form branches or result in ciliary splaying, as detected even by widefield microscopy (Figure 1G). Individual buds and branches that stained positive for ARL13B often, but not always, co-stained with acetylated tubulin. As ARL13B is associated with the ciliary membrane and acetylated tubulin is in the axoneme, these data suggest that these aberrant morphologies either may occur from the ciliary membrane alone or could also be in response to changes in the axonemal structure. While 22.8% of cilia in ELMOD2 KO cells have abnormal morphology, this is true in only 2.7% of cilia in WT MEFs based on widefield immunofluorescence imaging (Figure 1H).

We typically use centriolar markers, such as centrin, to mark and facilitate the identification of cilia, particularly when ciliary markers display high background (cytosolic) staining. A surprising but common change found in the ELMOD2 KO lines (Figure 1I) was the presence of centrin staining *inside* cilia in an average of 86.2% of KO cells versus 12.8% of WT cells (Figure 1J). Centrin is a canonical centriolar protein used to mark basal bodies and previously has not been reported in primary cilia, except in the transition zone of retinal cells (Wolfrum, 1995), though its presence in motile cilia has been reported (Huang et al., 1988; Piperno et al., 1992). Because of the striking increase in centrin staining inside cilia, we asked if there were conditions in which centrin staining becomes more prominent in WT cilia. That is, perhaps centrin routinely enters cilia but is rapidly exported so it only rarely reaches levels detectible by antibody staining. To test this, we inhibited retrograde ciliary transport by treating cells with the dynein motor inhibitor ciliobrevin (30μM for 1 hour) before fixing cells and staining for centrin. As a positive control, we stained for Gli3 with and without ciliobrevin treatment (Firestone et al., 2012). Without ciliobrevin, little to no Gli3 staining was present in cilia of WT or ELMOD2 null cells. However, ciliobrevin treatment led to increased Gli3 ciliary staining and, in many cases, Gli3 accumulation at the ciliary tip in both WT and ELMOD2 KOs. When we stained ciliobrevin-treated cells for centrin, we detected centrin in WT cilia at levels comparable to those seen in cilia of ELMOD2 KO lines without ciliobrevin treatment (Figure S1B). Ciliobrevin-treatment of ELMOD2 KOs resulted in even higher levels of centrin staining, including a subpopulation of cells demonstrating accumulation of centrin at the ciliary tip. These data are consistent with the conclusion that centrin enters cilia in WT cells but is normally rapidly exported. In ELMOD2 KO lines, export of centrin appears to be compromised, resulting in its accumulation in cilia, though we cannot exclude potential effects of ELMOD2 deletion on centrin protein half-life or rate of import. Interestingly, no differences were observed in Gli3 staining in WT vs ELMOD2 KO lines, suggesting that there is a level of selectivity to the effects of ELMOD2 KO on the ciliary proteome.

The discovery that ELMOD2 KO cilia have increased centrin staining led us to ask whether traffic of other ciliary proteins is also altered. We began by investigating Smoothened (Smo), which dynamically localizes to the ciliary membrane in response to SHH ligand (Corbit et al., 2005; Polizio et al., 2011). Cells were treated with SHH-conditioned medium under serum starvation conditions and were stained 24 hours later with antibodies directed against acetylated tubulin and Smo, as described under Materials and Methods. Controls included serum starved cells that were not exposed to SHH. WT cells displayed marked accumulation of Smo in cilia upon SHH treatment (Figure 2A) with 76.5% demonstrating strong staining, 15.3% weak staining, and 8.0% no Smo staining, determined as described under Materials and Methods. In contrast, ELMOD2 KO lines showed markedly reduced Smo staining in SHH-treated cells, with only 26.4% strong, 6.6% weak, and 67.0% having no evident Smo staining in cilia (Figure 2A-B). Expression of ELMOD2-myc reversed the defect in Smo recruitment seen in ELMOD2 KO lines (Figure 2B).

**Figure 2:**
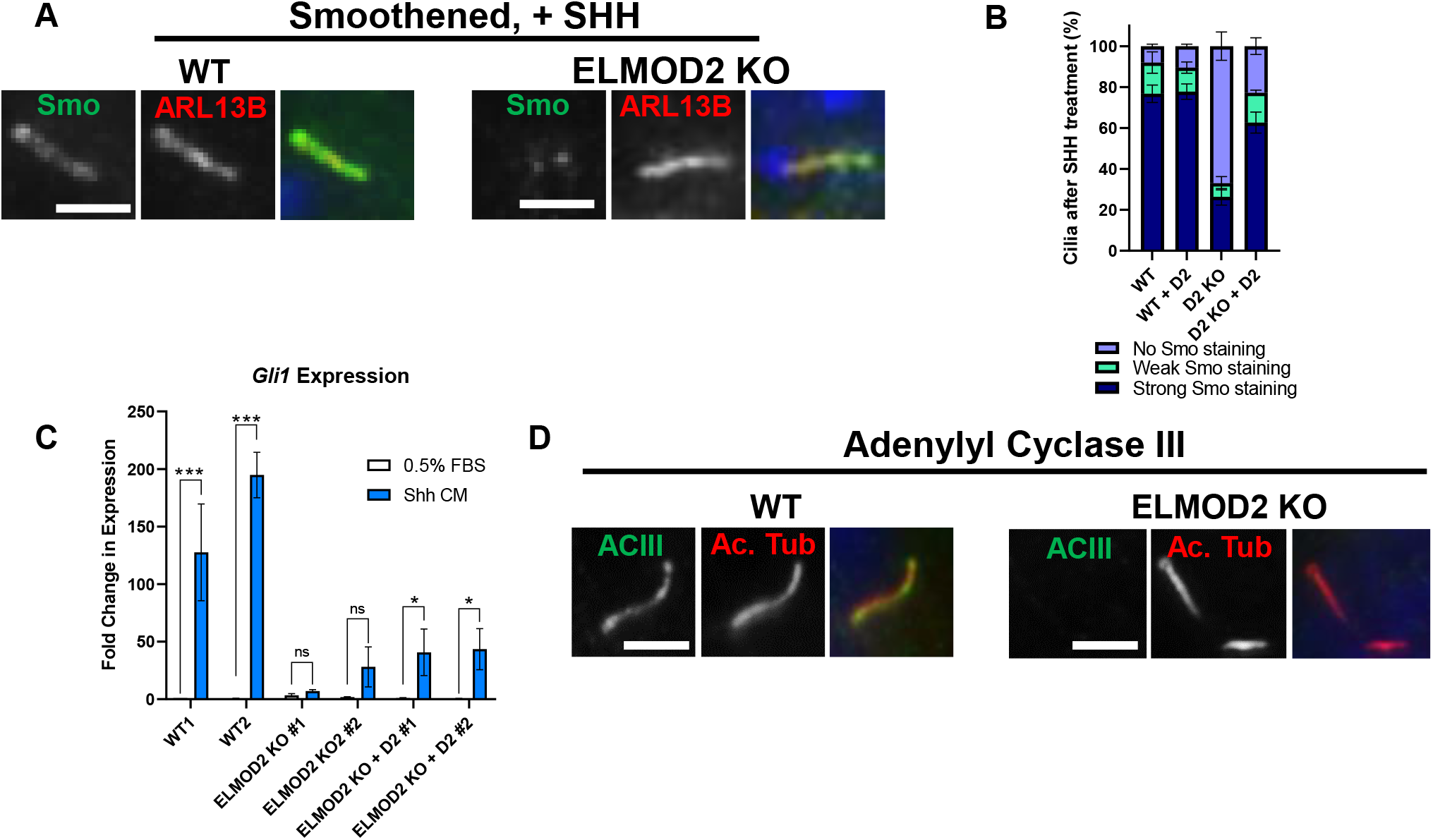
Ciliary signaling is disrupted in ELMOD2 KO lines. **(A)** ELMOD2 KO cells show decreased Smo recruitment after SHH treatment, compared to WT cells. Cells (2 WT, 4 ELMOD2 KO, and 4 ELMOD2 KO + ELMOD2-myc) were serum starved, treated with SHH-enriched medium for 24 hours to induce, fixed with 4% PFA, and permeabilized with 0.1% Triton X-100. Cells were co-stained for Smo, ARL13B, and Hoechst. Scale = 10μm. **(B)** Cells were stained as described in (A) and 100 were scored per line in triplicate. Ciliated cells were binned into either having strong, weak, or no Smo staining, as described under Materials and Methods. The average of the triplicates for each line was determined, and the data were plotted as a stacked bar graph. Error bars indicate SEM. **(C)** ELMOD2 KO MEFs show reduced SHH-stimulated *Gli1* transcriptional response, compared to WT cells. Cells were collected 48 hours after SHH treatment and levels of *Gli1* mRNA were determined using qPCR. Data are presented as mean fold change ± standard deviation (SD), and bar graphs indicate normalized mRNA expression. Statistical significance was assessed using Two-Way ANOVA; *=p<0.05; ***=p<0.0001, ns = not significant. **(D)** ELMOD2 KO cells show reduced recruitment of ACIII. Serum-starved cells were fixed and stained for ACIII, as described under Materials and Methods. Representative images were collected via widefield microscopy at 100x magnification. Scale bar = 10 μm.

Because Smo enrichment was defective, we next tested whether SHH-dependent signaling was also compromised by monitoring changes in the transcription of *Gli1*, a transcriptional target of the SHH pathway that is normally increased in response to SHH signaling. Using qPCR, we measured *Gli1* mRNAs in cells treated with 0.5% FBS with or without SHH conditioned media. In parental and control cells, we saw robust SHH-dependent increases in *Gli1* expression (Figure 2C). In contrast, ELMOD2 KO cells did not show a statistically significant increase in transcript levels above their baseline. The addition of ELMOD2 into ELMOD2 KO cell lines restored SHH responsivity, with ELMOD2-myc expression leading to SHH-dependent increases in *Gli1* above their respective baseline levels. However, the transcript levels in ELMOD2 KO+ELMOD2-myc cells were not increased above their SHH-treated KO counterparts, suggesting that the normal Smo enrichment permitted only a partial recovery of SHH-dependent gene transcription (Figure 2C). Because these cells are polyploid and multinucleated, due to cell cycle defects described previously (Turn et al., 2020), it is difficult to draw many conclusions regarding the deficient SHH signaling and downstream transcriptional output in ELMOD2 KO cells. Together, though, both the immunofluorescence and qPCR data monitoring the SHH pathway reveal defects in ciliary signaling.

To determine whether the defect was specific to the SHH pathway, we also looked at other ciliary membrane proteins. The GPCRs SSTR3 and GPR161 (see Figure S2A-D for images and scoring) as well as adenylyl cyclase ACIII (Figure 2D) each displayed reduced ciliary localization in ELMOD2 KO compared to WT cells. These deficiencies in ciliary receptors were each reversed upon expression of ELMOD2-myc. Together, these results show that deletion of ELMOD2 leads to alterations in the levels of several ciliary signaling proteins. These changes include both increases (*e.g*., centrin) and decreases (*e.g*., SSTR3, ACIII, GPR161) in protein abundance that might result from changes in import, export, or protein half-life.

### ELMOD2 localizes to the basal body in MEFs and can be found in cilia after treatment of cells with ciliobrevin

In previous studies, we and others have found that ELMOD2 localizes to the endoplasmic reticulum (ER), lipid droplets, mitochondria, Flemming bodies, and centrosomes (Newman et al., 2014; Suzuki et al., 2015; Turn et al., 2020). Based on the phenotypes described above and its known localization at centrosomes, we examined whether ELMOD2 may be retained at basal bodies or present in cilia. Using our rabbit polyclonal ELMOD2 antibody, we found no convincing evidence of ELMOD2 localization to cilia after 24-hour serum starvation and fixation with either methanol or PFA. We then asked if ELMOD2 may behave like centrin (*i.e.*, showing increased staining in cilia after inhibition of retrograde traffic via ciliobrevin treatment). We incubated serum-starved WT MEFs with or without 30 μM ciliobrevin for 1hr, fixed cells with 4% PFA, and immunostained for ELMDO2 and ARL13B. No sign of ELMOD2 staining was seen in WT MEFs without drug treatment. However, ELMOD2 staining was evident, though weak, in cilia from WT cells treated with ciliobrevin (60.5% of WT cilia after ciliobrevin treatment versus 10.0% of untreated cells) (Figure S3A-B). This staining was completely absent in ELMOD2 KO lines, providing further evidence of specificity of the antibody used. It is uncertain whether ELMOD2 functions inside cilia, but these results indicate that it can at least transiently localize there.

To see whether ELMOD2 localizes to basal bodies, we serum starved WT MEFs and immunostained for acetylated tubulin (to mark cilia) and ELMOD2. We observed specific staining of ELMOD2 at both the basal body (Figure 3A) as well as at non-ciliary centrosomes (as previously reported (Turn et al., 2020)). Our previous studies reported strong co-localization with both γ-tubulin and centrin, though appearing more similar in size/shape with γ-tubulin, a pericentriolar material (PCM) marker (Turn et al., 2020).

**Figure 3:**
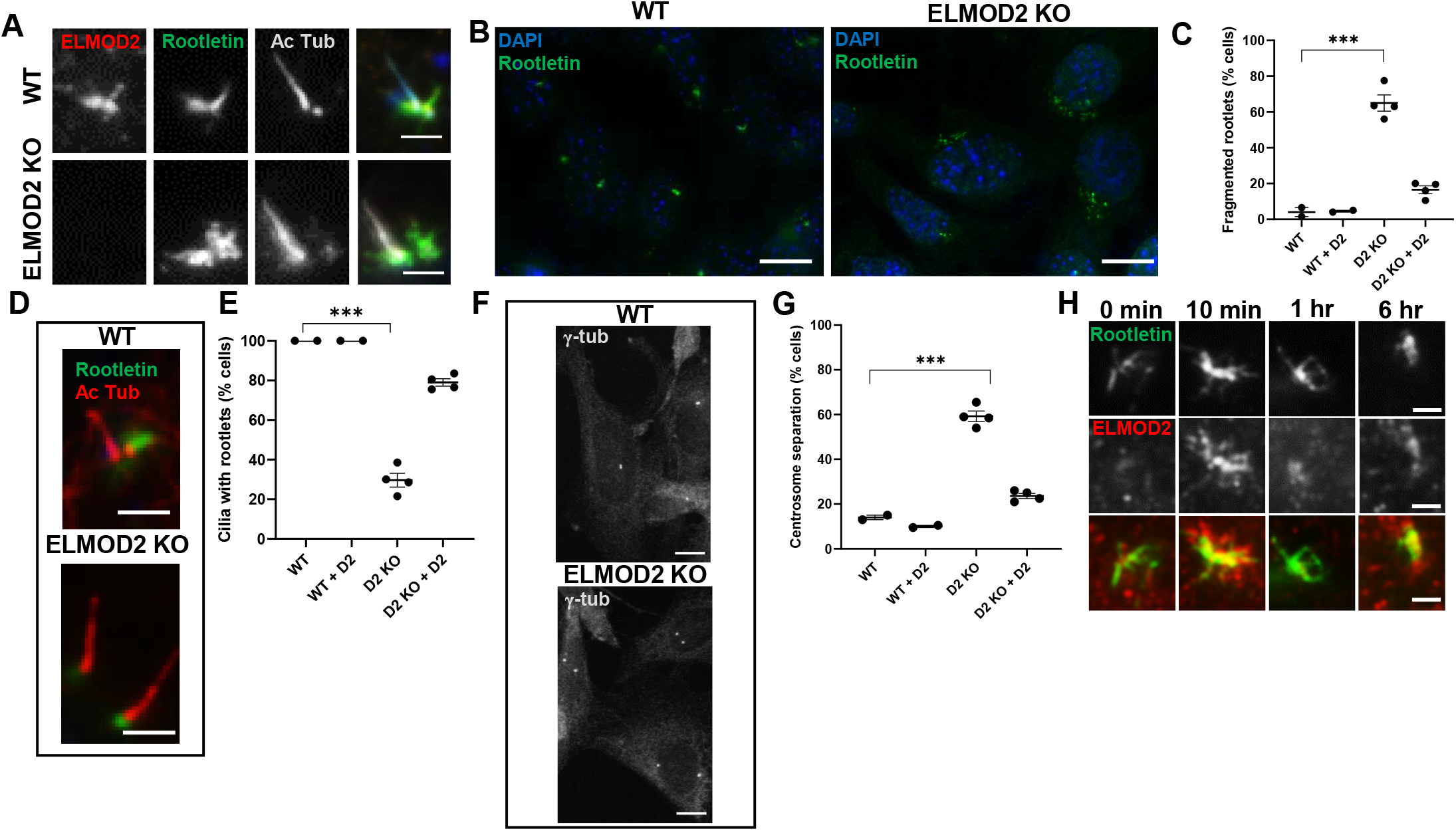
ELMOD2 localizes to rootlets, and its deletion causes rootlet defects. **(A)** ELMOD2 localizes to rootlets in WT MEFs. WT or KO cells were fixed for 5 min in ice-cold methanol and stained for ELMOD2, acetylated tubulin, and Rootletin, as described under Materials and Methods. Images were collected via widefield microscopy at 100x magnification. Scale = 10 μm. **(B, C)** ELMOD2 KO cells have increased rootlet fragmentation. Serum-starved, methanol fixed cells were stained for Rootletin and Hoechst. Images were collected using widefield microscopy at 100x magnification and (**C**) scored in duplicate for fragmented rootlets. **(D)** Rootletin staining in ELMOD2 KO cells is limited to the base of cilia and is more condensed than in WT cells. Growth and fixation conditions were the same as in (B). Cells were stained with Rootletin and acetylated tubulin (to mark cilia). Images were collected via widefield microscopy at 100x magnification. **(E)** The same conditions as described for (C) were used to score cell lines for cilia with rootlets. Only ciliated cells were scored. **(F)** ELMOD2 KO cells show increased centrosome separation. Serum-starved cells were fixed with ice-cold methanol, stained for γ-tubulin, and imaged via confocal microscopy at 100x magnification, with z-projections. Scale bar =10 μm. **(G)** Using the same conditions described in (B), cells were scored for centrosome separation using FIJI image processing software with the provided measuring tool. Cells were counted as “separated” if they were more than 2μm apart. **(H)** ELMOD2 and Rootletin staining each change after serum starvation. WT MEFs were fixed at different times after serum starvation and stained for ELMOD2 and Rootletin. Representative widefield images were collected at 100x magnification. Staining of each at basal bodies is strongly increased within 10 min, showing extensive overlap. At later times each becomes more concentrated into a smaller area, but filamentous staining of ELMOD2 is lost before that of Rootletin. When scoring was performed, the average of duplicates of individual lines were plotted using an interleaved scatterplot. Error bars indicate SEM. Statistical significance was assessed using One-Way ANOVA; ***=p<0.0001.

Interestingly, we noted that ELMOD2 staining at the basal body did not localize exclusively to a tight focus, but also showed staining of a structure, emanating apparently from the proximal end of the basal body, away from the cilium (Figure 3A). Though there are many instances in which ELMOD2 staining appeared as a single protrusion, extending from the basal body at a clearly distinct angle from the ciliary axoneme (acetylated tubulin staining), in other cases multiple smaller, fibrillar projections were apparent, and these varied in length. This staining was absent in ELMOD2 KO cells (Figure 3A). These projections did not co-localize with acetylated tubulin, suggesting that ELMOD2 was localizing to a distinct structure. Based on this staining, we predicted that perhaps ELMOD2 also localizes to ciliary rootlets, a cytoskeletal structure projecting from basal bodies and made up of polymers of the protein Rootletin. To test this, serum-starved WT MEFs were costained with acetylated tubulin (to mark cilia), Rootletin (to mark rootlets), and ELMOD2 to assess colocalization. Rootletin staining is apparent in all cells, with or without cilia, and strongly concentrated at centrosomes. In practically all cells studied, this staining appears as long tendrils/projections surrounding and extending from the centrosome, as described previously (Conroy et al., 2012; Flanagan et al., 2017; Vlijm et al., 2018). There was heterogeneity in the morphology of these rootlets, as some have many protrusions, and their length and shape varied from cell to cell. When comparing rootlets in ciliated versus non-ciliated cells, we note that rootlets at the base of cilia typically appeared as one, thick rootlet rather than many thinner, more tendril-like rootlets that were more typical of non-ciliary rootlets. Interestingly, ELMOD2-positive “projections” clearly overlapped and partially co-localized with the Rootletin fibers/feet (Figure 3A). Not only did their staining not completely overlap, but there were cases in which there were Rootletin-positive tubules extending from the basal bodies that showed no sign of ELMOD2 staining. ELMOD2 staining tends to co-localize with rootlets only when the rootlets are associated with the centrosome, and co-localization is most extensive when the strands of Rootletin are compact (Figure S4).

### Loss of ELMOD2 leads to fragmentation and abnormal morphology of ciliary rootlets

The discovery that ELMOD2 partially localizes to ciliary rootlets led us to ask whether loss of ELMOD2 disrupts rootlet organization or function. Using the same conditions described above to look at ciliary rootlets in WT MEFs, we repeated the experiment in ELMOD2 KO lines and found striking differences. In general, ELMOD2 KO lines displayed more fragmented Rootletin staining throughout the cell body, rather than bright fibrillar staining focused at centrosomes (Figure 3B). On average, 65.0% of KO cells show rootlet fragmentation (*i.e.*, the Rootletin staining is dispersed throughout the cell as bright puncta), while only 4.0% of WT cells had such fragmented rootlets (Figure 3C). Expression of ELMOD2-myc in the KO lines brought the extent of rootlet fragmentation back down to near WT levels (12.5%; Figure 3C). In ciliated ELMOD2 KO cells, Rootletin stains the proximal end of basal bodies as bright puncta but without evident rootlets, or what we define as a protrusion projecting from the base of the cilium (Figure 3D). While 100.0% of WT cilia have a rootlet, only 29.6% of ELMOD2 KO cilia have a rootlet (Figure 3E). Once again, this phenotype is largely rescued upon expression of ELMOD2-myc (79.0%) (Figure 3E). These data provide evidence that ELMOD2 is important in regulating rootlet recruitment to or organization at basal bodies.

Previous studies have revealed that Rootletin is a critical component in centrosome cohesion (or the linkage of two centrosomes to one another that dynamically changes during different stages of cell cycle), along with C-Nap1, Cep44, and Cep68 (Bahe et al., 2005; Conroy et al., 2012; Flanagan et al., 2017; Graser et al., 2007b; Hossain et al., 2020; Vlijm et al., 2018; Yang et al., 2006). Disruption of centrosome cohesion can lead to spurious centrosomal separation with potentially severe downstream consequences in the cell, such as aneuploidy and supernumerary centrosomes (Yang et al., 2006). We tested whether loss of ELMOD2 leads to increased centrosome separation by scoring the number of cells with centrosomes >2μm apart, a common metric in the field (*e.g.*, see (Bahe et al., 2005)). An average of 14.0% of WT cells had separated centrosomes, while 59.3% of ELMOD2 null cells had separated centrosomes (Figure 3F-G). The centrosome cohesion defect was largely reversed upon expression of ELMOD2-myc (23.6%) (Figure 3G). Thus, ELMOD2 plays a role in docking or retention of Rootletin at centrosomes. Based on these data, we conclude that ELMOD2 is important to both centrosome cohesion and ciliary licensing.

### ELMOD2 localization and rootlet morphology change dynamically during early ciliogenesis

We also noted that ciliated cells appeared to have tighter, more compact rootlets that extended from the centrioles in the basal body. In contrast, non-ciliated cells appeared to have larger, more spread-out rootlets encasing their centrosomes and extending thin tendrils into a larger area. Rootlets are dynamic structures, particularly with respect to centrosome separation (Mahen, 2018). Yet, the relationship between Rootletin dynamics and cilia-inducing conditions apparently has not been previously described. We performed live-cell imaging of Rootletin using widefield microscopy of GFP-Rootletin transfected MEFs at low magnification (20x) (Figure S5). Wild-type cells (both clonal and mother) were imaged with and without serum starvation to observe if serum starvation promotes changes in Rootletin morphology. As shown in Figure S5, rootlet fragmentation was observed upon induction of serum starvation (0.5% FBS), in which regions of strong Rootletin staining began to separate from the centrosomes and PCM. These fragmentation events began within the first ~15 minutes of imaging and were only observed after serum starvation. Swapping in fresh medium (10% FBS) caused no evidence of changes in rootlet morphology during the hour-long imaging window (Figure S5).

Based on the finding that rootlets dynamically change morphology in response to serum starvation, we asked if ELMOD2 also changed in localization to rootlets after serum starvation. Under normal conditions (10% FBS), rootlets typically appeared as large, dense networks of anemone-like structures encasing both centrosomes, while ELMOD2 staining only partially co-localized with that of Rootletin and is relatively weak in intensity often making it difficult to discern at basal bodies over background (Figure 3H, 0 min). Cells were fixed at different time points after serum starvation and stained for Rootletin, ELMOD2, and acetylated tubulin (Figure 3H). These experiments were repeated in triplicate with both mother WT and clonal WT lines, collecting time points at 0 min, 10 min, 20 min, 30 min, 40 min, 50 min, 1hr, 3hr, 6hr, and 24 hr. To avoid complications resulting from (over)expression of tagged protein, these experiments were done on fixed cells stained for endogenous proteins. And because the morphologies involved are highly dynamic and not uniform we chose not to score the differences. However, clear patterns of changes were evident, with strong enhancement of ELMOD2 at rootlets at early times (10 min), followed by diminution but retention of some staining at centrosomes above that seen in cells not serum starved, persisting throughout the time course. Rootletin staining at centrosomes follows a very similar pattern, with strongly increased staining at 10 min, diminishing over the rest of the time course, but at a slower rate, compared to ELMOD2. About 50% of WT cells display this strongly increased staining of both ELMOD2 and Rootletin at 10 min after initiation of serum starvation, similar to the ultimate percentage of cells that become ciliated (Figure 1C). Also, at later time points the vast majority of cells with compacted rootlets were ciliated while cells that still had less-compact, tendril-like rootlets lacked a cilium. Thus, we believe these are early events in the ciliogenesis pathway. A more detailed study at earlier time points, using fluorescent proteins may reveal the order of recruitment of these proteins to centrosomes but was not undertaken in our studies.

### Rootletin KO MEFs phenocopy several ELMOD2 KO phenotypes

As a result of the partial co-localization of ELMOD2 and Rootletin and effects of ELMOD2 deletion on rootlets, we sought additional evidence to test a model in which ELMOD2 and Rootletin act in the same pathway to regulate ciliogenesis. We again used CRISPR/Cas9 genome editing to generate MEF lines deleted for Rootletin as a consequence of the introduction of frameshifting insertions and deletions. We generated five clonally derived lines in which both alleles were shifted, using three different guides that each target a different exon. In this case we first screened for Rootletin KO by immunofluorescence and later used DNA sequencing around the targeted exons to confirm frameshifting occurred in both alleles, as shown in Figure S6A-B. Mouse Rootletin is composed of 2009 amino acids encoded in 37 exons on chromosome IV. NCBI predicts (*Crocc* gene ID: 230872) two transcripts that differ at the 5’ end, impacting N-terminal sequences. For this reason, we targeted exons downstream of these differences so that both transcripts would be disrupted. We confirmed the loss of Rootletin by immunoblotting total cell lysates with an antibody specific for Rootletin and confirmed the loss of the major band at ~240 kDa (seen in all WT lines) in all five KO lines (Figure 4A). This Rootletin antibody was raised against a C-terminal fragment of the holoprotein and failed to reveal any shorter bands, consistent with the loss of Rootletin protein products. The lack of immunoreactivity in these cells further confirmed the absence of Rootletin (Figure 4B). Furthermore, we immunoblotted for Rootletin in ELMOD2 KO cells and observed no change in total Rootletin expression (Figure 4, Figure S7). This would suggest that the changes of Rootletin morphology in ELMOD2 KO cells is not simply a side effect of changes in protein expression.

**Figure 4:**
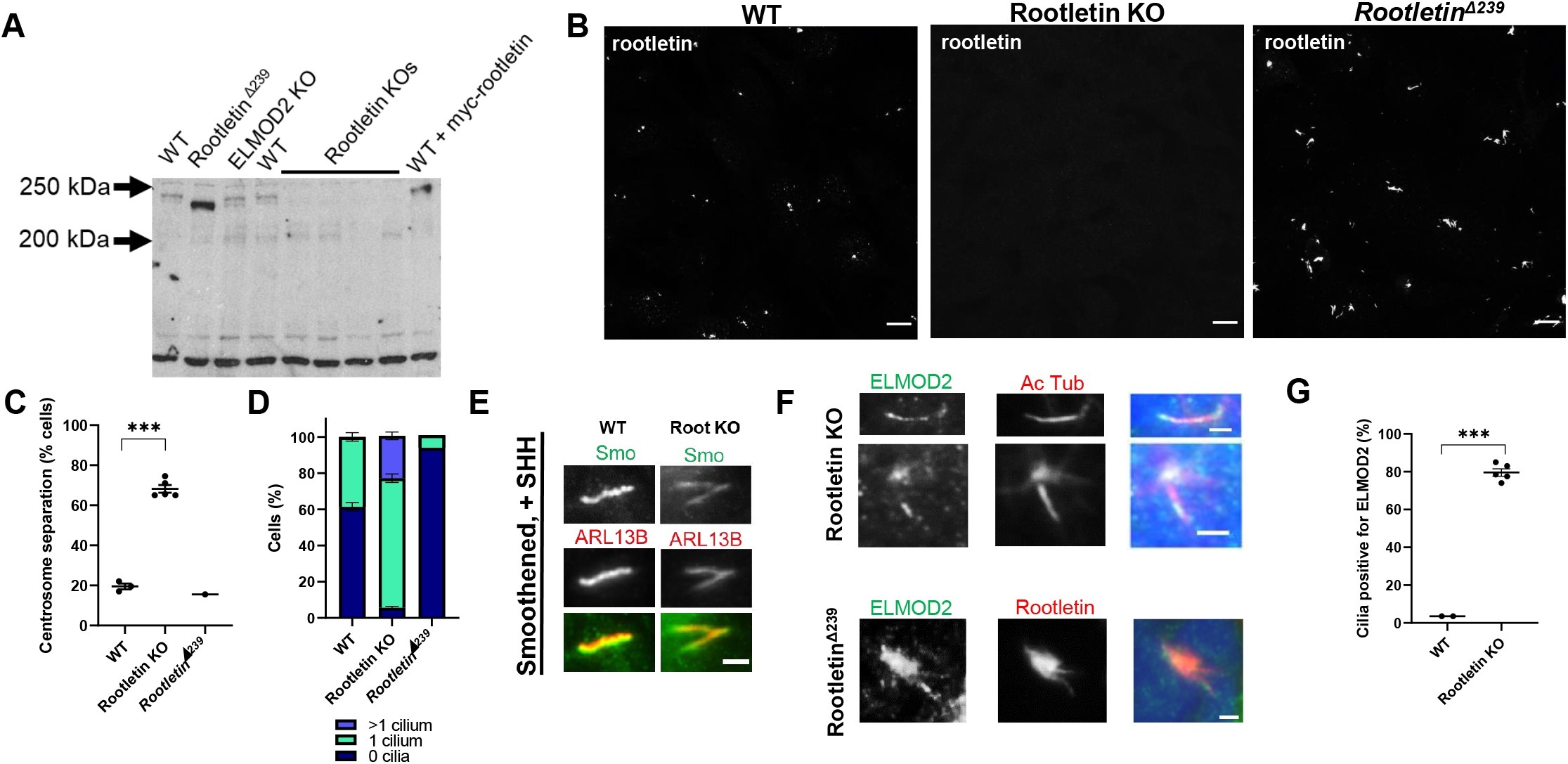
Rootletin KO lines phenocopy ELMOD2 KO ciliary and centrosomal cohesion defects. **(A)** Immunoblotting shows the absence of Rootletin in Rootletin KO, no changes from WT in ELMOD2 KO cells, and strongly increased expression in Rootletin^Δ239^ cells. Equal protein was loaded into a 7.5% polyacrylamide gel before being transferred to nitrocellulose membrane and stained for Rootletin, as described under Materials and Methods. The band migrating at ~240 kDa, based on comparison to protein standards, in WT and ELMOD2 KO MEFs is absent in Rootletin KO lines. This band is increased in intensity upon expression of myc-Rootletin (far right lane). The Rootletin^Δ239^ cell lysate, instead, has a stronger staining band that migrates ~20 kDa faster compared to WT. An image after 1-minute exposure to film is shown. See Fig. S6 for other images. **(B)** Confocal images (100x magnification, z-stacks) of WT, Rootletin KO, and Rootletin^Δ239^ cells stained for Rootletin are shown. Scale = 10μm. **(C)** Rootletin KO cells have increased centrosome separation compared to WT. Cells were fixed with ice cold methanol and stained for γ-tubulin to mark centrosomes. Fields of cells at 100x magnification were taken and processed using FIJI imaging software to measure the distance between centrosomes. Centrosomes that were more than 2μm apart were considered separated. This experiment was performed in duplicate, and the average of the duplicates of each line was plotted in an interleaved scatterplot. Error bars indicate SEM. Statistical significance was assessed using One-Way ANOVA; ***=p<0.0001. **(D)** Serum starved WT, Rootletin KO, and Rootletin^Δ239^ cells reveal that loss of Rootletin leads to increased ciliation while expression of Rootletin [Δ239] prevents ciliation. Cells were stained for acetylated tubulin or ARL13B and scored in duplicate for having either 0, 1, or >1 cilia. Data were graphed in GraphPad Prism using a stacked bar graph. Error bars indicate SEM. **(E)** Serum starved, SHH-treated WT and Rootletin KO cells were stained for Smo and ARL13B. Widefield images collected at 100x magnification are shown. Scale = 2μm. **(F)** ELMOD2 localizes to cilia in Rootletin KO and strongly to rootlets in the Rootletin^Δ239^ mutant. Serum starved cells were stained for ELMOD2 and either Rootletin or acetylated tubulin. Images were collected via widefield microscopy at 100x magnification. Scale = 10μm. **(G)** The Rootletin KO cells described in (F) were scored for % of cells with cilia positive for ELMOD2. The experiment was performed in duplicate, and the average of the duplicates of each line was plotted in an interleaved scatterplot. Error Bars indicate SEM. Statistical significance was assessed using One-Way ANOVA; ***=p<0.0001.

Deletion of Rootletin resulted in increased centrosome separation (68.2% of separated centrosomes in Rootletin KO versus 19.5% in WT), which is consistent with previous reports of Rootletin regulating centrosome cohesion (Bahe et al., 2005; Yang et al., 2006) (Figure 4C). Rootletin KO cells also display increased ciliation (94.7%) and multiciliation (23.3%) compared to WT cells (38.7% and 0.0%, respectively) (Figure 4D). As described below, myc-Rootletin expression reverses ciliary and centrosomal defects in Rootletin KO cells. Thus, results from Rootletin null MEFs phenocopy those in ELMOD2 KO lines with increased rates of ciliation (89.1%) and multiciliation (24.4%) that are also very similar in magnitude to those seen in ELMOD2 KO lines. Another similarity noted is that centrin staining is evident in Rootletin KO cilia, even without ciliobrevin treatment (Figure S8A). Yet, Rootletin nulls are distinct from ELMOD2 nulls in that they do not have cold sensitive microtubules, multinucleation, or evidence of cytokinesis defects (Turn et al., 2020). A more subtle difference is seen in multiciliated cells, as while ELMOD2 nulls have a wide range in the number of cilia per cell (typically 2-3, but can be 9 or more), multiciliated Rootletin KO cells almost never have more than two cilia. Finally, in contrast to ELMOD2 KO cells, Smo is recruited to cilia in response to treatment with SHH conditioned medium in Rootletin KO cells as well as in WT MEFs (Figure 4E). Thus, ELMOD2 and Rootletin KO cells share commonalities in defects in centrosome cohesion, increased ciliation, and multiciliation, yet there are also clear differences in aspects of these phenotypes.

We initially screened for Rootletin KO lines by immunofluorescence and noticed that a few cell lines displayed much stronger staining of Rootletin rather than loss (Figure 4B, right most panel). One of these lines (called G1, #21) was preserved and analyzed further by DNA sequencing. This clone had frameshifting mutations on both alleles. One allele is a 1bp insertion, differing from the 1bp insertion found in Clone G1, #31 only in the base inserted. The other allele is a 2bp insertion that was unique to that clone (Figure S6). Both exon skipping and use of alternative initiating methionines are now well-established consequences of CRISPR/Cas9 genome editing (Smits et al., 2019). Based on such data, we propose that this clone uses the first methionine after the insertion, Met240, to generate an N-terminal truncated protein that we term Rootletin^Δ239^. This results in a protein that is 239 residues shorter than the full-length mouse Rootletin, or a total of 1770 residues and thus ~26 kDa shorter. An immunoblot of total cell lysate from WT cells, probed with the Rootletin antibody, shows the major band at ~240 kDa, and this band is absent or replaced by a stronger band at ~215 kDa, consistent with our prediction of an N-terminal truncation mutant being generated (Figure 4, see Figure S8 for original blot). To further test our prediction, we generated a vector designed to express myc-Rootletin^Δ239^ under control of the CMV promoter and used it to express the truncated protein in WT and mutant cells. As expected, the exogenously expressed myc-Rootletin^Δ239^ appeared as a band of slightly stronger intensity and faster electrophoretic mobility than the full length, ~240 kDa band and was indistinguishable in electrophoretic mobility from the strongly staining band in the clone we label Rootletin^Δ239^ (Figure 4, Figure S9). Rootletin staining of Rootletin^Δ239^ cells revealed the presence of rootlets, though they are abnormally large, bright, and fibrous compared to rootlets in WT MEFs (Fig. 4B). Rootletin^Δ239^ cells displayed no change in the percentage of cells with centrosome separation (Figure 4C), consistent with retention of this function of Rootletin. However, they have severely reduced percentages of ciliation compared to WT cells, with an average of only 6.5% of cells being ciliated (compared to 38.7% in WT cells) after serum starvation. These cells also show no sign of multiciliation (Figure 4D). Together, these data indicate that, like ELMOD2, Rootletin expression is associated with suppression of ciliation while its absence results in increased ciliation and multiciliation.

### ELMOD2 localization is disrupted in Rootletin KO cells

Because ELMOD2 and Rootletin KO lines share similarities in phenotypes and the proteins extensively co-localize at rootlets and centrosomes, we predict that they act in a shared biochemical pathway. Both proteins are reorganized/recruited to basal bodies early in ciliogenesis, and ELMOD2 deletion caused disrupted rootlet organization. Thus, we next examined whether deletion of Rootletin would alter the localization/organization of ELMOD2 at the basal body. Rootletin null cells were serum starved for 24hr, fixed, and stained for ELMOD2, acetylated tubulin, and γ-tubulin. As described above, ELMOD2 typically localizes to basal bodies and rootlets in WT cells. As expected, Rootletin KO results in loss of ELMOD2 localization at rootlets (as Rootletin-positive rootlets are absent), though these cells retain ELMOD2 staining at basal bodies. Surprisingly, there is an acquisition of strong ciliary localization of ELMOD2 in cells lacking Rootletin (Figure 4F-G). This ciliary staining of ELMOD2 is punctate and distributes preferentially to the distal tip in a subpopulation of cilia. In Rootletin^Δ239^ cells, in which rootlets are retained and even magnified, ELMOD2 staining is strongly rootlet-associated, even more so than in WT rootlets, consistent with Rootletin playing a role in ELMOD2 recruitment (Figure 4F-G). We examined ELMOD2 localization at other sites (*e.g.* midbodies and mitochondria) and found no changes in ELMOD2 at those sites in Rootletin KO cells (Figure S10). This suggests that Rootletin specifically plays a role in ELMOD2 localization to cilia/rootlets but not to other cellular compartments. Together, these data reveal that Rootletin and ELMOD2 localization at basal bodies and rootlets are co-dependent and that loss of either results in very similar phenotypic consequences at that site.

### Rootletin over-expression rescues ELMOD2 null phenotypes

As described above, we have used expression of myc-tagged proteins to protect against off-target effects of CRISPR and shown that vectors directing expression of ELMOD2-myc or Rootletin-myc reverse phenotypes resulting from KO of either gene. Because of the close physical and functional relationships observed for the two proteins at the base of cilia, we tested whether expression of either protein could reverse the phenotypes resulting from deletion of the other gene. Transfected cells were fixed and stained for different combinations of myc, acetylated tubulin, γ-tubulin, and Rootletin to assess ciliary, centrosomal, and rootlet phenotypes, with empty vector (pCDNA3.1) serving as negative control. The elevated rates of ciliation seen in Rootletin KO cells (89.0%) were returned to near WT levels (37.8%) upon expression of Rootletin-myc (as described above) but stayed elevated in cells expressing ELMOD2-myc (89.6%) (Figure 5A). In contrast, expression of Rootletin-myc was sufficient to reverse the increased ciliation seen in ELMOD2 nulls (36.9%) (Figure 5A). Similarly, Rootletin-myc expression reversed the effects of ELMOD2 deletion on centrosome separation, while ELMOD2-myc fails to alter this phenotype in Rootletin KO cells (Figure 5B). As expected, empty vector had no effect (Figure 5A, B). The strong suppression of ciliation seen in Rootletin^Δ239^ was unaffected by expression of either myc-tagged ELMOD2 (1.0%) or Rootletin (1.5%) (Figure 5A, B). On the other hand, myc-Rootletin reversed both increased ciliation (WT: 27.3%; ELMOD2 KO: 36.9%; Rootletin KO: 37.8%; Rootletin^Δ239^: 1.5%) and increased centrosome separation defects (WT: 14.8%; ELMOD2 KO: 21.4%; Rootletin KO: 17.1%; Rootletin^Δ239^: 12.5%) in both ELMOD2 KO and Rootletin KO cells (Figure 6A-B). Expression of myc-Rootletin fails to reverse the phenotypes in Rootletin^Δ239^ expressing cells, consistent with these phenotypes resulting from an already increased level of Rootletin activity. These results further support a close functional link between ELMOD2 and Rootletin and even suggests that ELMOD2 may play a regulatory role in recruiting Rootletin to basal bodies that can be overcome in its absence by excess Rootletin. We interpret these results as consistent with a model in which the two proteins act in a common pathway, with Rootletin acting downstream of ELMOD2.

**Figure 5:**
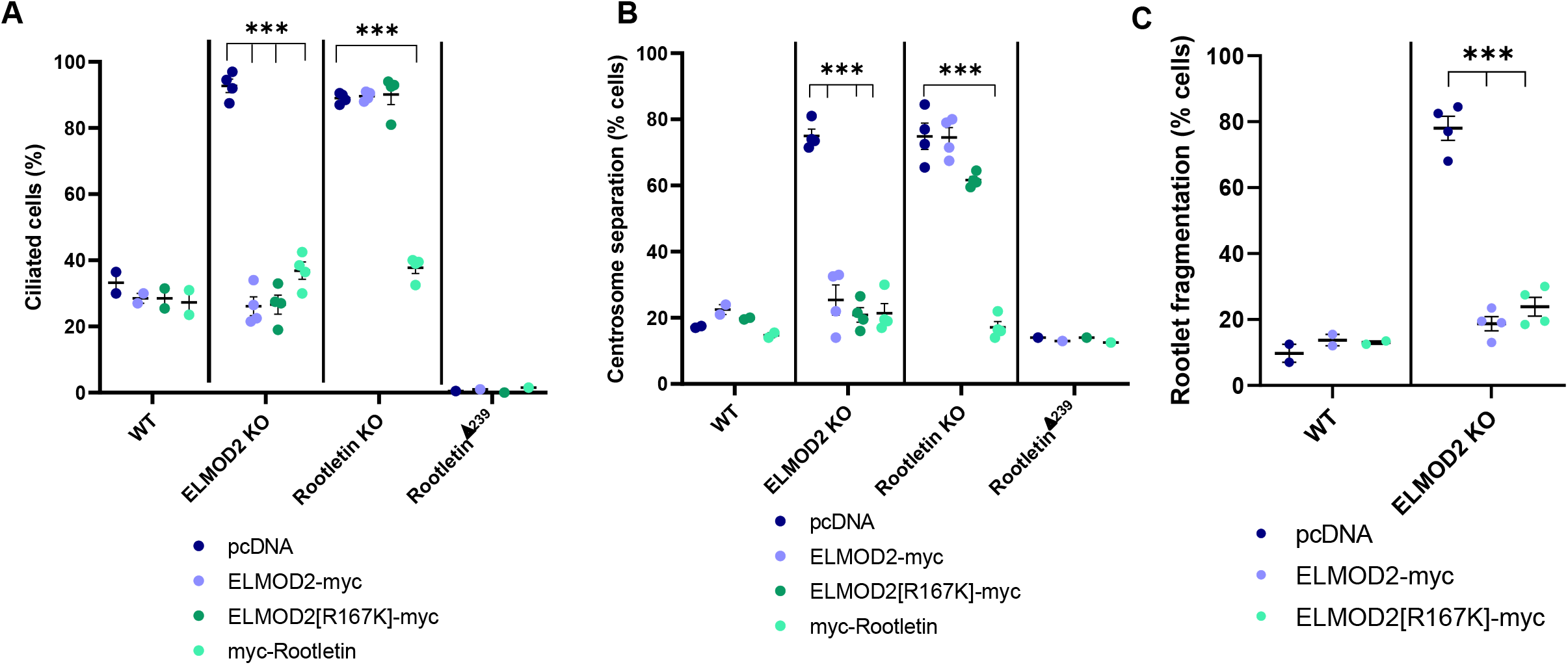
ELMOD2-myc and ELMOD2[R167K]-myc rescue ciliation and centrosomal cohesion defects in ELMOD2 KO but not Rootletin KO cells. Cell lines (2 WT, 4 ELMOD2 KO, 4 Rootletin KO, and Rootletin^Δ239^) were transfected with either empty vector, or plasmids directing expression of ELMOD2-myc or ELMOD2[R167K]-myc before being re-plated onto coverslips, serum starved, fixed with ice cold methanol, and stained for Rootletin, acetylated tubulin, and γ-tubulin. Cells were scored in duplicate for either **(A)** % ciliation, **(B)** centrosome separation (centrosomes >2μm apart), or **(C)** rootlet fragmentation, with 100 cells scored per replicate. The averages of individual lines were plotted as individual points in leafed scatterplots. Error bars indicate SEM. Statistical significance was assessed using One-Way ANOVA, comparing each of the test groups to WT. In cases where multiple conditions show the same statistically significant change compared to WT, a bracket pointing to each line showing that change is indicated; ***=p<0.0001.

**Figure 6:**
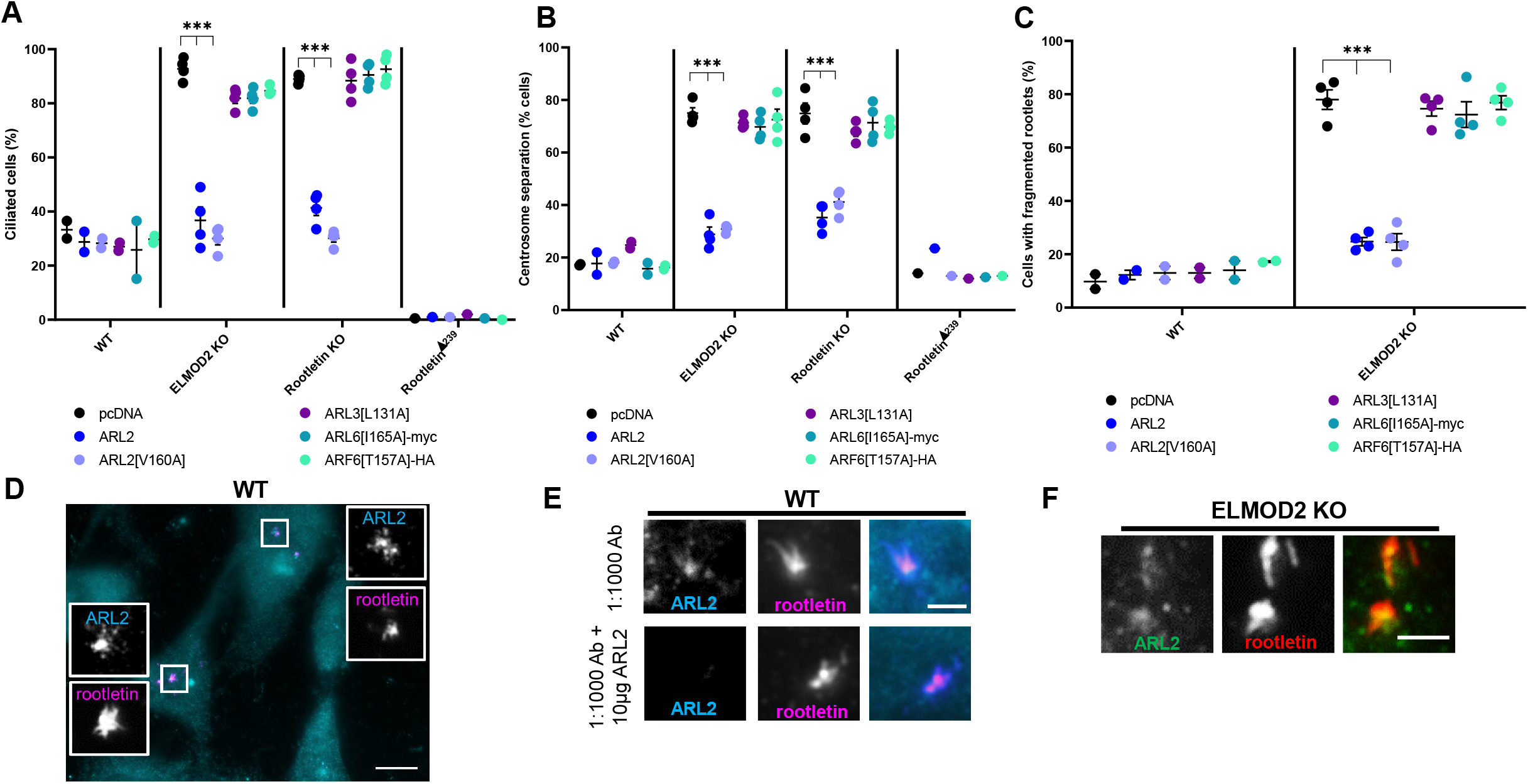
ARL2 and ARL2[V160A] reverse the increased ciliation, rootlet fragmentation, and centrosome separation defects seen in ELMOD2 and Rootletin KO cells. Cell lines (2 WT, 4 ELMOD2 KO, 4 Rootletin KO, and Rootletin^Δ239^ mutant) were transfected with the following constructs: pcDNA (empty vector control), ARL2, ARL2[V160A], ARL3[L131A], ARL6[I165A]-myc, or ARF6[T157A]-HA before being serum starved for 24 hours. Cells were then stained for Rootletin, acetylated tubulin, and γ-tubulin, then scored in duplicate for either **(A)** % of cells with at least one cilium, **(B)** centrosome separation (centrosomes >2μm apart), or **(C)** rootlet fragmentation, with 100 cells scored per replicate. The averages of individual lines were plotted as individual points in leafed scatterplots. Error bars indicate SEM. Statistical significance was assessed using One-Way ANOVA, comparing each of the test groups to WT. In cases where multiple conditions show the same statistically significant change compared to WT, a bracket pointing to each condition is shown; ***=p<0.0001. **(D)** WT cells show ARL2 co-localization with Rootletin. A representative image is shown using widefield microscopy at 100x magnification. Scale = 10μm. **(E)** ARL2 staining at rootlets is lost with antigen competition. Images were collected using the same conditions described in (**A**), except that in the lower panel the primary antibody was incubated with 10 μg purified recombinant human ARL2 prior to use in cell staining. **(F)** ARL2 localization to rootlets is maintained in ELMOD2 KO cells. Images were collected using the same conditions as described in (D), except that ELMOD2 KO cells rather than WT MEFs were used.

### ELMOD2 functions in cilia are at least in part independent of its GAP activity

With the history of ARF family GAPs acting as both GAPs (to provide temporal regulation of signaling) and effectors (components in the pathway that propagate the signal) (East and Kahn, 2011; Schiavon et al., 2019; Zhang et al., 1998), we asked whether GAP activity is required for the actions of ELMOD2 at cilia or rootlets. We used the point mutation that has previously been shown to result in loss of GAP activity as a result of the mutation of the “arginine finger” in the GAP domain (East et al., 2012; Schiavon et al., 2019). We transfected WT and ELMOD2 KO cells with either empty vector control, ELMOD2-myc, or the GAP-activity dead ELMOD2[R167K]-myc. Cells were then serum starved for 24 hours, and ciliated cells were scored using myc staining to identify transfected cells, along with acetylated tubulin to mark cilia. Ciliation rates in WT cells (28.5%) were unchanged after recombinant protein expression (Figure 5A) while in ELMOD2 KO lines the elevated rate of ciliation (89.1%) was reversed upon transient expression of ELMOD2-myc (26.1%), comparable to what we observed previously using lentiviral transduction (Figure 1B). The GAP dead mutant yielded reversal that was indistinguishable from that of ELMOD2-myc (Figure 5A). Both ELMOD2-myc and ELMOD2[R167K]-myc expression each also reversed the centrosome separation present in ELMOD2 KO lines (Figure 5B). Thus, the actions of ELMOD2 in regulating ciliation rates and centrosome cohesion are independent of its GAP activity and suggest that it is acting in a GTPase pathway to propagate the downstream effects.

We also asked whether ELMOD2 actions at rootlets require GAP activity. We used the same plasmids to transfect WT and ELMOD2 KO cells and scored for rootlet fragmentation (Figure 5C). Again, neither ELMOD2-myc nor ELMOD2[R167K]-myc expression had an effect on this phenotype of WT cells (empty vector: 9.8%; ELMOD2-myc: 13.8%; ELMOD2[R167K]-myc: 13.0%) (Figure 5C). Yet expression of either ELMOD2-myc (18.8%) or ELMOD2[R167K]-myc (23.9%) reversed the elevated rootlet fragmentation in ELMOD2 KO cells (78.0%) to levels comparable to those seen in WT cells (~12%) (Figure 6C). Together, these data suggest that ELMOD2 does not rely on GAP activity to mediate ciliary or rootlet functions. Note that the GAP-dead [R167K] mutant retains binding affinity to activated ARF family GTPases. Therefore, we predict that ELMOD2 is acting as a downstream effector rather than as a terminator of GTPase signaling.

### ARL2 can rescue ciliary and centrosomal defects in Rootletin and ELMOD2 KO cells and specifically localizes to rootlets

Because ELMODs are single domain proteins that bind the activated conformation of a number of ARF family GTPases, we predict that an ARF family GTPase is also involved in the basal body/rootlet actions of ELMOD2. A number of ARF family GTPases have been linked to ciliary functions including ARL2, ARL3, ARL6, and ARL13B (Fisher et al., 2020). We showed previously that ELMOD2 acts with ARL2 in mitochondria and tubulin assembly, and with ARF6 at recycling endosomes and Flemming bodies (Turn et al., 2020). Therefore, we asked if increased activity of any of these GTPases can influence the actions of ELMOD2 at cilia and centrosomes. We transiently expressed WT or activated mutants of ARL2, ARL3, ARL6, and ARF6 and scored effects on ciliation rates, centrosome separation, and rootlet fragmentation, as in previous sections. Although by far the most commonly exploited activating mutation in regulatory GTPases is that of changing the glutamine in the G-3 motif to leucine (Q to L), we have found that expression of such mutants are often quite toxic to cells, making analyses of their cellular actions difficult (Turn et al., 2020; Zhou et al., 2006). As a result, we have increasingly relied upon “fast cycling” point mutants, in which a conserved residue in the G-5 motif is mutated, resulting in decreased affinity for GDP with retention of GTP binding (Aspenstrom, 2018; Santy, 2002a). This allows for ready binding of the activating ligand, GTP, and inactivation by GAPs, thus preventing toxicity that may result from excess activity that cannot turn over. The fast cycling, activating mutations tested are ARL2[V160A], ARL3[L131A]-myc, ARF6[T157A]-HA, ARL6[I165A]-myc, and ARL13B[V168A]-myc. Of the recombinant proteins assessed in these assays, only ARL2 and ARL2[V160A] reversed elevated ciliation, defective centrosome cohesion, and rootlet fragmentation in ELMOD2 KO lines (Figure 6A-C). None of the GTPase constructs tested could reverse the strong block in ciliogenesis in Rootletin^Δ239^ cells. It is interesting to note that increasing cellular ARL2 activity, but not ELMOD2, reversed both phenotypes seen in Rootletin KO cells (ciliation rates and centrosome separation) and also reversed rootlet fragmentation in ELMOD2 KO cells. Thus, while we believe that these results support a role for ARL2 acting in a pathway with ELMOD2 and Rootletin, they also leave open the possibility of ARL2 acting with Rootletin independently of ELMOD2.

Previous work in our lab and others uncovered specific localization and roles for ARL2 at centrosomes, mediating microtubule nucleation as well as tubulin folding (Francis et al., 2017a; Francis et al., 2017b; Zhou et al., 2006). Other studies have also implicated ARL2 in stabilizing photoreceptor cilia (Wright et al., 2018) and in regulating ciliary length (Davidson et al., 2013). We immunostained for ARL2 using multiple fixation and permeabilization conditions but did not detect evidence of ciliary staining. Instead, and in addition to centrosomal or basal body staining, we observed that ARL2 co-localizes with Rootletin staining at rootlets after cold methanol fixation and that this staining is lost in response to antigen competition using purified, bacterially expressed ARL2 (Figure 6D-E). ARL2 also localizes to cilia in Rootletin KO cells suggesting that, like ELMOD2, rootlets can influence staining of ARL2 in cilia (Figure S7B). Despite its presence at centrosomes and in cilia, ARL2’s closest paralog, ARL3, shows no such rootlet staining (Figure S11). Thus, we found a high degree of specificity among ARF family members in the ability of ARL2 to restore basal body and Rootletin functionalities in the absence of ELMOD2 (or Rootletin). In ELMOD2 KO cells, ARL2 still localizes to rootlets, suggesting that ELMOD2 is not required for ARL2’s recruitment to rootlets (Figure 6F). We found specific localization of ARL2 to basal bodies and rootlets, consistent with ARL2 acting in concert with ELMOD2 and Rootletin in the regulation of ciliogenesis and centrosome separation.

### ELMOD2 localizes to the base of the connecting cilium and the axoneme of the outer segment in human and mouse retinal cells, while ARL2 is found at the ciliary rootlet

With the unexpected observations regarding functions of ELMOD2 and ARL2 and localizations at cilia in MEFs, we sought to determine if they are also found at cilia in better studied models of ciliary function. Photoreceptor cells (rods and cones) in the retina are perhaps the most commonly used model for ciliary signaling due to their large cilia, resulting from their specialized role in phototransduction. Mouse and human retinas were processed for immunofluorescence imaging, as described under Materials and Methods and previously (Davidson et al., 2013; Trojan et al., 2008). Staining of centrin (to mark the connecting cilium as well as the mother (basal body) and daughter centrioles of photoreceptor cells) and ELMOD2 are shown in Figure 7A-F. In both human and mouse retinal tissues, ELMOD2 is found at the base of the connecting cilium. This compartment is equivalent to the transition zone (TZ) of primary cilia that connects the outer and inner segments of the photoreceptor cells. Higher magnification of the periciliary region reveal the localization of ELMOD2 right between the basal body and the adjacent daughter centriole (Figure 7C, E, F), which are connected by rootlet-like fibers (Yang et al. 2002). The ciliary rootlet originates from here to project into the inner segment and is weakly stained for ELMOD2 in human photoreceptor cells (Figure 7D,E). Interestingly, ELMOD2 is also found above the connecting cilium along the highly modified axoneme of the photoreceptor outer segment (arrows in Figure 7C, E) (Birtel et al., 2017; May-Simera et al., 2017). Here, at the basal part of the outer segment, photo-sensitive disk membranes are assembled via a highly regulated process driven by actin polymerization which involves molecules specific for photoreceptor cilia (Corral-Serrano et al., 2020; Salinas et al., 2017; Spencer et al., 2019). There is growing evidence that this region resembles the ciliary tip in highly modified photoreceptor cilia (Corral-Serrano et al., 2020).

**Figure 7:**
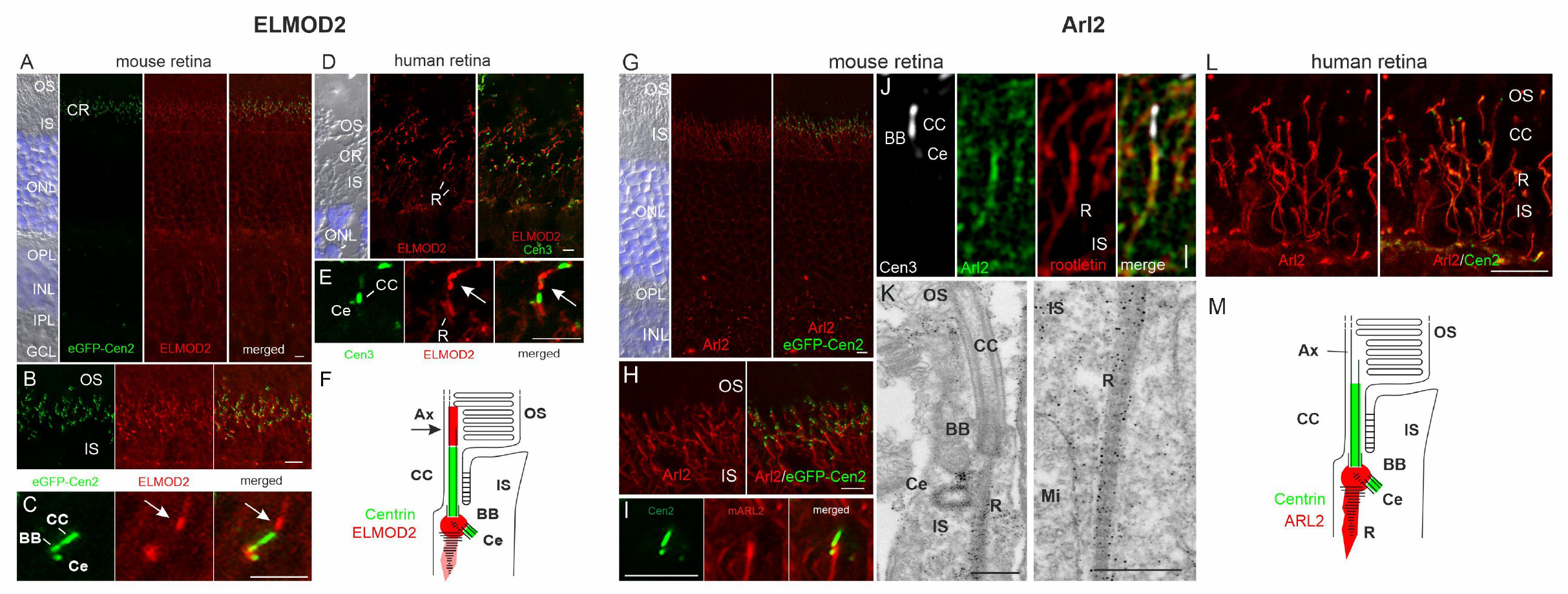
ELMOD2 and ARL2 display specific localizations to either the periciliary region and the base of the axoneme or the basal body and the ciliary rootlet, respectively, in human and mouse retinal photoreceptor cells. Human and mouse retinas harvested from patient donors or WT (bl6) and transgenic eGFP-CETN2 mice were cryosectioned, immunolabeled and analyzed with either a deconvolution microscope, a confocal laser scanning microscope or transmission electron microscopy, as described under Materials and Methods. **(A,B)** In the retina of transgenic eGFP-CETN2 mice, immunolabeling revealed prominent immunofluorescence of ELMOD2 in the ciliary region (CR) of photoreceptor cell layer at the junction between the outer segment (OS) and the inner segment (IS). The other retina a layers, blue DAPI-stained outer and inner nuclear layer (ONL, INL), outer and inner plexiform layer (OPL, IPL) and ganglion cell layer (GCL) did not show substantial staining. **(C)** Higher magnification imaging revealed ELMOD2 in the periciliary region between GFP-Centrin signal at the basal body (BB) and the adjacent centriole (Ce) and in continuum of the connecting cilium (CC), the axoneme of the photoreceptor OS base (arrow). **(D,E)** Co-immunolabeling using ELMOD2 and Centrin 3 antibodies validated the periciliary region and axoneme (arrow) localization of ELMOD2, but also indicate weak staining of ciliary rootlets (R) in human retinal photoreceptor cells. **(F)** Scheme of ELMOD2 localization in photoreceptor cells. ELMOD2 localizes to the base of the axoneme (Ax), the periciliary region, and to the ciliary rootlet (in human). **(G,H,I)** In transgenic eGFP-CETN2 mice, immunostaining indicated the localization of ARL2 in the periciliary region and at the ciliary rootlets (R) of the photoreceptor CC. **(J)** Furthermore, coimmunolabeling of ARL2 and rootletin in mouse retinas revealed co-localization of both proteins. **(K)** Immunoelectron microscopic pre-embedding labeling revealed ARL2 localization in ciliary rootlets (R) and at the adjacent centriole (Ce) and basal body (BB) in the periciliary region and thereby confirmed the co-localization of ARL2 and rootletin. Mi, mitochondrion; **(L)** In human retinas immunostaining of ARL2 and Centrin 2 validated the rootlet localization as it was previously shown in the mouse retina. **(M)** Scheme of ARL2 localization in photoreceptor cells. ARL2 is localized to periciliary region (at BB and Ce) and the ciliary rootlet (R). Scale in G-J and L: 5 μm; in K: 400 nm.

Because ARL2 has not previously been shown to localize to rootlets, and because rootlets are typically not studied in MEFs, we again turned to the far better characterized retinal cells to assess its localization in mouse and human retinal cells (Figure 7G-M). A previous study explored effects of excess ARL2 activity in photoreceptor cells using a rod-specific promoter to express the dominant activated ARL2[Q70L]-HA-FLAG mutant and reported changes in ciliary morphology and function (Wright et al., 2018). ARL2 localizes to the rootlet itself in photoreceptor cells in both mice and humans as evidenced by its co-localization with Rootletin by immunofluorescence (Figure 7J, L) and by immunoelectron microscopy, in which rootlet staining is clearly evident (Figure 7K). The latter also highlights staining of ARL2 at the mother centriole but is not observed in the outer segment, even at the base where ELMOD2 was seen. Thus, ELMOD2 and ARL2 each display specific localization to centriole, rootlets, the base of the axoneme, and the axoneme in the case of ELMOD2. While the overlap in staining of ELMOD2 and ARL2 in the periciliary region, where the ciliary rootlet originates, provides a likely site of action in a shared pathway, the distinct localizations leave open the possibility of each acting separately from the other.

### ELMOD2 and Rootletin regulate the ciliogenesis pathway by preventing spurious licensing through CP110 release

Ciliogenesis is a multi-step process and ciliary defects can arise as a result of lesions in any of a number of pathways including, but not limited to, ciliary licensing, centrosomal docking, formation and maintenance of the TZ, and intraflagellar transport (IFT). Despite the lack of detailed molecular models of each step in this process, a number of studies have identified roles for key proteins that also serve as markers of specific steps in ciliogenesis. We screened a number of such markers to assess their integrity in ELMOD2 KO lines, 24 hr after serum starvation (Figure S12). IFT88 is a component of the IFT-B complex, active in anterograde ciliary traffic. It displays bright punctate staining throughout the length of the cilium, often with preferential staining at the base and tip (Figure S12C). No differences were evident in the staining of IFT88 between WT and ELMOD2 KO lines. Two TZ markers, NPHP4 and Cep290, each localize strongly to the TZ at the proximal end of the cilium, and again no differences were evident in their staining profiles (Figure S12A-B). Thus, despite the evidence described above that loss of ELMOD2 results in apparent defects in ciliary protein import, export, and/or retention, we found no gross changes in the transition zone or localization of IFT based on the use of these few diagnostic markers of each.

In marked contrast to the unaltered appearance of markers of the TZ and IFT, we observed spurious localization of proteins involved in ciliary activation (aka “licensing”). Early in ciliary assembly, the basal body is primed to dock to the plasma membrane and to project a cilium by the regulated and sequential recruitment and later dissociation of a number of proteins. Key regulators of this process include Cep164 (a distal appendage protein that facilitates basal body docking to the membrane) (Schmidt et al., 2012), TTBK2 (a kinase that binds to Cep164 at distal appendages and phosphorylates key players that facilitate CP110 release) (Goetz et al., 2012; Lo et al., 2019), and CP110 (a centriolar capping protein that must be removed to allow docking of the ciliary vesicle which will eventually fuse with the plasma membrane to initiate growth of the cilium at the cell surface) (Spektor et al., 2007). In a WT cell, a centriole that is ready to assemble a cilium stains positive for Cep164 and TTBK2 but negative for CP110. To ensure that only one cilium is generated per cell, any other centrosome(s) in a cell should show the opposite staining pattern so that they cannot recruit ciliary vesicles or dock at the plasma membrane. This predicted outcome was confirmed in WT MEFs, which display at most one centrosome per cell that stains positive for Cep164 and TTBK2 but negative for CP110.

While only 2.0% of WT cells showed >1 centrosome that was positive for Cep164, it was a far more common occurrence in ELMOD2 KO cells (33.8%) (Figure 8A, S13A). TTBK2 followed a similar trend, as 37.4% of ELMOD2 KO cells had >1 centrosome positive for TTBK2, compared to only 1.3% of WT cells (Figure 8B, S12B). On the other hand, loss of the capping protein CP110 (>1 centrosome negative for CP110) was observed in 8.5% of WT cells vs 42.3% of ELMOD2 KO cells (Figure 8C, S13C). This spurious activation of centrioles into basal bodies is reversed in ELMOD2 MEFs expressing ELMOD2-myc (Figure 8). We interpret these data as evidence that ELMOD2 normally plays a role in preventing spurious ciliary activation and that its loss leads to misregulation of the licensing events. Specifically, ELMOD2 acts between the Cep44 and CP110-dependent steps of the pathway that ensure that only one centriole becomes a basal body per cell, resulting in increased ciliation and multiciliation.

**Figure 8:**
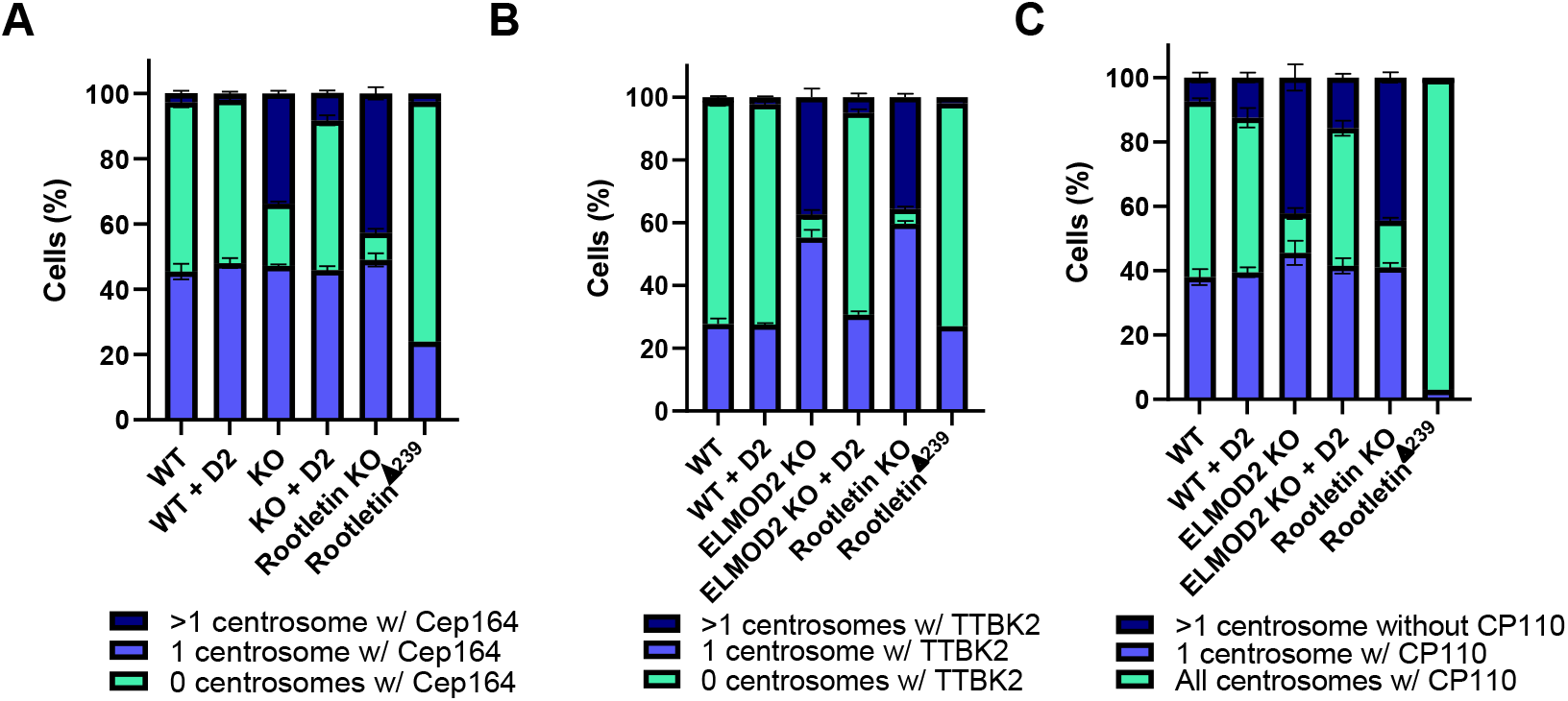
ELMOD2 KO causes misregulation of markers of different steps in ciliogenesis. **(A)** Loss of ELMOD2 or Rootletin leads to increased Cep164 recruitment. Cells (2 WT, 4 ELMOD2 KO, 4 ELMOD2 KO + ELMOD2-myc, 4 Rootletin KO, and 1 Rootletin^Δ239^) were serum starved and scored for changes in Cep164 localization, using γ-tubulin to mark cilia, as described under Materials and Methods. Cells were scored in duplicate and binned as either having 0, 1, or >1 centrosome positive for Cep164. Data were plotted in a stacked bar graph, and error bars indicate SEM. **(B)** TTBK2 is increased at centrosomes in ELMOD2 and Rootletin KO cells. The same conditions as shown for (A) were used to monitor changes in TTBK2 recruitment, except cells were costained with both γ-tubulin and acetylated tubulin to track both centrosomes and cilia, and cells were fixed for only 5 minutes. **(C)** Deletion of either ELMOD2 or Rootletin leads to increased CP110-negative centrosomes, even cells with >1 centrosome being negative for CP110. The same conditions as shown for (A) were used to determine if CP110 localization to centrosomes changes in ELMOD2 KO cells.

We performed comparable analyses of markers of ciliogenesis in stained Rootletin KO and Rootletin^Δ239^ cells. Like in ELMOD2 KOs, loss of Rootletin led to an increased percentage of cells with >1 centrosome being positive for Cep164 (42.7%) compared to WT cells (3.5%) (Figure 8A). The Rootletin KO lines also yielded similar increases in TTBK2 to the ELMOD2 nulls (Figure 8B), as well as a similar extent in loss in CP110 staining (44.5% of Rootletin KO versus 7.3% of WT) (Figure 8C). Thus, deletion of either ELMOD2 or Rootletin causes spurious licensing of ciliogenesis that results in increased rates of ciliation and likely contribute to multiciliation.

As described above, Rootletin^Δ239^ cells demonstrate reduced ciliation rates after 24 hr of serum starvation. Therefore, we predicted that we would see a reduction in Cep164 and TTBK2 and an increase in CP110 localization to centrosomes compared to WT. Interestingly, the Rootletin^Δ239^ line had only slightly decreased levels of Cep164 localization at centrosomes (46.3% of cells with ≥1 centrosome positive for Cep164 in WT vs 26.5% in Rootletin^Δ239^) (Figure 8A). The same was true for TTBK2 (29.0% in WT vs 29.0% in Rootletin^Δ239^) (Figure 8B). In marked contrast, the CP110 cap is overwhelmingly retained in Rootletin^Δ239^ cells, as 96.5% versus 58.7% of WT cells have all their centrosomes positive for CP110 (Figure 8C), consistent with their far lower percentage (6.5%) of ciliated cells after serum starvation. We interpret these data as evidence that cellular actions of ELMOD2 and Rootletin (and by extension rootlets) include regulating ciliary licensing, specifically acting proximal to Cep164 and basal body docking and uncapping of distal appendages via release of CP110 release (Figure 9, Figure S13).

**Figure 9:**
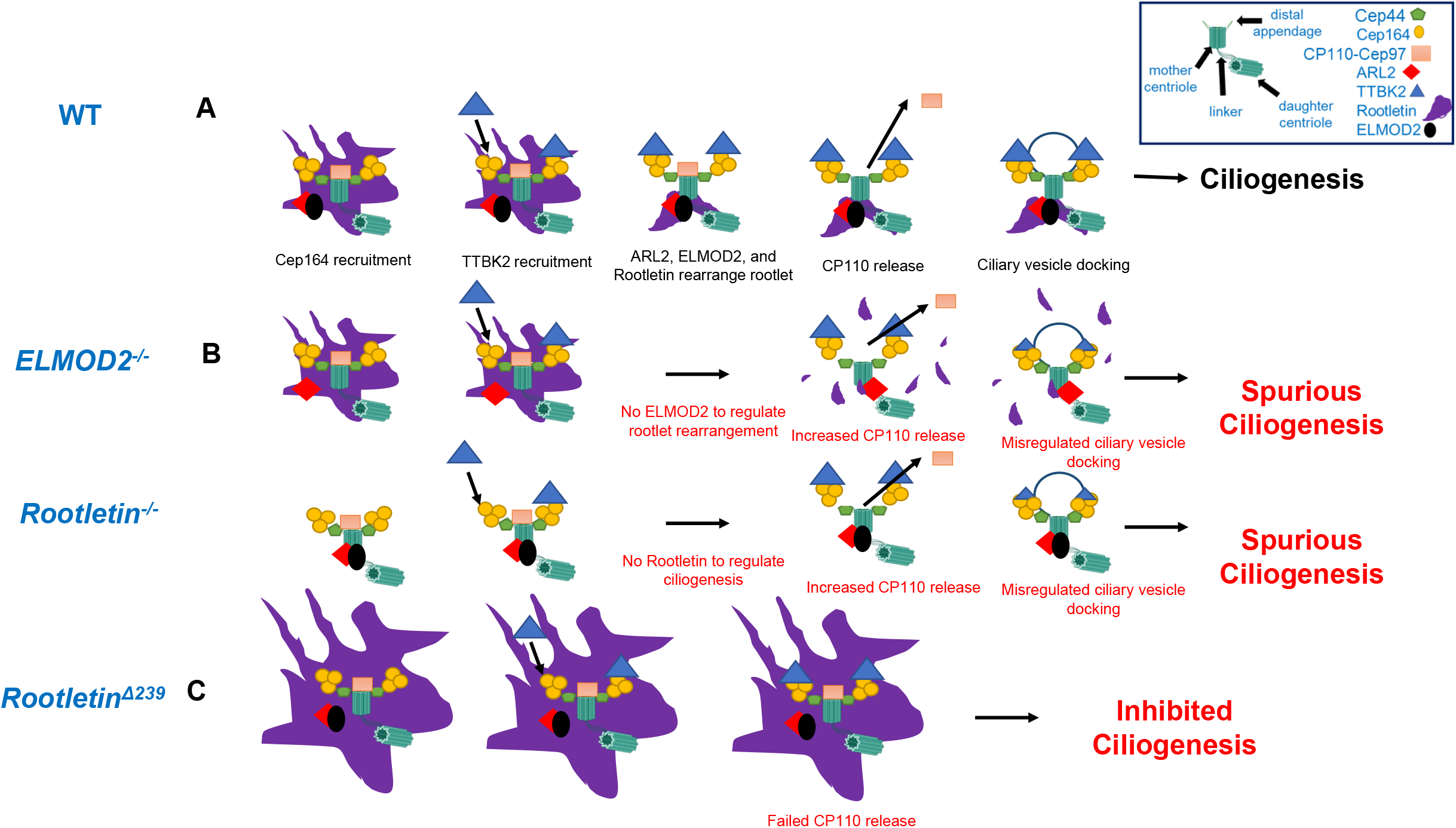
ELMOD2, ARL2, and Rootletin work together to prevent spurious ciliogenesis. **(A)** In WT cells responding to serum starvation or signal to ciliate, ELMOD2 is shown bound to ARL2 at basal bodies and with Cep44, the CP110-Cep97 complex, all surrounded by rootlets and the recent recruitment of Cep164 highlighted at the start (left) of the pathway. The presence of Cep164 recruits TTBK2 directly and subsequently rootlets are re-organized as licensing continues. The release of CP110-Cep97 allows for ciliary vesicle docking and ciliogenesis to occur. We propose that ELMOD2 and Rootletin act early in suppress ciliogenesis to regulate licensing, by preventing spurious CP110-Cep97 complex release. **(B)** In the absence of ELMOD2 or Rootletin we see increased incidence of Cep164 and TTBK2 recruitment, loss of rootletin organization around the basal body in ELMOD2 KO or simply no rootlets in the Rootletin KO, and increased CP110 release, resulting in consequent increased ciliation and multiciliation. **(C)** The Rootletin^Δ239^ line shows increased localization of Rootletin, ARL2, and ELMOD2 at centrosomes and strongly reduced ciliation compared to WT. These cells have slightly reduced Cep164 and TTBK2 recruitment and strong retention of CP110, resulting in inhibition in ciliogenesis progression.

## Discussion

This study has provided new insight into the cellular actions of ELMOD2, close functional links to rootlets at basal bodies, and roles of each in ciliogenesis. We found that loss of ELMOD2 causes increased ciliation, multiciliation, abnormal ciliary morphology, altered recruitment of a subset of ciliary membrane receptors and centrin, loss of rootlet attachment to centrosomes/basal bodies, and loss of centrosome cohesion. These phenotypes were reversed upon expression of WT or activated ARL2, linking this GTPase and ELMOD2 in the same pathway(s). These findings are also consistent with the original purification of ELMOD2 as an ARL2 GAP, despite displaying activity in in vitro GAP assays for some other ARF family members (Bowzard et al., 2007; Ivanova et al., 2014). The connection between ELMOD2 and rootlets was greatly strengthened by the observations that deletion of Rootletin resulted in many of the same phenotypes seen in ELMOD2 KO cells and that increased expression of Rootletin was sufficient to reverse these phenotypes in ELMOD2 KO lines. These results prompted us to (re)examine the localization of ELMOD2 and ARL2 in MEFs and photoreceptor cells, where we found novel localizations for each, including overlap in the periciliary region and ARL2 at rootlets. We propose that ELMOD2 and Rootletin inhibit uncontrolled, spurious ciliogenesis by regulating rootlet morphology, thereby affecting the ability of mother centrioles to develop into basal bodies and to carry out the regulated recruitment and release of key factors, *e.g.*, CP110, that drive ciliogenesis. We summarize these observations with our proposed model for the actions of ELMOD2, ARL2, and Rootletin in ciliary licensing in Figure 9.

Given our previous data showing roles for ELMOD2 at centrosomes and in microtubule nucleation and stability, we were initially concerned that the changes in ciliary functions may be indirect, resulting from alterations in microtubules or axonemes or from centrosome amplification described previously in ELMOD2 null MEFs (Turn et al., 2020). This concern was perhaps increased by the finding that increasing ARL2 activity is sufficient to rescue each. While we do not believe that we can completely separate these actions of ELMOD2 in cells, we currently conclude that they are distinct actions. Perhaps the strongest single piece of evidence arguing for distinct actions of ELMOD2 in microtubule and ciliary functions is the finding that expression of the GAP dead mutant (ELMOD2[R167K]) was equally active as WT to restore ciliary integrity (Figure 5) but failed to reverse the defects at microtubules/centrosomes (Turn et al., 2020). Thus, the GAP activity of ELMOD2 is required at centrosomes/microtubules but not at cilia. In addition, we showed that expression of activated ARF6 reversed the centrosome amplification phenotype in ELMOD2 KO MEFs, consistent with it being a consequence of failed abscission (Turn et al., 2020). Expression of the same ARF6 construct, though, had no effect on ciliary phenotypes (Figure 6). There are numerous examples in which centrosome amplification is not accompanied by increased ciliation (Coelho et al., 2015; Jonassen et al., 2008; Lee et al., 2011; Zhou et al., 2014). Also, while loss of microtubule nucleation/stability as well as stalling in cytokinesis might be expected to *decrease* ciliation, we found *increased* ciliation in ELMOD2 null lines. In addition, the multiple commonalities found between deletion of ELMOD2 and Rootletin strongly support the conclusion that they act together, yet previous studies of rootlets consistently find no clear links to microtubules. We also found no differences in microtubule density or sensitivity to nocodazole in Rootletin KO MEFs compared to WT, suggesting clear distinctions between ELMOD2 acting on microtubules and Rootletin at basal bodies and rootlets. We screened Rootletin KO cells for defects in cell cycle via flow cytometry for DNA content, and we observed no evidence of the polyploidy that is seen in ELMOD2 KO cells (Figure S14). Neither Rootletin KO nor ELMOD2 KO cells show changes in total protein content or in the expression of markers such as ARL3 or centrin, making it seem unlikely that simply a change in protein expression due to polyploidy could explain the mislocalization of proteins to cilia (Figure S15A-B). Finally, the fact that multiciliation (Figure 1F) and spurious licensing of ciliogenesis (Figure S16) is present in cells with both abnormal and normal cellular morphology (e.g., one nucleus, 1-2 centrosomes) all points to the unlikelihood that cell cycle defects alone can explain the spurious ciliation observed in Rootletin and ELMOD2 KO cells. Perhaps its unique position as a regulator of ciliogenesis, microtubule anchoring, and cytokinesis allows ELMOD2 to mediate all three pathways, allowing communication that ensures proper regulation of cell cycling and signaling (East and Kahn, 2011; Francis et al., 2016; Turn et al., 2020).

The molecular details of neither the early steps in ciliogenesis (licensing) nor the later steps that are critical to the formation of a functional cilium are completely understood, despite significant advances in recent years (Chen et al., 2020; Gigante and Caspary, 2020; Nachury, 2018; Nachury and Mick, 2019; Satir, 2017). We used the previous identification of specific steps or components in licensing as markers in the pathway to identify likely sites of action for ELMOD2 and Rootletin. An incomplete model of this pathway is shown in Figure 9. Cep164 recruits to distal appendages early in this process (Graser et al., 2007a; Schmidt et al., 2012) to allow for the docking of the serine-threonine kinase, TTBK2. TTBK2 is required for the removal of the CP110-Cep97 protein complex from the basal body, allowing ciliary vesicle docking (Goetz et al., 2012). Recent work has revealed that TTBK2 phosphorylates M-phase phosphoprotein 9 (MPP9) (Huang et al., 2018), a factor that recruits to the distal appendages to mediate CP110-Cep97 docking. MPP9 phosphorylation by TTBK2 promotes its degradation, leading to the release of CP110-Cep97 complex. Cep83 has also been identified as a substrate for TTBK2, as its phosphorylation by TTBK2 is critical for Cep83 to drive pre-ciliary vesicle docking to distal appendages, CP110 release, and the formation of the ciliary vesicle (Lo et al., 2019).

Of the markers we tested, the first step in the pathway that displayed differences in the KO lines is increased staining of Cep164 and TTBK2 in both ELMOD2 and Rootletin nulls. Because Cep44 is unaltered in either ELMOD2 or Rootletin KO, while Cep164 and TTBK2 are increased at centrioles, we propose that ELMOD2 acts in the regulated recruitment of rootlets/Rootletin to centrioles. Consistent with this conclusion is the observation that rootlet attachment at basal bodies (and centrosomes) is lost in cells that lack ELMOD2. The normal, physical linkages between centrioles and rootlets may present a physical barrier to proteins being recruited or removed. Thus, the loss of rootlet attachment to the basal body also may help explain the increased recruitment of Cep164 and TTBK2 (Figure 9). Alternatively, the rootlets linked to centrioles may serve as a scaffold for recruitment of regulators of ciliogenesis. In ELMOD2 KO cells, the increases in both Cep164 and TTBK2 are predicted to cause the spurious release of the CP110 complex that caps and prevents ciliation.

We further propose that, like ELMOD2, Rootletin plays a role in inhibiting spurious ciliation by preventing the recruitment of Cep164 and TTBK2 and later release of CP110 (Figure 9). Rootletin KO cells also display increased staining of Cep164 and TTBK2 at centrosomes and decreased CP110, leading to increased ciliogenesis. There is some debate in the field over the role that rootlets play in ciliogenesis, as Nigg’s group (Graser et al., 2007a) reported no change in ciliogenesis upon knockdown of Rootletin in hTERT-RPE1 cells, though the Morrison group reported a *reduction* in ciliogenesis in hTERT-RPE1 cells upon knockdown of Rootletin (Conroy et al., 2012). Furthermore, several groups depleted C-NAP1 (a known binder of the ciliary rootlet that is important for its proper localization to centrosomes) and observed normal cilium assembly (Flanagan et al., 2017; Mazo et al., 2016; Panic et al., 2015). Our data add to this debate, as we propose that Rootletin normally *inhibits* ciliogenesis in MEFs, and thus its loss caused increased ciliation. Differences in findings between labs could result from cell type differences (MEFs vs RPE cells), as much of the other work was either performed in multiciliated cells (in which it may be difficult to see *increased* ciliation) or in the highly specialized retinal photoreceptor cells in which additional control mechanisms may be in place. Alternatively, differences may result from the approaches used (*i.e*., knockout vs knockdown), as the latter results in incomplete loss of protein. A Rootletin null mouse was generated (Yang et al., 2005) by inserting a neo cassette between exon 3 and 4 to splice into the transcript rather than removing all or part of the gene. They no longer detected rootlets in these cells, but protein is still detectable by Western (the type of tissue and age of samples used in the immunoblot were not reported), which would suggest that they do not necessarily have a functional null mouse line. This could potentially explain discrepancies between our findings and those reported by the Li Lab, though as noted, other explanations also exist. Data in our lab from knockout of another protein with no prior evidence linking it to cilia or centrosomes resulted in strong changes in rootletin staining, with losses at centrosomes, yet retention of centrosome cohesion. Thus, we believe that there is more to be learned about the connections between rootlets, centrosomes, and ciliation that should prove to be important and will help resolve these apparent differences.

Results from testing in our Rootletin^Δ239^ cell line further support our model that Rootletin is acting proximal to the release of CP110 to restrain ciliary licensing. It is not clear at this time whether the phenotypes observed in this line result from the increased expression, the loss of key N-terminal residues, or both. These cells display increased CP110 retention at both centrosomes, despite the presence of Cep164 and its near normal increase in response to serum starvation. Yet, this cell line displays a profound *loss* of ciliation. The discovery of Rootletin^Δ239^ highlights once again both advantages and cautions in the use of CRISPR/Cas9 genome editing, including the use of internal methionines to initiate translation (Smits et al., 2019). We believe the resulting frame shift leads to the use of Met240 to initiate translation, resulting in the N-terminal truncation mutant that lacks the first 239 amino acids. The faster migrating band in the immunoblot from these cells (Figure 4A) and its co-migration with the protein made in cells expressing our custom-made truncation mutant Rootletin^Δ239^ (Figure S6D, E) are consistent with this interpretation.

How are these processes regulated? In short, we don’t know. We believe the evidence strongly supports a role for the regulatory GTPase ARL2 acting with ELMOD2. We speculate that TTBK2 may phosphorylate Rootletin and, in so doing, regulate its actions or half-life. TTBK2 is a relatively understudied kinase but is predicted to have an unusual preference for a phosphotyrosine at the +2 position relative to its site of serine/threonine phosphorylation (Bouskila et al., 2011). ELMOD2 contains no (S/T)XY sequence, but Rootletin does: serine 155 is followed by tyrosine at residue 157, making it a potential substrate for TTBK2. Interestingly, the Rootletin^Δ239^ mutant is missing these residues, and the truncated protein is expressed to much higher levels than the wild type protein. Thus, the N-terminus of Rootletin may include sites of regulated protein-protein interaction and protein half-life. Results from our studies of cells expressing Rootletin^Δ239^ are also consistent with those from the Li lab and others that have used the N-terminal truncation protein, termed R234, that lacks 500 residues from the N-terminus (Akiyama et al., 2017; Yang et al., 2006; Yang et al., 2002), yet retains the ability to form rootlets. Recent structural studies confirm the importance of more C-terminal portions of Rootletin, specifically coiled-coil domain 3, to oligomerization and centrosome binding (Ko et al., 2020), though these previous studies have not investigated protein half-life or roles in ciliation. Future studies focusing on functions and binding partners of the N-terminal portion of Rootletin are likely to provide new insight into the mechanisms by which rootletin regulates ciliary function.

Rescue experiments provide evidence linking ARL2 to ELMOD2 and Rootletin in ciliogenesis as well as the conclusion that ELMOD2 is acting as an effector in this pathway, and not as a GAP (Figures 6, 7). ARL2, ARL2[V160A], and myc-Rootletin each rescue ciliary and centrosome cohesion defects (and Rootletin defects in the case of ELMOD2 KOs) in both Rootletin and ELMOD2 KO cells (Figures 6, 7). This effect is specific to ARL2 as neither its closest structural paralog, ARL3, nor functionally linked paralogs like ARL6 and ARF6, rescue these defects. Furthermore, expression of ELMOD2-myc or the GAP dead point mutant each reverse ciliary, rootlet, and centrosome cohesion defects seen in ELMOD2 KO, showing quite clearly that GAP activity is not required and, rather, the role of ELMOD2 as an effector (East and Kahn, 2011; Zhang et al., 2003; Zhang et al., 1998). While ELMOD2 expression does not rescue ciliary phenotypes in Rootletin KO cells, the converse is true: *i.e*., increased expression of Rootletin reverses ELMOD2 KO defects. The former is perhaps not surprising as deletion of Rootletin results in the loss of an important cytoskeletal polymer with clear roles in centrosome cohesion. Still, we believe this finding also suggests that Rootletin acts very close to ELMOD2 and likely immediately downstream of it in this pathway, suggesting that ELMOD2 plays a non-essential, but critical, role in regulating Rootletin morphology to direct ciliogenesis. Perhaps ELMOD2 facilitates the proper tethering and bundling of rootlets at centrioles to accommodate cellular needs (*e.g*., ciliation, mitosis), consistent with both being present at centrioles in photoreceptor cells (Figure 7).

Given the importance of Rootletin to form rootlets, it was surprising to find that activated ARL2 (ARL2[V160A]) and ARL2 each reverse ciliary defects in Rootletin KO lines. One possible explanation may be that ARL2 has multiple actions including one downstream of Rootletin. Perhaps in WT cells rootlets allow for anchorage and recruitment of ARL2 so that it may regulate ciliogenesis, and therefore loss of the rootlet leads to the dilution of ARL2’s signal and prevents it from inhibiting ciliation. Overexpression of ARL2 or the activated mutant would override the system, allowing for greater access of ARL2 to the site and therefore restoring inhibition of ciliogenesis. This would be consistent with general GTPase biology, as the tight regulation of an ARF’s localization in time and space is pivotal for the proper coordination of cell functions (D’Souza-Schorey and Chavrier, 2006; Fisher et al., 2020; Jackson and Casanova, 2000; Kahn, 2009; Kahn et al., 2005; Mizuno-Yamasaki et al., 2012; Nie et al., 2003; Sztul et al., 2019). Alternatively, ARL2 may be acting through components of rootlets rather than Rootletin directly, such as C-Nap1 or Cep68 (Vlijm et al., 2018). This might explain the reversal of ciliary defects and centrosome separation despite the absence of Rootletin. Clearly more work is required to better understand and interpret these results.

Roles for ELMOD2 and ARL2 in centrosome cohesion are also implicated by our results (Figures 3G, 5B). Previous work demonstrated roles for rootlets to surround centrosomes and help maintain their cohesion throughout the cell cycle with loss during cell division, and Rootletin siRNA caused spurious centrosome separation (Bahe et al., 2005; Graser et al., 2007b; Hossain et al., 2020; Meraldi and Nigg, 2001; Vlijm et al., 2018; Yang et al., 2006). Failure to regulate centrosome cohesion can lead to cell cycle defects, such as centrosome amplification or aneuploidy. ARL2 has also been linked to aspects of breast cancer biology (Beghin et al., 2009; Beghin et al., 2008), and we showed that ELMOD2 KO MEFs take on properties of cell transformation, including loss of contact inhibition and anchorage independent growth (Turn et al., 2020).

Our studies increase both the number of sites and pathways at which ARL2 and ELMOD2 have been shown to act. The ARF family includes at least four members with known roles in ciliary biology, including ARL2, ARL3, ARL6, and ARL13B. Previous studies revealed that ELMOD2 has *in vitro* GAP activity toward ARL2, ARL3, ARF1, ARF3, and ARF6, but not ARL13B (Ivanova et al., 2014). Thus, despite the specificity observed in our rescue experiments (Figures 5, 6), we cannot exclude the possibility that ELMOD2 is acting with one or more of these other GTPases to regulate aspects of ciliary biology. It is interesting to note that ELMOD2 and ARL2 do not completely co-localize at cilia or with rootlets in MEFs or in photoreceptor cells. In MEFs, there are some rootlet branches that do not stain for ELMOD2 or ARL2, and there are some cases in which it looks like ARL2 or ELMOD2 create their own rootlet-like projections that are negative for Rootletin. ARL2, ARL3, and RP2 (which can act as a GAP for ARL2 and ARL3) are implicated in protein traffic to cilia, particularly that of prenylated (acting with PDE6δ) or N-myristoylated (acting with UNC119) protein cargos (Evans et al., 2010; Hanke-Gogokhia et al., 2016; Jaiswal et al., 2016; Veltel et al., 2008; Wright et al., 2011; Wright et al., 2016). Clean dissection of the actions of each of these GTPases and their GAPs/effectors clearly requires additional studies. Such research is certain to further our currently incomplete understanding of the pathways involved, as well as likely uncover links to other aspects of cell biology.

In ELMOD2 KO MEFs, ARL2 co-localizes with Rootletin at centrosomes, though not to Rootletin positive structures at other sites in the cell. In Rootletin KO MEFs, ARL2 and ELMOD2 each localize to basal bodies, though they no longer form rootlet-like projections. We interpret these data as evidence that while these three proteins act in shared pathways, they are unlikely to act in or as a single complex, as suggested for other components in ciliary licensing or ciliation (Humbert et al., 2012; Jin et al., 2010; Kobayashi et al., 2014). Further adding to the complexity in defining roles, there appears to be some level of tissue or cell type specificity to their localizations. ARL2 localizes to the rootlet of retinal cells and basal body, while ELMOD2 localizes to the axoneme, rootlet, and base of the connecting cilium. In contrast, ARL2 appears almost exclusively along the length of the cilium in bronchial cells, while ELMOD2 shows strong staining at the ciliary tip as well as faint co-localization with ciliary rootlets (Figure S17). It is noteworthy that the localization of ELMOD2 at the ciliary tip of motile cilia is in line with the localization of ELMOD2 to the outer segment base of photoreceptor cells (Figure 7C,D), which corresponds to the ciliary tip of the highly modified cilium there (Corral-Serrano et al., 2020). A related concern is that ARF family GTPases bind membranes only transiently and in a highly regulated fashion, and their localization at sites like the Golgi are often missed (e.g., see Human Protein Atlas data), despite the overwhelming evidence of their actions there (D’Souza-Schorey and Chavrier, 2006; Donaldson and Honda, 2005; Kahn, 2009; Liu et al., 2010; Liu et al., 2014). These findings suggest the possibility, if not likelihood, of tissue specificity to ELMOD2 and ARL2 localization and action, perhaps changing in response to different cellular/ciliary requirements. Although there has been no published follow up to the earlier linkage between the ELMOD2 gene and idiopathic pulmonary fibrosis (Hodgson et al., 2006; Pulkkinen et al., 2010), we note the importance of multiciliated cells in this condition and perhaps an argument to look deeper into these questions.

We also observed alterations in localization of ciliary signaling proteins in ELMOD2 KO MEFs, though these defects may be secondary to ciliogenesis structural defects, *e.g*., resulting from dissociation or loss of rootlets/Rootletin. Previous reports (Mahjoub, 2013; Mahjoub and Stearns, 2012) that increasing PLK4 induces centrosome amplification that leads to the production of multiple primary cilia were interpreted with respect to the dilution effect of multiciliation on ciliary signaling components. ELMOD2 KO caused decreased recruitment of multiple signaling proteins, including SSTR3, Smo, GPR161 (Figure S2), and ACIII and decreased SHH pathway output (Figure 2). Therefore, receptor traffic and signaling defects could be secondary to multiciliation or altered ciliary morphology. However, monociliated cells also display reduced recruitment of ciliary receptors, so simple dilution of components into multiple cilia appears unlikely to fully explain the changes that we observed. Furthermore, if it was simply a matter of diluted signal, Rootletin KO cells would also show disrupted Smo accumulation, but neither mono-nor bi-ciliated cells display alterations in Smo staining. How the loss of ELMOD2 alters the localization of ciliary signaling proteins remains unclear and is a fascinating direction to explore in the future. Recently the Nachury group reported a novel role for ubiquitin in marking GPCRs that are ready for removal from the cilium by the BBSome (Shinde et al., 2020). ELMOD2 could be regulating signaling through any number of pathways, and one interesting future direction would be to explore ELMOD2’s functions in this capacity. We also see a number of other proteins that localize to cilia upon loss of ELMOD2 or Rootletin, including centrin, ELMOD2, and ARL2. It is quite possible that these proteins normally cycle in and out of cilia but, under normal conditions, they do not accumulate and thus escape detection by immunofluorescence imaging. Previous studies have shown that centrin localizes to motile cilia (Yamamoto et al., 2008) and to the TZ of photoreceptor cilia (Wolfrum, 1995), but there is only one report showing centrin at the base of primary cilia, but not in the ciliary axoneme (Trojan et al., 2008). Another possibility to explain the ciliary localization of these proteins is that IFT or transition zone are defective in ELMOD2 KO cells but not Rootletin KO cells. We did not observe defects in transition zone or IFT based on gross immunofluorescence analysis (Figure S11), but future studies should be directed to examination of IFT traffic and TZ integrity, perhaps using super resolution or electron microscopy. Together, these data suggest that though ELMOD2 and Rootletin work together for some ciliary functions, ELMOD2 may have discrete functions at other ciliary compartments.

Finally, we also observed both defects in the axoneme (acetylated tubulin) and components of the ciliary membrane (ARL13B), noting branching and splaying of cilia that is virtually nonexistent in WT primary cilia. Though we do not have clear answers as to the source of these abnormal morphologies, there are a number of possibilities. For example, the increase in spurious ciliogenesis may lead to the skipping of checkpoints that are critical for ensuring proper ciliary morphology. Other reports of cells with increased ciliation have also noted abnormal ciliary morphologies, so are unlikely to be specific to the proteins or cells used in our study (Yasar et al., 2017).

Together, this study has uncovered multiple new functions and localizations for ELMOD2, ARL2, and Rootletin in ciliary and rootlet function. There are still many more questions than answers concerning how these proteins act together and independently to direct so many different ciliary and other essential cellular functions. With the additional information provided herein, we hope to incentivize future work into understanding the signaling events that mediate the communication between cilia, cell cycle, microtubule dynamics, and other essential cellular functions.

## Materials and Methods

### Reagents, antibodies, plasmids

The commercially obtained antibodies and dilutions used in imaging herein include those directed towards: γ-tubulin (1:5000) (Sigma; T6557), γ-tubulin (1:5000) (Abcam; ab11317), centrin clone 20H5 (1:1000) (Sigma; 04-1624), myc (1:1000) (Invitrogen; R950-25), HA (1:1000) (Covance; MMS-101P), acetylated tubulin (1:2000) (Sigma; T6793-2ML), ARL13B (1:500) (Proteintech; 10083-118), ARL13B (1:500) (Abcam; ab136648), Gli3 (1:1000) (R&D Systems; AF3690), Cep164 (1:100) (Santa Cruz; sc-515403), CP110 (1:100) (VWR; 76045-052), IFT88 (1:500) (VWR; 10088-640), NPHP4 (1:100) (VWR; 10091-250), Cep290 (1:100) (VWR; 10084-648), Rootletin (1:500) (Millipore-Sigma; ABN1686), TTBK2 (1:100) (Sigma; HPA018113-100UL), Cep44 (1:100) (Proteintech; 10084-652). We initially used an SSTR3 antibody from Santa Cruz; however, this antibody has been discontinued, so later studies of SSTR3 used expression of the tagged protein. Rabbit polyclonal antibodies against the following human proteins were generated by the Kahn lab and have been previously characterized: ARL2 (Sharer and Kahn, 1999; Sharer et al., 2002), ARL3 (Cavenagh et al., 1994), and ELMOD2 (Newman et al., 2014). We are grateful for the generous gifts of other antibodies: ARF6 polyclonal antibody from Jim Casanova (Univ. of Virginia) and Smoothened from Kathryn Anderson (Sloan Kettering).

As described in our previous manuscripts (Turn et al., 2020), the CRISPR-Cas9 system used to generate the null lines involved use of a plasmid obtained from Addgene (pSpCas9(BB)-2A-Puro (PX459) V2.0 (#62988)). Plasmids directing expression of human ARL2, ARL2[Q70L], ELMOD2-myc, or ELMOD2[R167K]-myc/his in pcDNA3.1 were described previously (Bowzard et al., 2007; East et al., 2012; Zhou et al., 2006). Jim Casanova provided us with plasmids used for transient expression of ARF6-HA, ARF6[Q71L]-HA, or ARF6[T157A]-HA (Altschuler et al., 1999; Santy, 2002b). All fast-cycling point mutants were generated in pcDNA3.1 using site-directed mutagenesis and confirmed by DNA sequencing. Max Nachury (UCSF) provided us with SSTR3-GFP (Ye et al., 2013). Ciliobrevin D was purchased from Sigma (#250401). Human bronchial cells (NH BE009) were provided by Mike Koval and were grown on transwell plates.

### Cell Culture

All CRISPR/Cas9 KO lines were grown under the same conditions, maintaining cells at low passage (below passage 10). WT MEFs were purchased from ATCC (CRL-2991), and all KO lines were generated from this original line. Cells were grown in DMEM (Fisher; 11965092) with 10% FBS (Atlanta Biologicals; S11150) and 2 mM glutamine at 37°C, 5% CO_2_ and were screened at least monthly for mycoplasma contamination. Serum starvation was used to induce ciliation and involved growth of cells in DMEM with 0.5% FBS and 2 mM glutamine for 24hrs. No antibiotics were used in the routine maintenance of cells. We treat replicates of individual lines on separate days as technical replicates, and we treat the average of these technical replicates for each line as biological replicates.

### Generation of CRISPR null lines

ELMOD2 and Rootletin KO lines were generated using CRISPR-Cas9, as described in our previous publication (Turn et al., 2020). In brief, four guides per gene targeted (20 nt long) were designed using Benchling software. Guides were cloned into pSpCas9(BB)-2A-Puro (PX459) V2.0 using BbsI restriction sites and then transfected into WT MEFs at a 1:3 ratio of DNA (4 μg) to Lipofectamine 2000 (12 μg) for 4 hours in Opti-MEM medium (Fisher; 31985070). Cells were then replated into 10 cm dishes and grown overnight. The next day, puromycin selection (3 μg/ml, Sigma #P8833) was initiated and lasted for a total of 4 days, to enrich for transfected cells. Cells were then grown to near confluence in our regular culture medium (DMEM + 10% FBS). Cloning was performed by plating into 2 96-well plates at ~3 cells/well. Clones were screened for frameshifting mutations in both strands using DNA sequencing, with primers flanking the predicted cut site. At least two clones from each of two guides were generated to protect against off-target effects.

Rootletin guide RNAs targeted exons 6-9, to induce frame shifting mutations with the goal of making non-functional protein products. Screening was performed initially by immunofluorescence of MEF lines cloned by limiting dilution, followed by DNA sequencing surrounding the targeted exon to identify frame shifting mutations in both alleles. We generated 5 predicted KO lines from three different guide RNAs (see Fig. S5A-B).

### Lentiviral Transduction

The mouse ELMOD2-myc open reading frame was cloned into the pFUGW vector using EcoRI and BamHI sites, to generate lentivirus by Emory’s Viral Vector Core, as previously described (Turn et al., 2020). Cells were incubated with virus for 48 hrs and then fresh medium was exchanged, cells were grown up and frozen. Transduction efficiency was determined to be ~70-90%, based upon myc staining. Because of this high transduction efficiency, use of these lines in “rescue” experiments involved counting all cells in the population, rather than only those expressing ELMOD2. Therefore, because 10-30% of the cells scored in such experiments are not expressing ELMOD2-myc, less than complete rescue is expected, as routinely observed.

### Transfection of MEFs

As described in our previous manuscript (Turn et al., 2020), Lipofectamine 2000 proved toxic to several of our cell lines (particularly ELMOD2 KOs). Therefore, we used polyethyleneimine (PEI) for later transfections, due to its reduced cellular toxicity. The day before transfection, cells were plated at 70% density in 6-well dishes. These cells were transfected with a 1:3 ratio of DNA to PEI for 24 hours in medium containing 2% FBS in DMEM. Unless otherwise stated, 4 μg of DNA, 12 μg PEI, and 100 μL of serum-free medium per reaction were combined in an Eppendorf tube, vortexed, and incubated for 20 minutes at room temperature before being added dropwise to the sample. Cells were replated onto Matrigel coated coverslips at the appropriate density, serum starved for 24hrs, and processed for immunofluorescence imaging. For rescue experiments, JetOPTIMUS transfection reagent was used (VWR; 76299-634). Cells were plated at 70% density in 6-well dishes before transfection with 4μg of DNA with 4μL JetOptimus for 24 hours, in DMEM with 2% FBS.

### Western blotting

Cells were plated at approximately 90% density and harvested the next day. Cell pellets were resuspended into 1x Laemmli sample buffer, heated at 95°C for 5 minutes, and spun down to remove insoluble, before resolving proteins in a 7.5% polyacrylamide SDS gel. Proteins were transferred to nitrocellulose filters overnight at 20V. Membranes were stained for Ponceau to check for equal loading, blocked with filtered 5% blotto in PBST (BioRad; 1706404) for 1hr at room temperature, and stained with primary antibody against Rootletin (1:500 dilution) overnight at 4°C. Membranes were washed 3 x 10 min in PBST, incubated with HRP-anti-chicken secondary antibody for 1hr at room temperature and developed using enhanced chemiluminescence.

### Immunofluorescence

After plating cells onto Matrigel coated coverslips and performing other drug/serum starvation treatments as needed, we used different fixation and permeabilization conditions, depending upon the primary antibody used.

### PFA fixation

For ciliary markers (except for γ-tubulin) such as ARL13B, Gli3, or IFT88, cells were fixed with pre-warmed (37°C) 4% PFA for 15 minutes on the bench top. Cells were then rinsed 4x with PBS before being permeabilized with 0.1% Triton X-100 for 10 minutes and blocked with 1% BSA in PBS for 1 hr at room temperature. Primary antibody was diluted into blocking solution and incubated overnight at 4°C, before rinsing 4x with PBS, application of secondary antibody (1:500) for 1hr at room temperature in the dark, 4x wash with PBS, Hoechst (1:5000) applied, and coverslips were mounted on slides using MOWIOL + PPD (9:1 ratio).

### 5 min methanol fixation

For centrosomal markers (centrin, γ-tubulin, ARL2, ELMOD2, TTBK2) and Rootletin, cells on coverslips were fixed for 5 min at −20°C in ice-cold methanol. Coverslips were then rinsed 4x with PBS, blocked and incubated with antibodies as described above. TTBK2 was unique in that lower background staining required 3% BSA or 10% FBS for blocking.

### 10 min methanol fixation

Some centrosomal and ciliary markers (*e.g*. Cep164, CP110, Cep290, NPHP4, Rootletin) required longer fixation; cells were fixed at −20°C with ice-cold methanol for 10 min before washing 3X with PBS at room temperature. Cells were blocked with 10% FBS for 30 min, primary antibodies added in 10% FBS and incubated overnight at 4°C. Coverslips were rinsed 3x, 5 minutes each, with PBS and then incubated in secondary for 1hr at RT in the dark. Coverslips were rinsed and mounted onto slides as described above.

### Ciliobrevin treatment

Cells were plated onto Matrigel-coated coverslips and serum-starved for 24 hours as described above. Ciliobrevin (30 μM) (Sigma-Aldrich; 250401-10MG) was added to the cultures for 1hr at 37°C before fixing cells with 4% PFA for 15 minutes at 37°C. Samples were processed as described above, except PBS containing 0.1% Tween detergent (PBST) was used for washes to remove autofluorescent residue left by the drug.

### qPCR of SHH pathway targets

Cellular responses to SHH included measuring transcriptional changes in *Gli1* mRNAs, as previously described (Mariani et al., 2016). Cells were incubated with 0.5% FBS SHH-conditioned or 0.5% FBS control medium for 48 hours, with a media replacement after the first 24-hour period. Cells were harvested, and RNA extracted using the Qiagen RNeasy Kit with QIAshredder homogenizer columns according to manufacturer’s protocols. RNA (200 ng) was used to generate cDNAs using BioRad iScript Reverse Transcription supermix. The following primers were used during qPCR to detect transcript levels:

### Pold3 (housekeeping gene)

F: 5’ - ACGCTTGACAGGAGGGGGCT - 3’

R: 5’ - AGGAGAAAAGCAGGGGCAAGCG-3’

### Gli1

F: 5’-CTTCACCCTGCCATGAAACT-3’;

R: 5’-TCCAGCTGAGTGTTGTCCAG-3’

In brief, the cDNA was combined with primers and Bio-Rad SsoAdvanced Universal SYBR Supermix according to manufacturer’s protocols (1725270). Samples were run on a Bio-Rad CFX96 Touch Real-Time PCR Detection System, and data were analyzed using Bio-Rad CFX Manager 3.1. The following program conditions were used: 95°C for 5 min; 45 cycles of 95°C for 15 s; 57°C for 30 s. Reactions were performed in technical duplicate on three biological replicates. Data were then analyzed by the ΔΔCT method and normalized to control WT levels for each transcriptional target (Livak and Schmittgen, 2001). Data were subjected to a two-way ANOVA followed by Sidak’s posthoc test for comparisons within cell types between control and SHH-treated conditions in Prism software (GraphPad). Significant differences were determined by a p-value <0.05. Graphed data are presented as mean (± standard deviation) fold change over untreated WT transcript levels.

### Analyzing DNA content

Unsynchronized cells at ~70% confluence were harvested in 50mL conical tubes, washed with ice-cold 1xPBS, and fixed with ice-cold 70% ethanol and stored at 4°C. The day of flow cytometry, cells were spun down, resuspended and washed in phosphate citrate buffer (0.1M citric acid in 1xPBS, pH 7.8), spun down, and treated with RNase (100 μg/mL; Sigma; R5125) for 15 minutes at room temperature. The cells were then stained with propidium iodide (50 μg/mL; Sigma, P4170) for at least 45 minutes at room temperature. Cells were run on a FACSymphony A3 at low speed, and 10,000 events were collected per condition using the same gating conditions and centering the G1 peak at 50K. Data were processed in FloJo Software.

### Microscopy

All fixed immunofluorescence experiments were performed using Matrigel (BD Bioscience) coated 18 mm glass coverslips (#1.5, Fisher Scientific; 12-545-81) prepared in the lab. Samples were visualized using confocal (Olympus FV1000 microscope and Olympus Fluoview v1.7 software; 100x magnification (1.45 NA, Oil); 405, 488, 543, and 635 laser lines) and widefield microscopes (Olympus IX81 microscope and Slidebook software; 100x magnification (UPIanFI, 1.30 NA Oil). For the majority of the images shown (as indicated in the figure legends), confocal microscopy was used to collect z-stacks (0.37μm steps) using image processing to ensure that the full cilium/basal body/rootlet are visible. The same acquisition settings (gain, laser power, offset, *etc*.) were used for every sample within each experiment. FIJI imaging software was used to process the z-projections, and the same brightness, contrast, cropping, and other processing settings were used across the experimental test group to ensure the accuracy of comparisons. Retinal sections were analyzed with a Leica DM6000B deconvolution microscope (Leica Microsystems, Bensheim, Germany) or a Leica SP8 laser scanning confocal microscope (Leica Microsystems, Bensheim, Germany) and images were processed with Adobe Photoshop CS (Adobe Systems, San Jose, CA, USA).

### Super resolution microscopy

3D-SIM images of cilia were collected using a Nikon super-resolution microscope (N-SIM) at 100x magnification (1.49 NA, Oil) using a 488-laser line and an EMCCD-Andor iXon3 DU-897E-CS0-#BV camera. Data were acquired and processed using Nikon Elements v5.0.2 software. Widefield images along with raw SIM data were collected for every cilium studied. Nikon Elements was used to reconstruct images collected via SIM, and reconstruction parameters were adjusted as needed to prevent the introduction of artifacts that did not coincide with the original widefield image.

g-STED images were collected using a Leica gSTED 3× microscope at 100x magnification (NA 1.4, oil). Z-stack projections were collected for each cell with a 0.22 μm step-size, and images were collected using 488 and 561 laser lines for excitation and 592 and 660 laser lines for depletion. Images were acquired and processed using Leica X software. Confocal, gSTED, and deconvolved gSTED images were collected.

### Live cell Imaging

Wild-type cells were transfected with plasmid directing expression of GFP-Rootletin, using the JetOPTIMUS protocol described above, and cells were replated onto 35 mm MatTek dishes (#P35GC-1.5-14-C). The next day, cells were imaged using a BioTek Lionheart FX widefield microscope at 20x magnification (NA.45) using the 488 channel and phase-contrast, maintained with 5% CO_2_ at 37°C throughout the imaging window. Images were collected every 5 minutes for 12 hours immediately after initiation of serum starvation (DMEM+0.5% FBS). Videos were processed via Lionheart imaging software.

### Animals

Wildtype (bl6) and transgenic eGFP-CETN2 mice (Higginbotham et al., 2004) were kept on a 12-hour light-dark schedule at 22°C, with free access to food and water. Animal health was monitored on a regular basis, and all other procedures complied with the German Law on Animal Protection and the Institute for Laboratory Animal Research Guide for the Care and Use of Laboratory Animals, 2011.

### Human tissue

The human donor eye tissue applied in the present study we obtained 11.5 h *post mortem* from a female donor (# 252-09), 65 years of age without any underlying health conditions, from the Department of Ophthalmology, University Medical Center Mainz, Germany. The guidelines to the declaration of Helsinki (http://www.wma.net/en/30publications/10policies/b3/) were followed.

### Immunohistochemistry of retinal sections

Human and mouse retinae were dissected from enucleated eye balls and cryofixed in melting isopentane and cryosectioned as previously described (Karlstetter et al., 2014; Wolfrum, 1991). Cryosections (10 μm thick) were placed on poly-L-lysine-precoated coverslips and incubated with 0.01% Tween 20 in PBS for 20 min. After washing with PBS, sections were covered with blocking solution (0.5% cold-water fish gelatin plus 0.1% ovalbumin in PBS) and incubated for at least 30 min followed by an overnight incubation with primary antibodies at 4°C in blocking solution (Trojan et al., 2008). Washed cryosections were incubated with secondary antibodies conjugated to Alexa 488 or Alexa 568 (Invitrogen) in PBS with DAPI (Sigma-Aldrich) to stain the DNA of the cell nuclei, for 1.5 h at room temperature in the dark. After repeated washes with PBS, sections were mounted in MOWIOL 4.88 (Hoechst, Frankfurt, Germany).

### Immunoelectron microscopy

For immunoelectron microscopy a previously established pre-embedding labeling protocol was applied (Sedmak et al., 2009; Sedmak and Wolfrum, 2010). Vibratome sections through mouse retinas were stained by antibodies against ARL2 and visualized by appropriate biotin-labeled secondary anti-mouse IgG antibody combined with a peroxidase-based detection system (Vectastain ABC-Kit, Vector, UK). After fixation with 0.5% OsO_4_, specimens were embedded in Araldite and ultrathin sections analyzed in a Tecnai 12 BioTwin transmission electron microscope (FEI, Eindhoven, The Netherlands) and documented with a charge-coupled device camera (SIS Megaview3; Surface Imaging Systems) as previously described (e.g. (Maerker et al., 2008). Images were processed using Adobe Photoshop CS (Adobe Systems).

### Reproducibility/Statistics

Unless otherwise stated, 100 cells were scored per each replicate, and all experiments were performed in at least triplicate and scored in at least duplicate. Data were processed using Excel and graphed using GraphPad Prism, and error bars shown indicate the standard error of the mean (SEM) for the data set. Individual data points signify the average of technical replicates for each individual cell line. Statistical significance of the difference between individual test groups was assessed using either One-Way or Two-Way ANOVA tests: * = p < 0.05; ** = p < 0.01; *** = p < 0.0001. Technical replicates are considered the replicates of individual lines performed on separate days, while biological replicates are considered the average of the technical replicates for individual lines.

## Acknowledgements

This work was supported by NIH Grants R35GM122568 (RAK), T32GM8367 (RET; ORCID: 0000-0001-5389-4560), F31CA236493-02 (RET), T32NS090650 (EDG; ORCID: 0000-0002-1486-5377), F31NS106755 (EDG), R35GM122549 (TC; ORCID: 0000-0002-6579-7589), and by Foundation Fighting Blindness (FFB) PPA-0717-0719-RAD (UW). We thank colleagues for their generous sharing of key reagents. We give special thanks to Monica Bettencourt-Diaz, Max Nachury, Win Sale, Dorothy Lerit, Maureen Barr, Rytis Prekeris, Tim Stearns, and Peter Jackson for providing input to help us better test and explore models. Emory University Integrated Cellular Imaging (ICI) Microscopy Core and Emory Viral Vector Core of the Emory Neuroscience NINDS Core Facilities Grant 5P30NS055077 provided access to key reagents and instrumentation. We also thank Elisabeth Sehn for her skillful technical assistance at the Facility for Biologic Electron Microscopy, JGU Mainz.

## Author Contributions

Conceptualization, RET, RAK, TC.; Methodology, RET, TC, RAK; Investigation, RET, JL, EDG; Resources, TC, UW, RAK; Writing – Draft Preparation, RET, RAK; Review & Editing, RET, JL, EDF, UW, TC, RAK; Visualization, RET, JL, EDG; Supervision, RAK; Funding Acquisition, RET, RAK, TC, UW.

## Abbreviations used in the text include

FBS: fetal bovine serum
GAP: GTPase activating protein
GEF: guanine nucleotide exchange factor
GFP: green fluorescent protein
GPCR: G-protein coupled receptor
IFT: intraflagellar transport
KO: knockout
MEF: mouse embryonic fibroblast
PCM: pericentriolar material
PFA: paraformaldehyde
SHH: Sonic Hedgehog
Smo: Smoothened
TZ: transition zone
WT: wild-type

---

**Figure S1:** *Loss of ELMOD2 leads to disrupted ciliary morphology and protein localization.* **(A)** Representative image of branching in ELMOD2 KO cilia. SIM images were collected at 100x magnification using cells stained for ARL13B. Scale = 2μm. **(B)** Centrin staining in cilia is increased in both WT and ELMOD2 KO cells after treatment with ciliobrevin. Representative images collected via widefield microscopy (100x magnification) are shown. WT and KO cells were serum starved for 24 hours before being treated with either DMSO or 30μM ciliobrevin for 1hr at 37°C. Insets highlight individual cilia and the presence or absence of centrin.

**Figure S2:** *SSTR3 and GPR161 localization is decreased/lost in ELMOD2 KO cilia.* **(A)** WT or ELMOD2 KO cells were transfected with plasmid directing expression of SSTR3-GFP and the next day were serum starved. KO cells displayed strongly reduced ciliary GFP. Cells were fixed with 4% PFA, permeabilized with 0.1% Triton X-100, and co-stained with ARL13B. Representative images were collected using widefield microscopy (100x magnification). Scale bar = 2 μm. **(B)** ELMOD2 KO cells show decreased recruitment of (endogenous) SSTR3 and GPR161. Serum-starved cells were fixed and stained using protocols required for detecting the appropriate antigen, as described under Materials and Methods. Representative images were collected via widefield microscopy at 100x magnification. Samples were co-stained with either acetylated tubulin or ARL13B to mark cilia. Scale bar = 10 μm. **(C)** Cells were scored for endogenous SSTR3 staining (which is naturally faint, so were either binned as existent or non-existent). The experiment was performed in duplicate, and the average of the duplicates of individual lines are shown here as individual points of interleaved scatterplot. Error bars represent SEM, and statistical significance was assessed via One-Way ANOVA. ***=p<0.0001.

**Figure S3:** *ELMOD2 localizes to cilia in WT MEFs upon ciliobrevin treatment.* WT MEFs treated either with 0.6% DMSO (top) or 30μM ciliobrevin (bottom) for 1hr were fixed with 4% PFA, permeabilized with 0.1% Triton X-100, and stained for ELMOD2 and ARL13B. Only upon blocking of ciliary retrograde transport via ciliobrevin do we observe ELMOD2 localization to cilia. Images were collected via widefield imaging at 100x magnification.

**Figure S4:** *ELMOD2 does not localize to non-centrosomal rootlets.* ELMOD2 specifically localizes to centrosome-associated rootlets rather than all Rootletin staining. Serum-starved, WT MEFs were fixed with ice-cold methanol and stained for ELMOD2, acetylated tubulin, and Rootletin. Widefield images were collected at 100x magnification. Scale = 10μm.

**Figure S5:** *Live cell imaging of GFP-Rootletin expressing MEFs reveal that serum starvation induces rootlet tendrils to dissociate from the centrosome.* Cells were imaged every 5 minutes over a 1-hour imaging window using widefield microscopy, 20x magnification. Cells were maintained at 37°C and 5% CO_2_ and imaged for 1 hour without (top panels) or with (bottom panels) serum starvation. While no rootlet release was evident without serum starvation, but after serum starvation rootlet release was evident within minutes.

**Figure S6:** *Summary of Rootletin alleles generated by CRISPR/Cas9.* (**A**) We designed 4 guides to use in CRISPR/Cas9 genome editing and the sites they target are shown above the targeted exons. The mouse *Crocc* gene encodes 37 exons, with the open reading frame shown below the spliced exons. (**B**) A total of 5 Rootletin KO lines were generated, along with the Rootletin^Δ239^ line (G1, #21). Two clones were generated using guide 1, two others from guide 2, and 1 from guide 4. Rootletin^Δ239^ was generated from guide 1. Genomic DNA sequencing was performed on genomic DNA surrounding each targeted region and the indels are listed under Alleles, along with the resulting protein sequences. The black font indicates WT protein sequence while the red font indicates nonsense protein sequence resulting from a frame shift and an asterisk indicates a stop codon. Each of the knockout clones led to frameshifting mutations which were predicted to generate nonfunctional protein products, as later confirmed by Western blot. In contrast, the line termed Rootletin^Δ239^ displays very strong staining of Rootletin, despite having both alleles frameshifted at the targeted site. The use of a downstream methionine to initiate protein translation is proposed as an explanation of the shorter protein product seen in immunoblots.

**Figure S7:** *ELMOD2 KO cells show no change in Rootletin protein expression.* (**A**) Cell pellets from confluent wells of a 6-well plate were thawed and immediately immersed in 1x sample buffer + BME, as in our hands rootletin is unstable after cell lysis. Equal volumes of protein were loaded into a 7.5% acrylamide gel and transferred onto nitrocellulose membrane. Membranes were blotted for chicken-anti-rootletin (1:1000 dilution) to check for Rootletin expression in WT versus ELMOD2 KO cells. (**B**) Ponceau staining of the membrane was performed to check for equal protein loading.

**Figure S8:** *Centrin and ARL2 localize to Rootletin KO cilia.* Serum-starved Rootletin KO cells have increased recruitment of both ARL2 and centrin. Cells were stained for ARL13B as a marker of cilia and either centrin or ARL2. Wide-field mages were collected at 100x magnification. Scale = 10μm.

**Figure S9:** *Western blotting confirms the loss of Rootletin in KO lines and increased expression of an N-terminal truncation in Rootletin^Δ239^ MEFs.* Raw data of the Western shown in Figure 5A are shown, including (**A**) Ponceau S staining of the nitrocellulose membrane to confirm equal protein loading, and (**B-C**) uncropped images of the films collected at 1 min and 3 min exposures respectively. Membranes were stained with chicken-anti-Rootletin at 1:1000 dilution in 5% Blotto. (**D-F**) Western blot and Ponceau of WT, WT + myc-Rootletin^Δ239^, *Rootletin*^Δ239^ cells, and *Rootletin*^Δ239^ + myc-Rootletin^Δ239^, using the same conditions as described for (A).

**Figure S10:** *ELMOD2 still localizes to mitochondria, Flemming bodies, and centrosomes in Rootletin KO cells.* To test whether the deletion of Rootletin alters ELMOD2 staining at sites other than rootlets, we used a number of fixation conditions to stain Rootletin KO cells for ELMOD2. Loss of Rootletin does not alter ELMOD2 staining at mitochondria (**A**), at Flemming bodies (**B**), or at metaphase centrosomes (**C**). Cells were fixed and stained for ELMOD2 and γ-tubulin (to mark both midbodies and centrosomes). Widefield images at 100x magnification are shown. Scale = 10μm.

**Figure S11:** *ARL3 localizes to cilia and centrosomes but not rootlets in WT MEFs.* Representative widefield images (100x magnification) of ARL3 localization in WT MEFs are shown. With 4% PFA fixation, ARL3 staining at cilia is observed, as seen by co-staining with γ-tubulin and acetylated tubulin. With ice-cold methanol fixation, centrosomal staining of ARL3 is evident, but it does not extend to rootlets (using conditions in which one can readily detect ELMOD2 and ARL2 at rootlets). Scale = 10μm.

**Figure S12:** *ELMOD2 KO does not alter the localization of IFT or transition zone (TZ) markers.* Cells were fixed and stained for either markers of transition zone (**A-B**) or intraflagellar transport (**C**) to determine if there are overt defects in these compartments in ELMOD2 KO cells. For transition zone, cells were fixed for 10 min with ice-cold methanol, blocked with 10% FBS, and stained for either NPHP4 or Cep290, along with a ciliary marker (ARL13B or acetylated tubulin). To look at IFT, cells were fixed with 4% PFA, permeabilized with 0.1% Triton X-100, and stained for IFT88. Representative images were collected via widefield microscopy at 100x magnification. Scale = 10μm.

**Figure S13:** *Representative images demonstrating that ELMOD2 KO leads to misregulation of specific markers of ciliogenesis.* Widefield images were collected at 100x magnification of (**A**) Cep164, (**B**) TTBK2, and (**C**) CP110 staining at centrosomes (γ-tubulin) in WT versus ELMOD2 KO cells. Scale = 10 μm.

**Figure S14:** *Loss of Rootletin has no obvious effects on cell cycle of unsynchronized cycling cells.* WT, ELMOD2 KO, and Rootletin KO cells were harvested, fixed, and stained for propidium iodide staining to check the DNA content of cells via flow cytometry. 10,000 events were counted per cell line, and these experiments were performed in duplicate for each of the lines shown. Data were processed and plotted via FlowJo software, ensuring to use the same gating parameters for each sample. Representative graphs are shown here.

**Figure S15:** *Loss of either ELMOD2 or Rootletin does not affect ARL3 or centrin expression in MEFs.* Equal protein from whole cell lysates was loaded onto 15% acrylamide gels and transferred onto nitrocellulose membrane. Membranes were blotted for either mouse-anti-centrin (1:500 dilution) (**A**) or rabbit-anti-ARL3 (1:500 dilution) (**B**) to assess changes in protein expression that may be a product of increases in ploidy. Ponceau staining of each respective membrane (**C-D**) was performed to check for equal loading.

**Figure S16:** *TTBK2 is increased at centrosomes of ELMOD2 KO and Rootletin KO mononucleated cells having no more than two centrosomes.* Cells were scored as described in Figure 8B, but with scoring only mononucleated cells with 1-2 centrosomes. This was a control to ensure that increased ciliogenesis is not simply a side effect of cells with obvious cell cycle defects. Data are shown as a stacked bar graph, representing the average of duplicates of multiple cell lines. Error bars indicate SEM.

**Figure S17:** *ARL2 localizes along the length of cilia in human (multiciliated) bronchial epithelial cells, while ELMOD2 localizes to the tips of cilia and rootlets.* Primary cultures of human bronchial cells were grown on transwell plates before being fixed with ice-cold methanol and stained for Rootletin, acetylated tubulin and either (**A**) ARL2 or (**B**) ELMOD2. Confocal images were collected at 100x magnification, and z-projections were generated. Representative images of fields of bronchial cells are shown on the left. On the right, insets that highlight cells in which one can readily distinguish cilia from plasma membrane from rootlets. ARL2 is found almost exclusively at cilia in these cells, while ELMOD2 stains both the tip of cilia and (more faintly) the rootlets.

