## Supplemental Figures for "“Roles for ELMOD2 and Rootletin in Ciliogenesis”"

### Figure S1

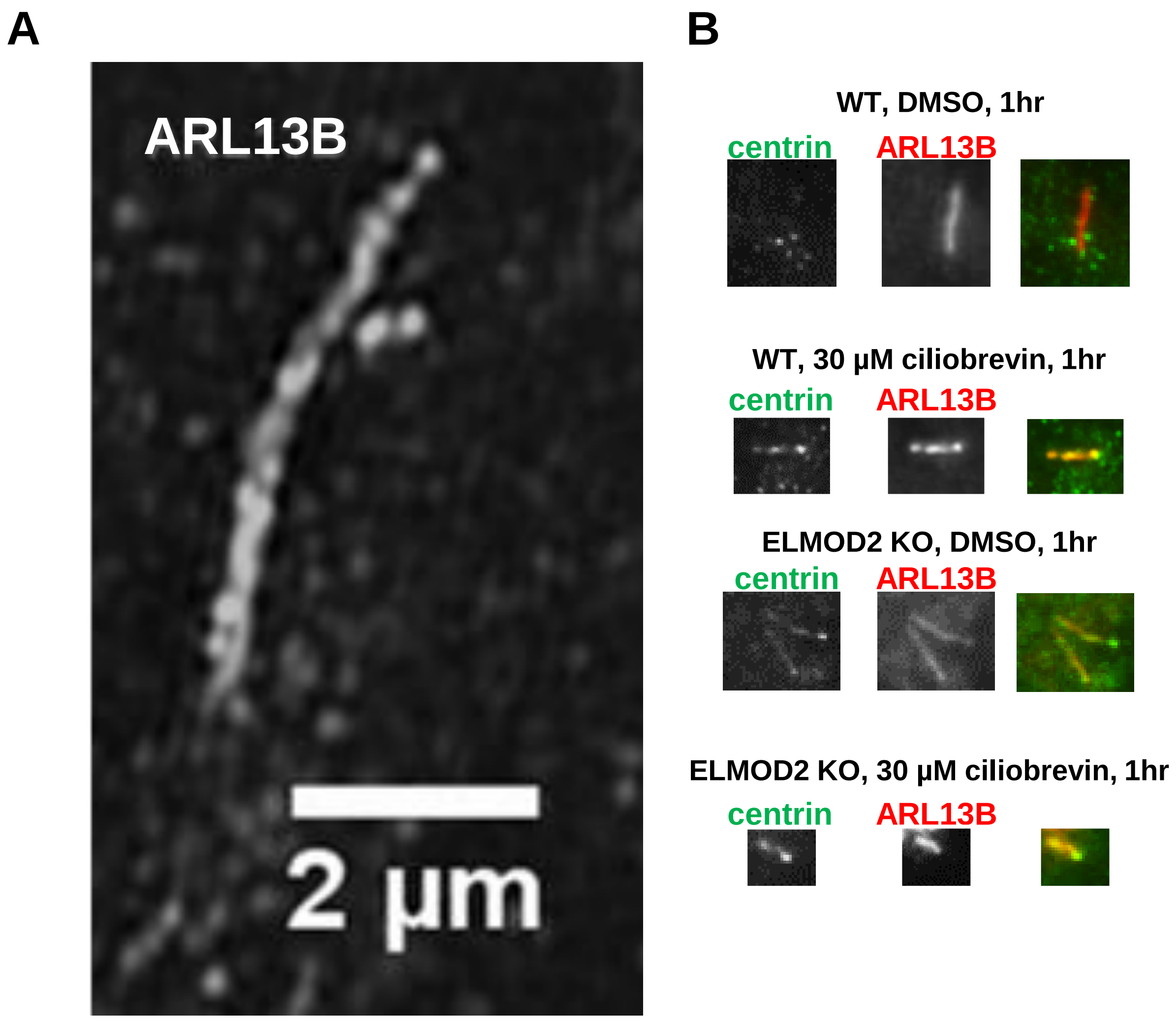
A
ARL13B
2 µm
B
WT, DMSO, 1hr
centrin
ARL13B
WT, 30 µM ciliobrevin, 1hr
centrin
ARL13B
ELMOD2 KO, DMSO, 1hr
centrin
ARL13B
ELMOD2 KO, 30 µM ciliobrevin, 1hr
centrin
ARL13B

### Figure S2

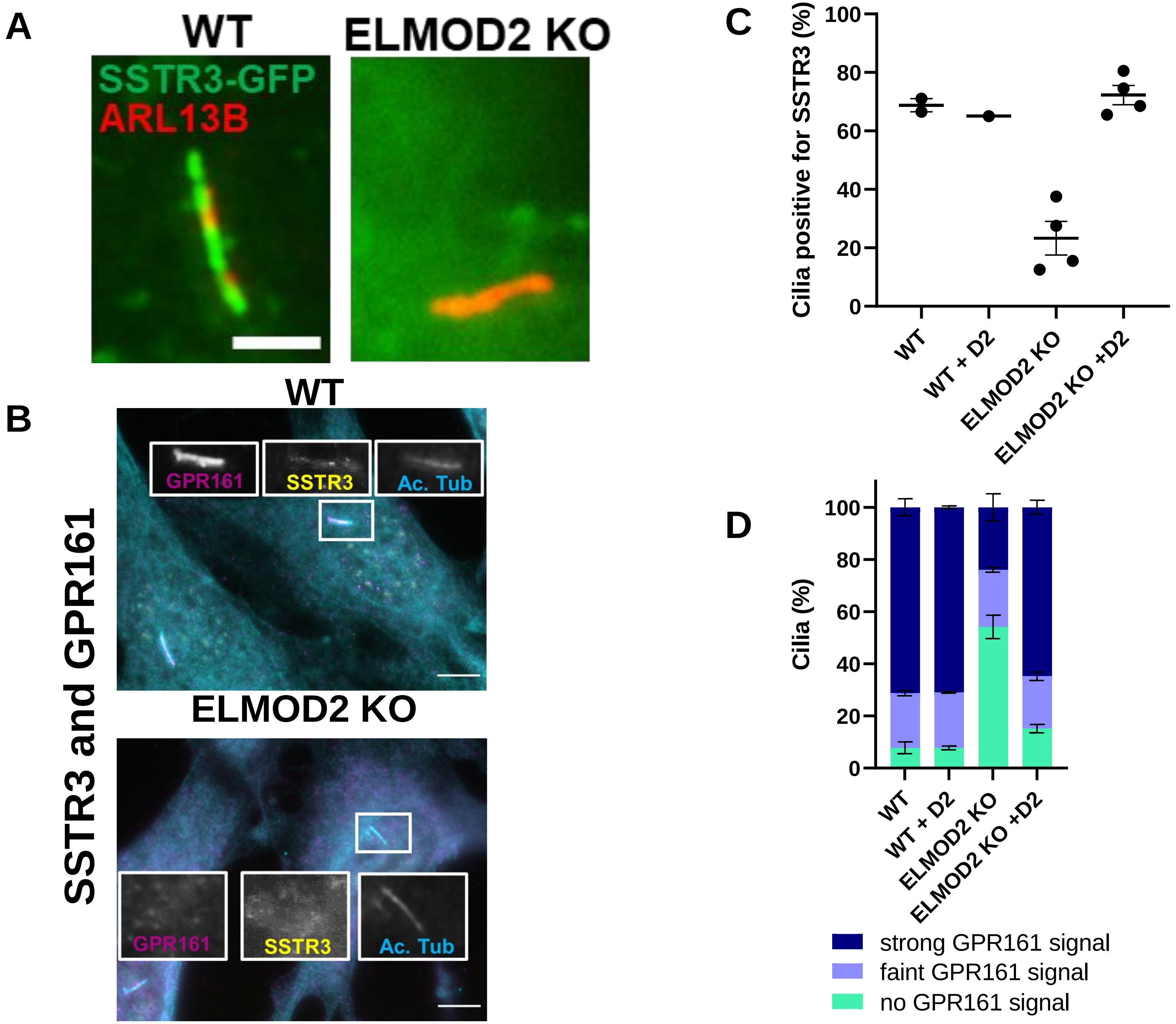
A
WT
ELMOD2 KO
SSTR3-GFP
ARL13B
B
WT
ELMOD2 KO
SSTR3 and GPR161
GPR161
SSTR3
Ac. Tub
C
Cilia positive for SSTR3 (%)
100
80
60
40
20
0
WT
WT + D2
ELMOD2 KO
ELMOD2 KO +D2
D
Cilia (%)
100
80
60
40
20
0
WT
WT + D2
ELMOD2 KO
ELMOD2 KO +D2
strong GPR161 signal
faint GPR161 signal
no GPR161 signal

### Figure S3

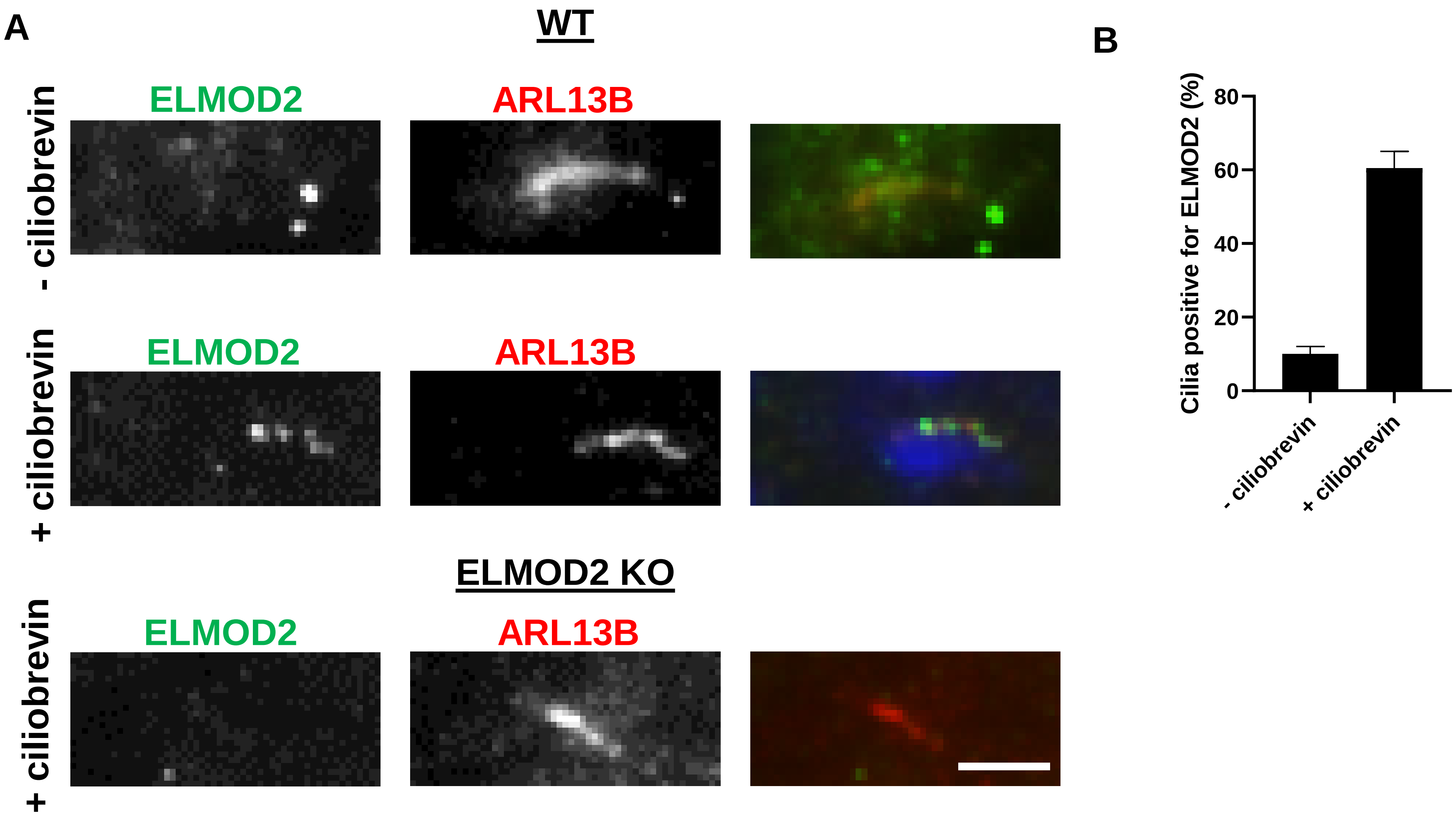
A
WT
- ciliobrevin
ELMOD2
ARL13B
+ ciliobrevin
ELMOD2
ARL13B
ELMOD2 KO
+ ciliobrevin
ELMOD2
ARL13B
B
Cilia positive for ELMOD2 (%)
80
60
40
20
0
- ciliobrevin
+ ciliobrevin

**Figure S4**

**ELMOD2 localizes to centrosomal rootlets in WT MEFs**

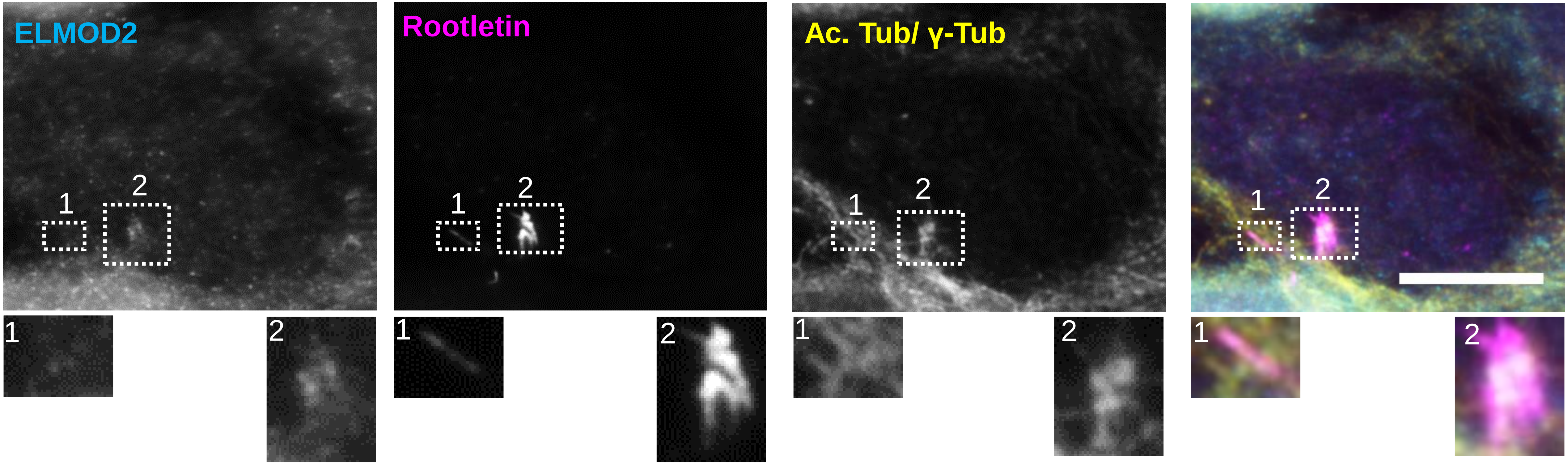

**Figure S5**

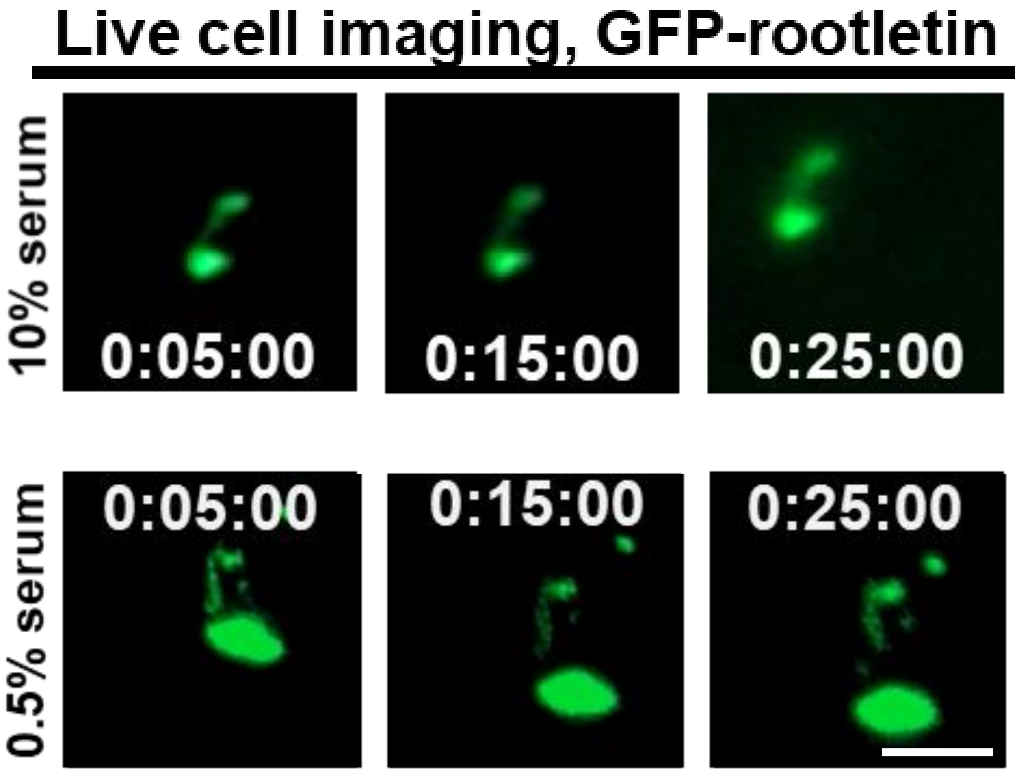
Live cell imaging, GFP-rootletin
10% serum
0:05:00
0:15:00
0:25:00
0.5% serum
0:05:00
0:15:00
0:25:00

### Figure S6

A

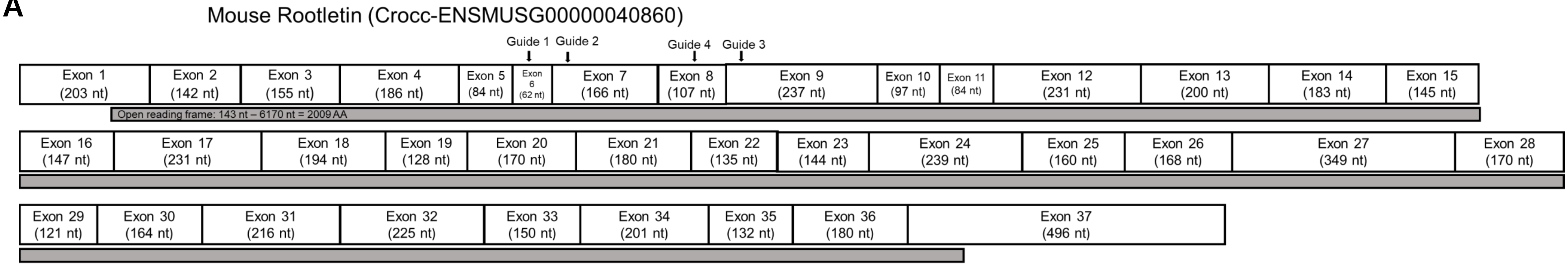


B

| Clone ID | Guide | Alleles |
|---|---|---|
| G1, #12 | 1 | Both alleles: 1 bp deletion (C); **AA sequence:** …TEHSQDLDSALLR* |
| G1, #31 | 1 | Both alleles: 1 bp insertion (C); **AA sequence:** …TEHSQDLDSALLRPRGGTAEVRGWDSPWAPSG* |
| G2, #17 | 2 | Both alleles: 1 bp insertion (C); **AA sequence:** …VSKCPLNTPTPPRSASLAQVNAMLREQLDQANLANQALSEDIPQGDQ* |
| G2, #20 | 2 | Both alleles: 4bp deletion (TACG); **AA sequence:** …VSKCPLNTPTPPRSASLAQVNAMLREQLDQANLANQALSEDTR* |
| G4, #2 | 4 | Both alleles: 1 bp insertion (A) ; **AA sequence:** …SFNAYFSSEHSRLLRLWRQVMGLRQAGQRGEDGHGEVRLLGAQGASCPG* |
| Rootletin$^{\Delta 239}$ | 1 | 1: 1 bp insertion (A); **AA sequence: …**TEHSQDLDSALLRHRGGTAEVRGWDSPWAPSG*<br>2: 2 bp insertion (CT); **AA sequence: …** TEHSQDLDSALLRP* |

**Figure S7**

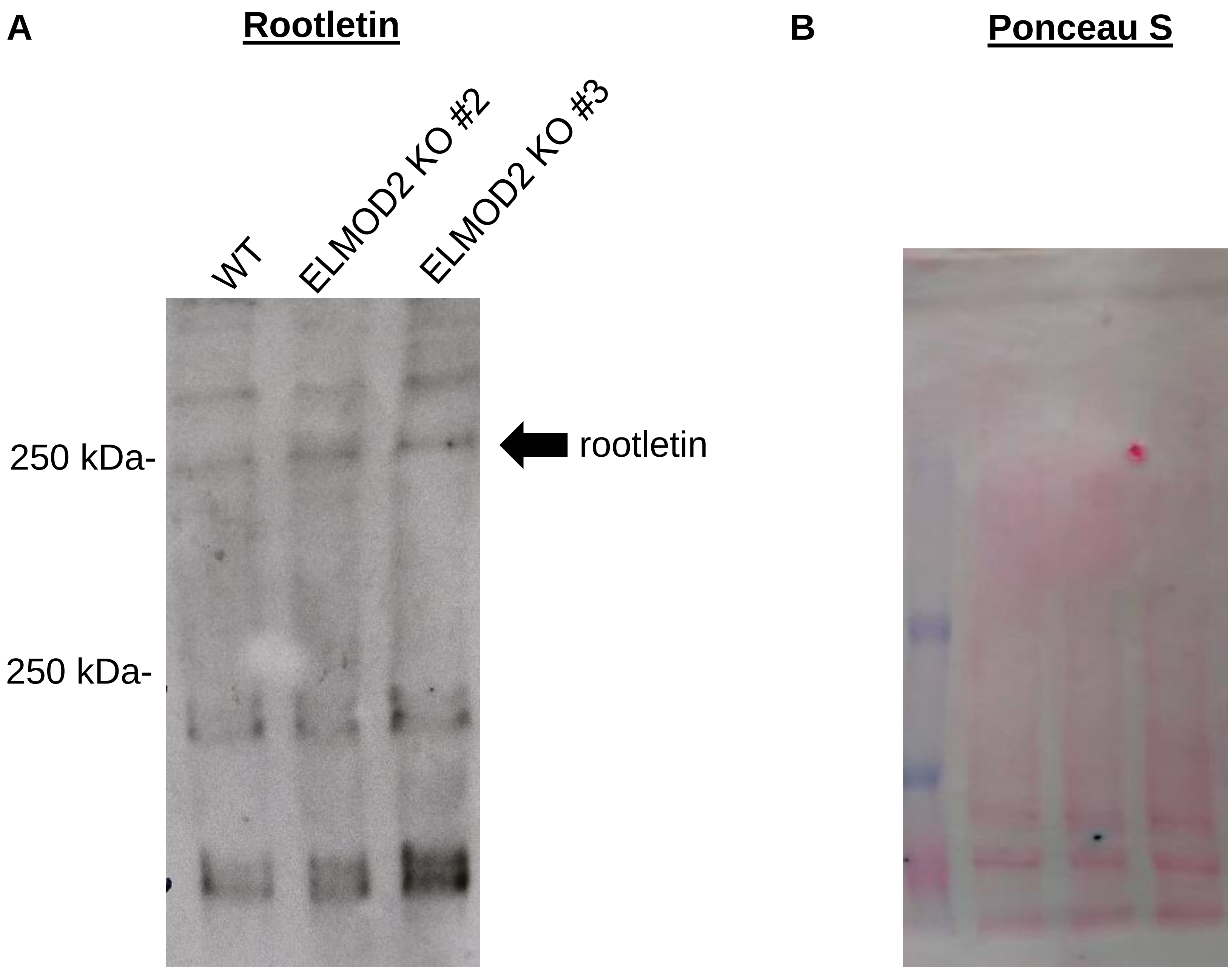
A
Rootletin
WT
ELMOD2 KO #2
ELMOD2 KO #3
250 kDa-
250 kDa-
rootletin
B
Ponceau S

**Figure S8**

#### Centrin inside Rootletin KO Cilia

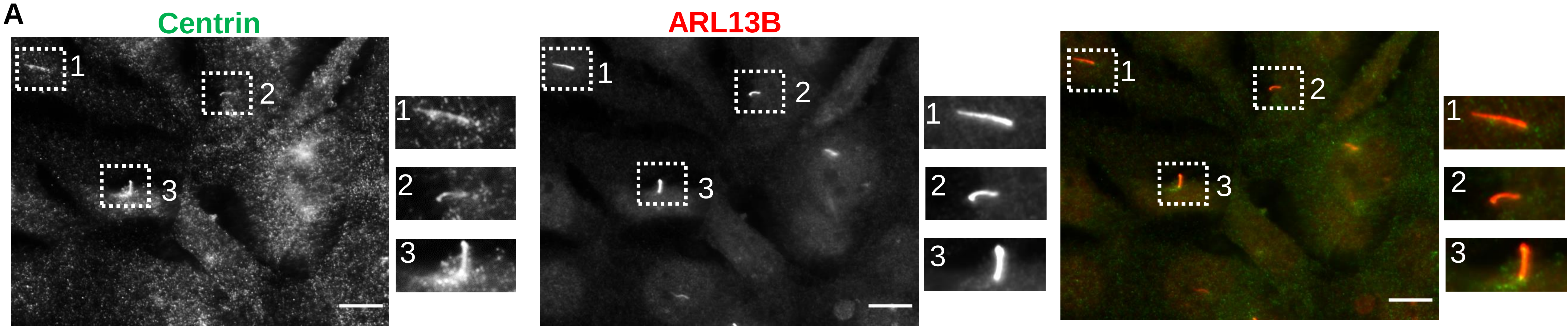


#### ARL2 inside Rootletin KO Cilia

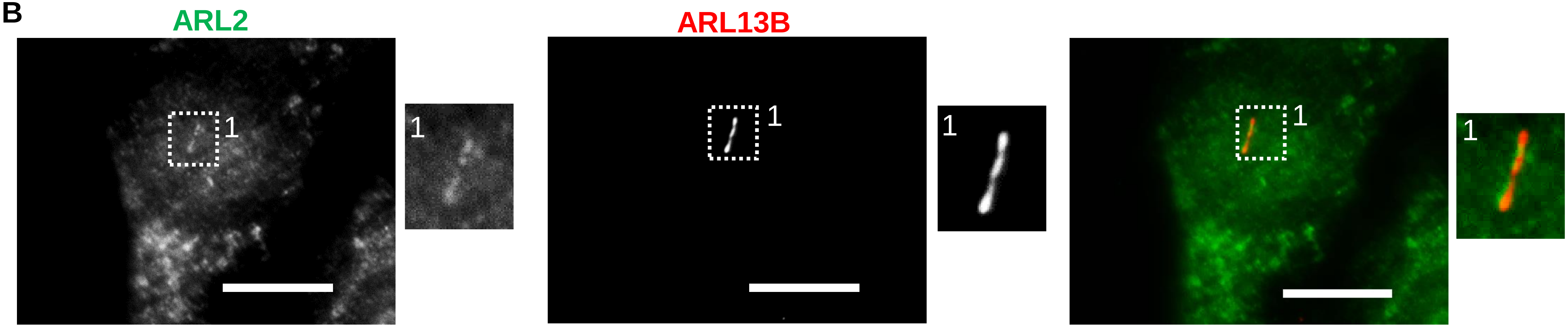

### Figure S9

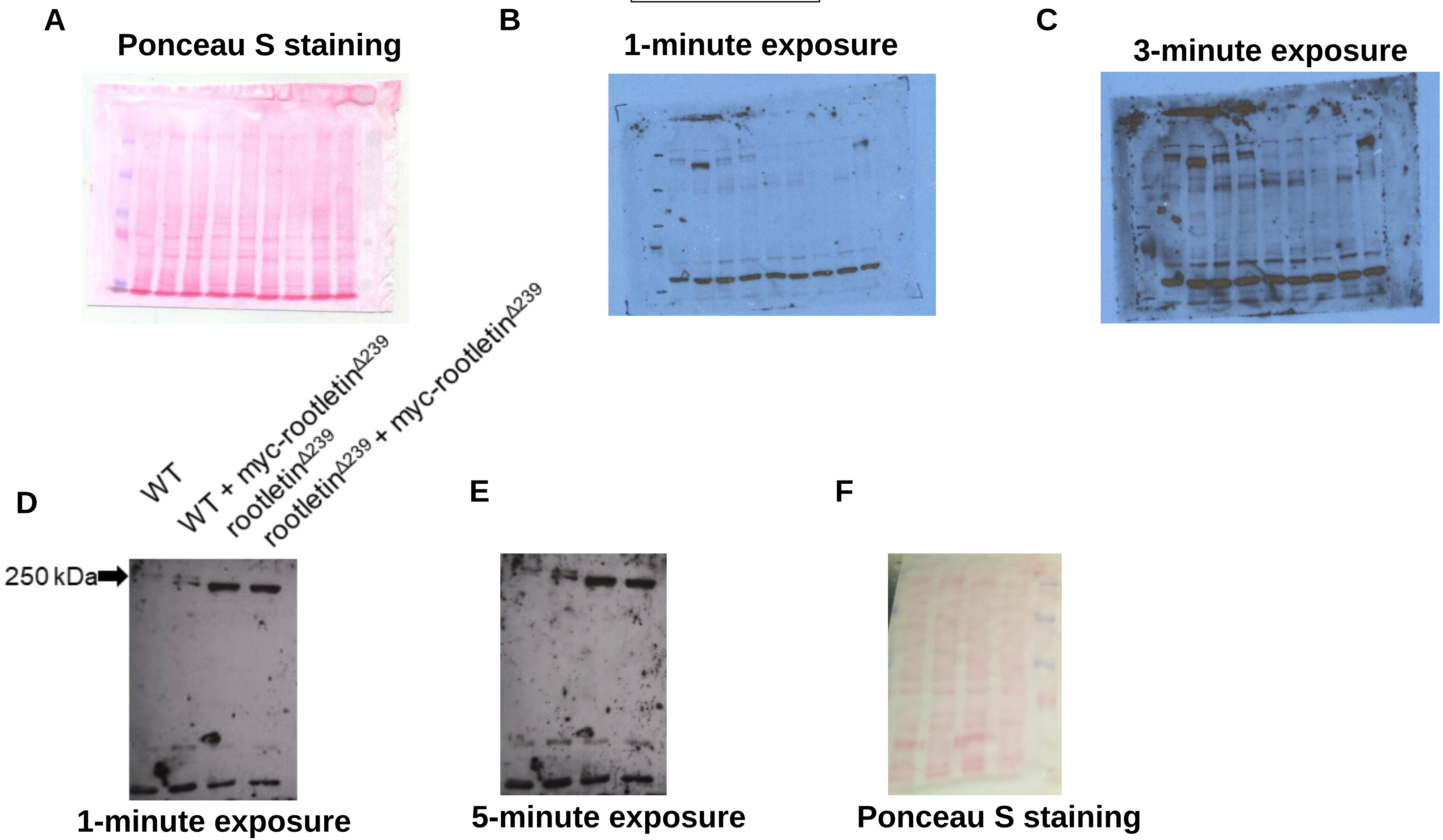
A
Ponceau S staining
B
1-minute exposure
C
3-minute exposure
D
WT
WT + myc-rootletinΔ239
rootletinΔ239
rootletinΔ239 + myc-rootletinΔ239
250 kDa
1-minute exposure
E
5-minute exposure
F
Ponceau S staining

Figure S10

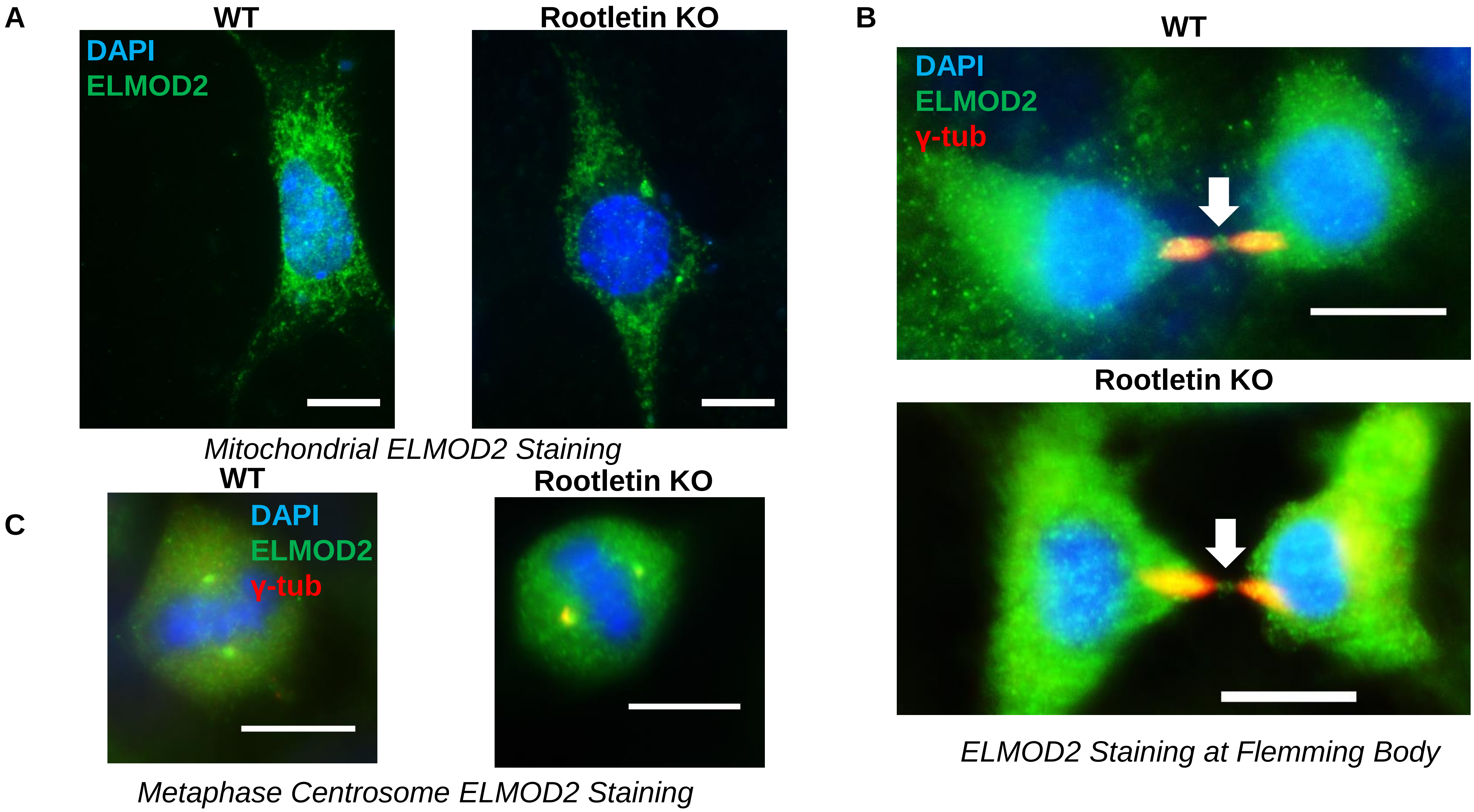
A
WT
DAPI
ELMOD2
Rootletin KO
Mitochondrial ELMOD2 Staining
B
WT
DAPI
ELMOD2
γ-tub
Rootletin KO
ELMOD2 Staining at Flemming Body
C
WT
DAPI
ELMOD2
γ-tub
Rootletin KO
Metaphase Centrosome ELMOD2 Staining

### Figure S11

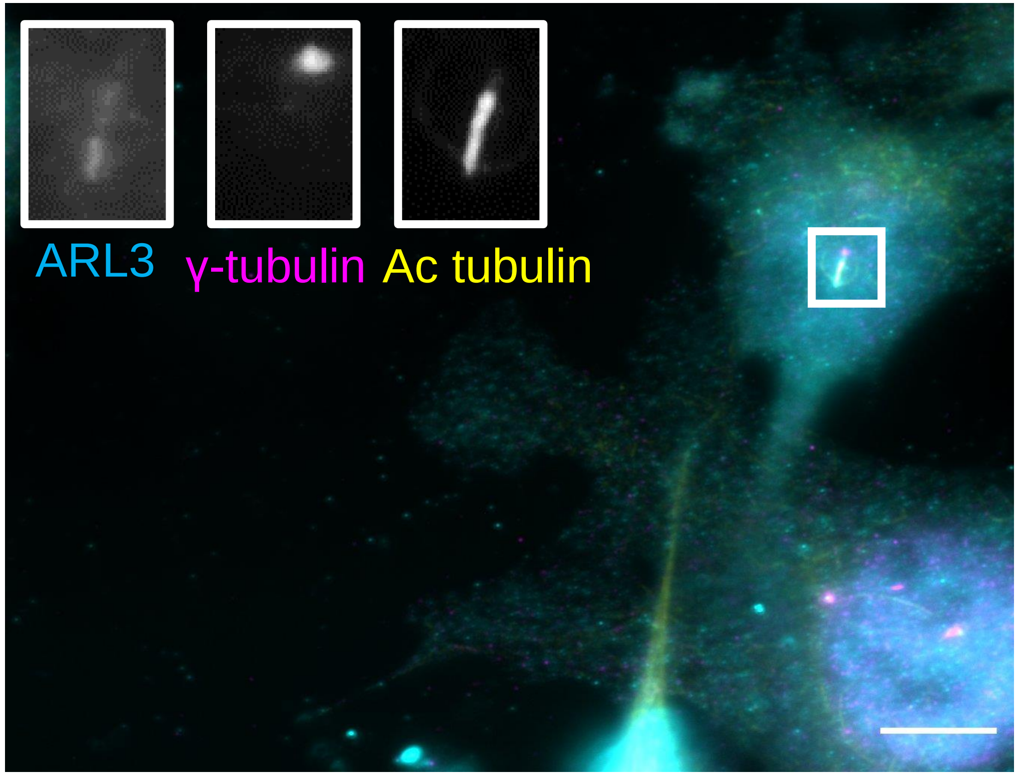

4% PFA

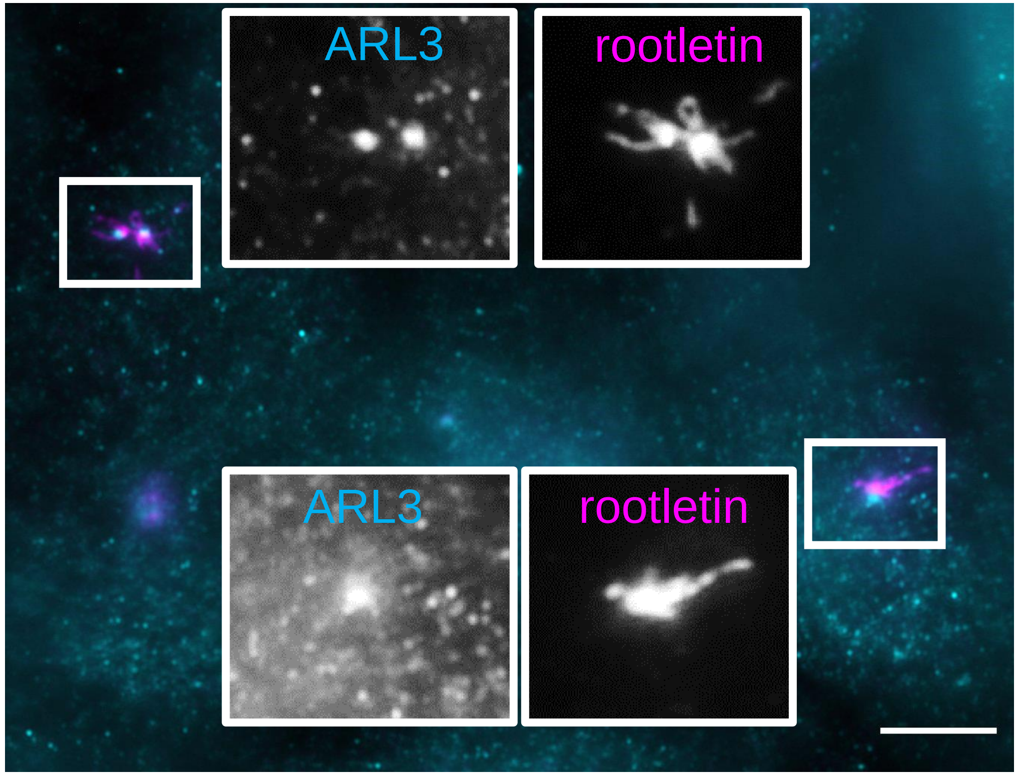

5 min ice cold methanol

### Figure S12

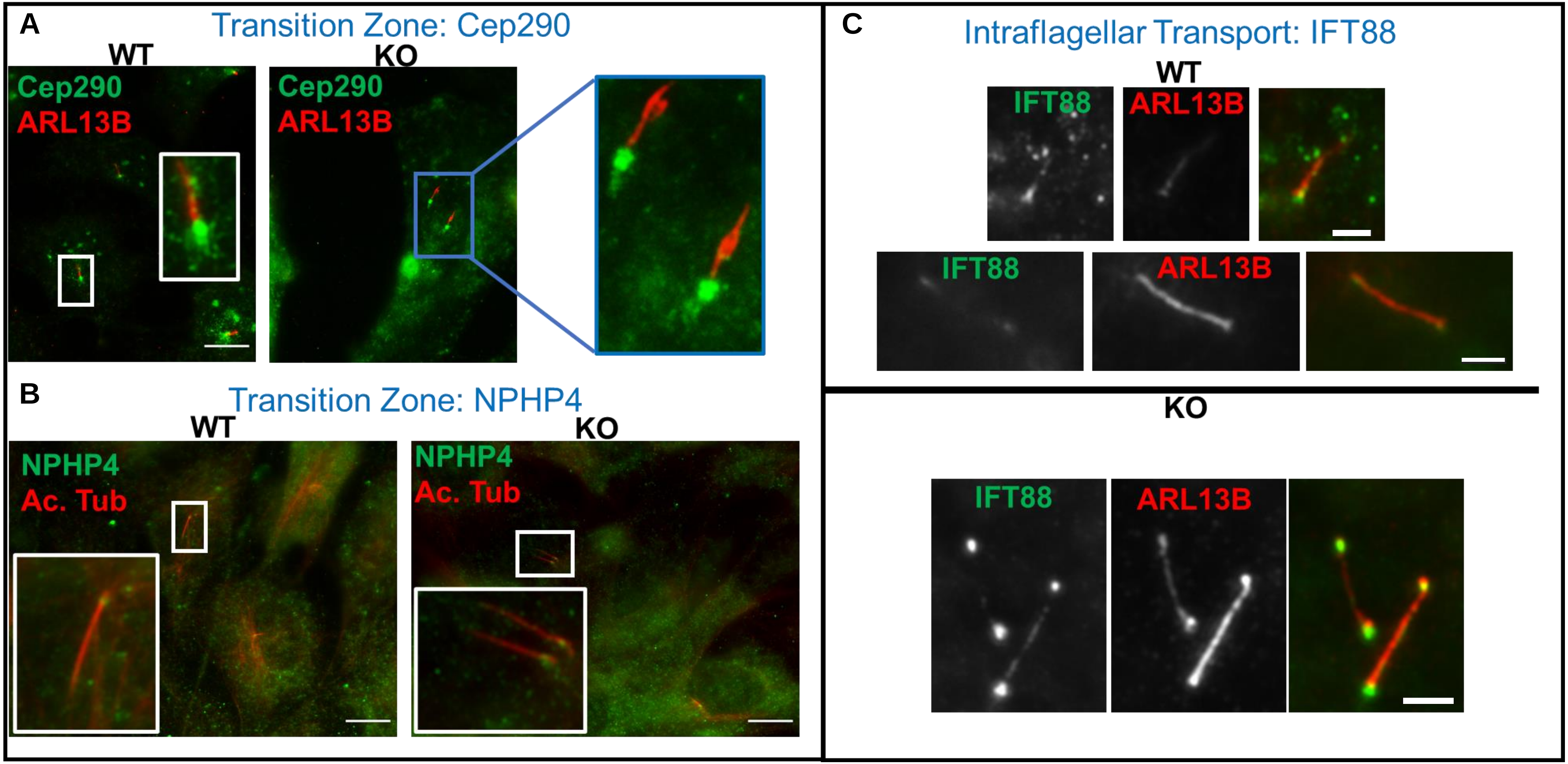
A
Transition Zone: Cep290
WT
KO
Cep290
ARL13B
Cep290
ARL13B
B
Transition Zone: NPHP4
WT
KO
NPHP4
Ac. Tub
NPHP4
Ac. Tub
C
Intraflagellar Transport: IFT88
WT
IFT88
ARL13B
IFT88
ARL13B
KO
IFT88
ARL13B

### Figure S13

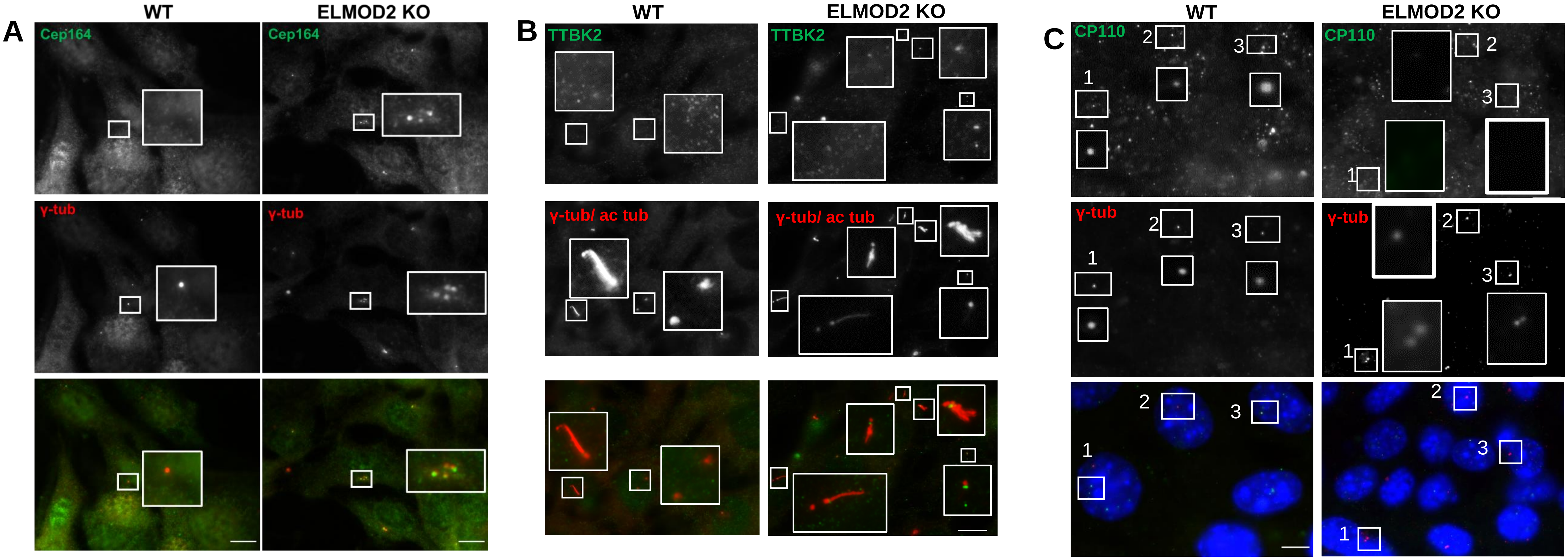

A
WT
ELMOD2 KO
Cep164
Cep164
γ-tub
γ-tub
B
WT
ELMOD2 KO
TTBK2
TTBK2
γ-tub/ ac tub
γ-tub/ ac tub
C
WT
ELMOD2 KO
CP110
CP110
γ-tub
γ-tub
1
2
3

### Figure S14

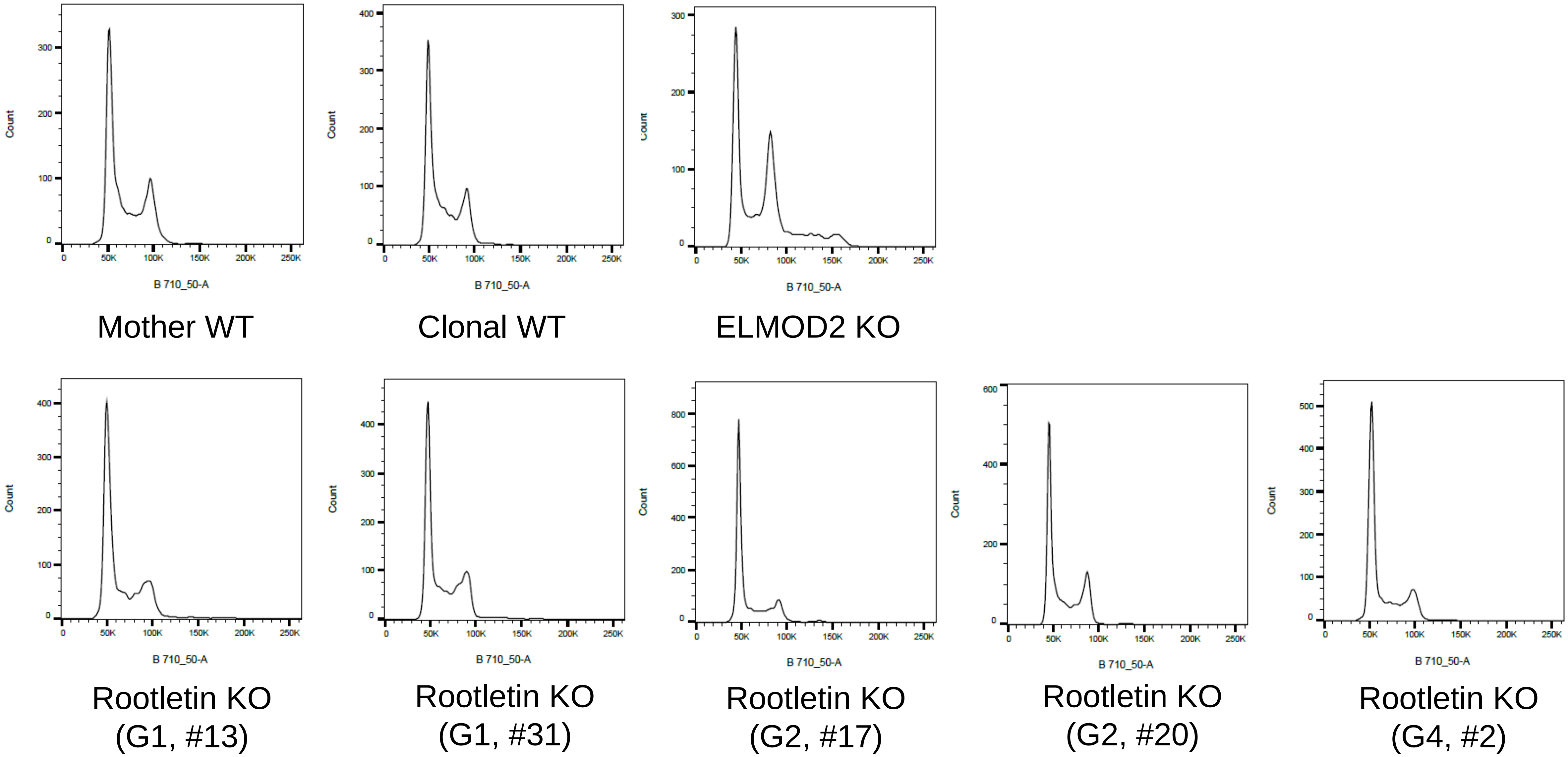

Count
B 710_50-A
Mother WT
Clonal WT
ELMOD2 KO
Rootletin KO (G1, #13)
Rootletin KO (G1, #31)
Rootletin KO (G2, #17)
Rootletin KO (G2, #20)
Rootletin KO (G4, #2)

**Figure S15**

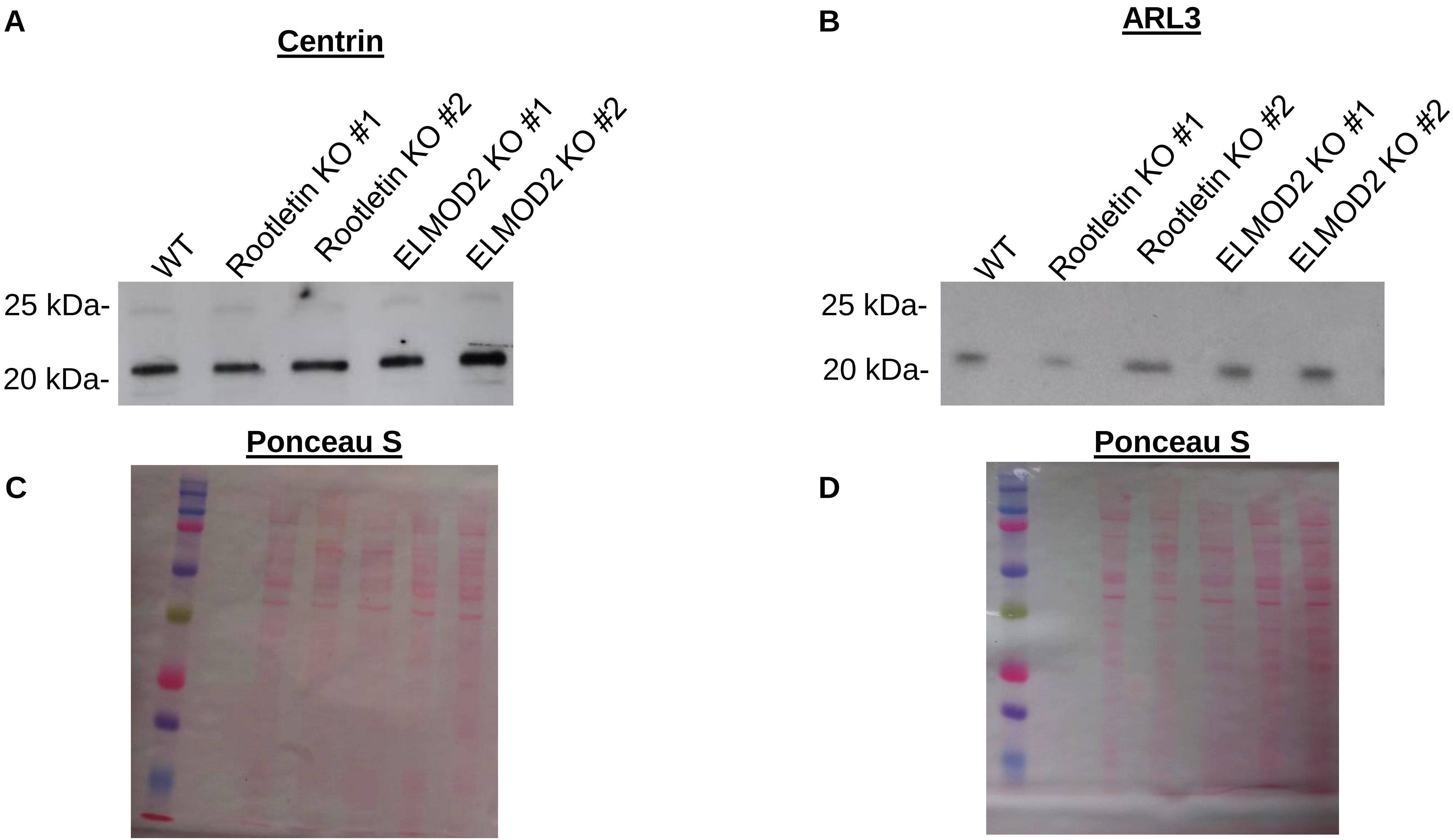
A
Centrin
WT
Rootletin KO #1
Rootletin KO #2
ELMOD2 KO #1
ELMOD2 KO #2
25 kDa-
20 kDa-
B
ARL3
WT
Rootletin KO #1
Rootletin KO #2
ELMOD2 KO #1
ELMOD2 KO #2
25 kDa-
20 kDa-
Ponceau S
C
Ponceau S
D

### Figure S16

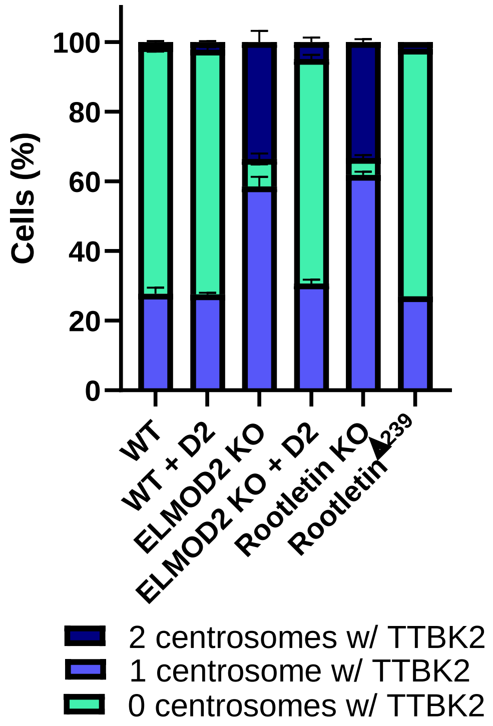

Cells (%)
0
20
40
60
80
100
WT
WT + D2
ELMOD2 KO
ELMOD2 KO + D2
Rootletin KO
Rootletin Δ239
2 centrosomes w/ TTBK2
1 centrosome w/ TTBK2
0 centrosomes w/ TTBK2

**Figure S17**

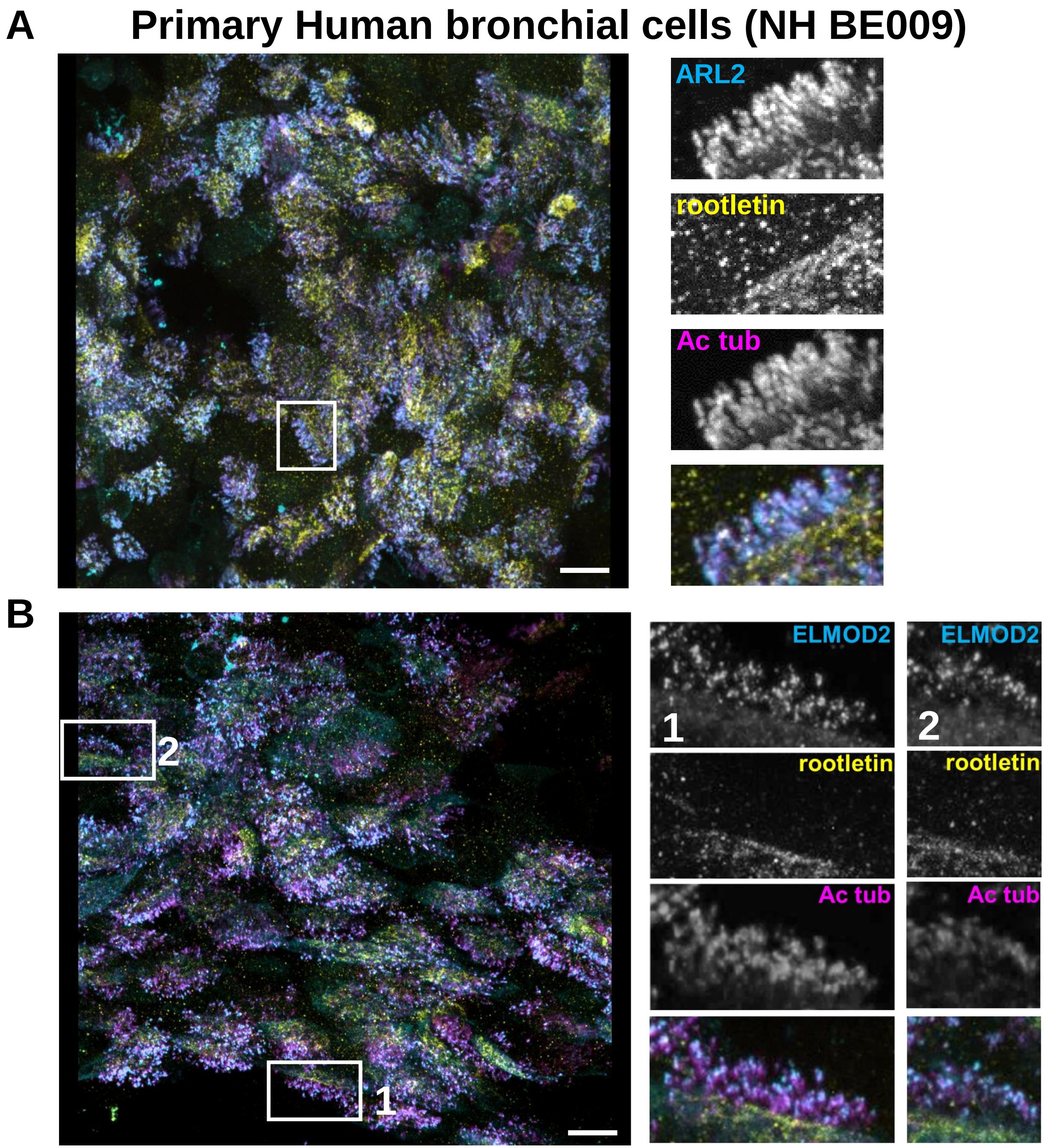
A
Primary Human bronchial cells (NH BE009)
ARL2
rootletin
Ac tub
B
2
1
ELMOD2
ELMOD2
1
2
rootletin
rootletin
Ac tub
Ac tub
